# Identification of residues potentially involved in optical shifts in the water-soluble chlorophyll-a binding protein through molecular dynamics simulations

**DOI:** 10.1101/2023.10.11.561876

**Authors:** Martina Mai, Valter Zazubovich, R. A. Mansbach

## Abstract

Reversible light- and thermally-induced spectral shifts are universally observed in a wide variety of pigment-protein complexes, at temperatures ranging from cryogenic to ambient. They can be observed either directly, in single-molecule spectroscopy experiments, or via non-photochemical spectral hole burning. These shifts are important to understand, for example, to gain a clearer picture of the primary processes of photosynthesis, or of general features of the protein energy landscapes. In this article, we have employed large-scale molecular dynamics simulations of a prototypical pigment-protein complex to better understand these shifts at a molecular scale. Although multiple mechanisms have been proposed over the years, no verification of these proposals via MD simulations has thus far been performed; our work represents the first step in this direction. The common requirement for all these mechanisms is the presence of double-well (or multiple-well) features of the protein energy landscapes. In this work, from large-scale molecular dynamics simulations of the Water-Soluble Chlorophyll-binding Protein complex, we identified side chain rotations of certain amino acid residues as likely candidates for relevant multi-well landscape features. The protein free energy landscapes associated with side chain rotations feature energy barriers of around 1100- 1600 cm*^−^*^1^, in agreement with optical spectroscopy results, with the most promising residue type associated with experimental signatures being serine, which possesses a symmetric landscape and moment of inertia of a relevant magnitude.

## 1 Introduction

Over the years, high-resolution optical spectroscopy methods such as single-molecule spectroscopy and non-photochemical spectral hole burning (NPHB) have been successfully employed to study various non-crystalline systems including glasses, polymers, and proteins, and, in particular, pigment-protein complexes involved in photosynthesis.^1–18^ References 1-9 and 10-18 represent just a small select sample of works employing NPHB and single complex spectroscopy, respectively, to photosynthesis-related research. The very existence of NPHB and of light- and thermally-induced line shifts in the single-complex optical spectra indicates the occurrence of small structural changes in the pigment environment, even at cryogenic temperatures. In other words, these complexes can exist in multiple slightly different structural states, represented as minima of the free energy landscape, even when the protein is properly folded. At cryogenic temperatures, as spectral lines (zero-phonon lines) of the pigment get narrower, spectral shifts become more easily observable, rendering them highly sensitive to local protein dynamics. NPHB and line shifts in single complex spectra have been observed in all pigment-protein complexes studied by these techniques so far.^1–6,8,9,11–18^ It is not clear if the presence of such spectral shifts is useful from an evolutionary perspective, or if it is merely an inevitable consequence of the amorphous character of proteins. While it is believed that some light-induced structural changes are involved in non-photochemical quenching (NPQ), a mechanism protecting pigment-protein complexes in excessive illumination conditions^19^, small structural changes manifesting via NPHB and shifts of the single-molecule spectral lines are a more universal phenomenon than NPQ.

The minimal model capable of explaining results of NPHB and single-complex experiments of pigment-protein complexes is the Two-Level System (TLS) model, in which the free energy landscape (FEL) of a protein near its native state is assumed to be well-represented by a double-well potential along some generalized coordinate. It was first developed to explain anomalous low-temperature properties in glasses^20,21^ and was later applied to NPHB, see Jankowiak et al. ^1^ for more details. (We do not suggest that exactly the same TLS are responsible for both low-temperature thermodynamic anomalies and spectral shifts, but merely that TLS/MLS are currently widely accepted as good qualitative explanations for both types of phenomena.) The TLS model implies the existence of two conformational substates of similar free energy corresponding to slightly different optical transition frequencies of the pigment embedded in the protein. One may also generalize this model to account for the possibility of more than two sub-states of similar free energy in the near-native-state FEL, in which case one refers to it as a Multi-Level System (MLS) model. In either case, the relevant structural changes at cryogenic temperatures occur (with reasonable probability) on a timescale of nanoseconds when the pigment is in its excited electronic state and on a timescale of hours or even days when the pigment is in its ground electronic state (cf. Fig. 1). Analysis of hole burning and hole recovery data in NPHB and spectral shifts in single-complex experiments allows for making some estimates of the heights of the energy barriers between the wells (on the order of 1000 cm*^−^*^1^ for the ground state), the sizes of the entities involved in these small structural changes and the magnitudes of their movements (*md*^2^ ∼ 1.4-4.0×10*^−^*^46^ kg-m^2^).^3–6^ Several specific mechanisms, including TLS/MLS resulting from single hydrogen bonds, cooperative rearrangements of hydrogen bonding networks, and movements of light protein side-groups, such as hydroxyl and methyl groups, have been proposed to explain these spectral shifts qualitatively.^3,5,6,8,9,22^ However, these proposals have not been independently verified and the precise nature of the atomic / molecular entities involved remains largely unknown.

**Figure 1:**
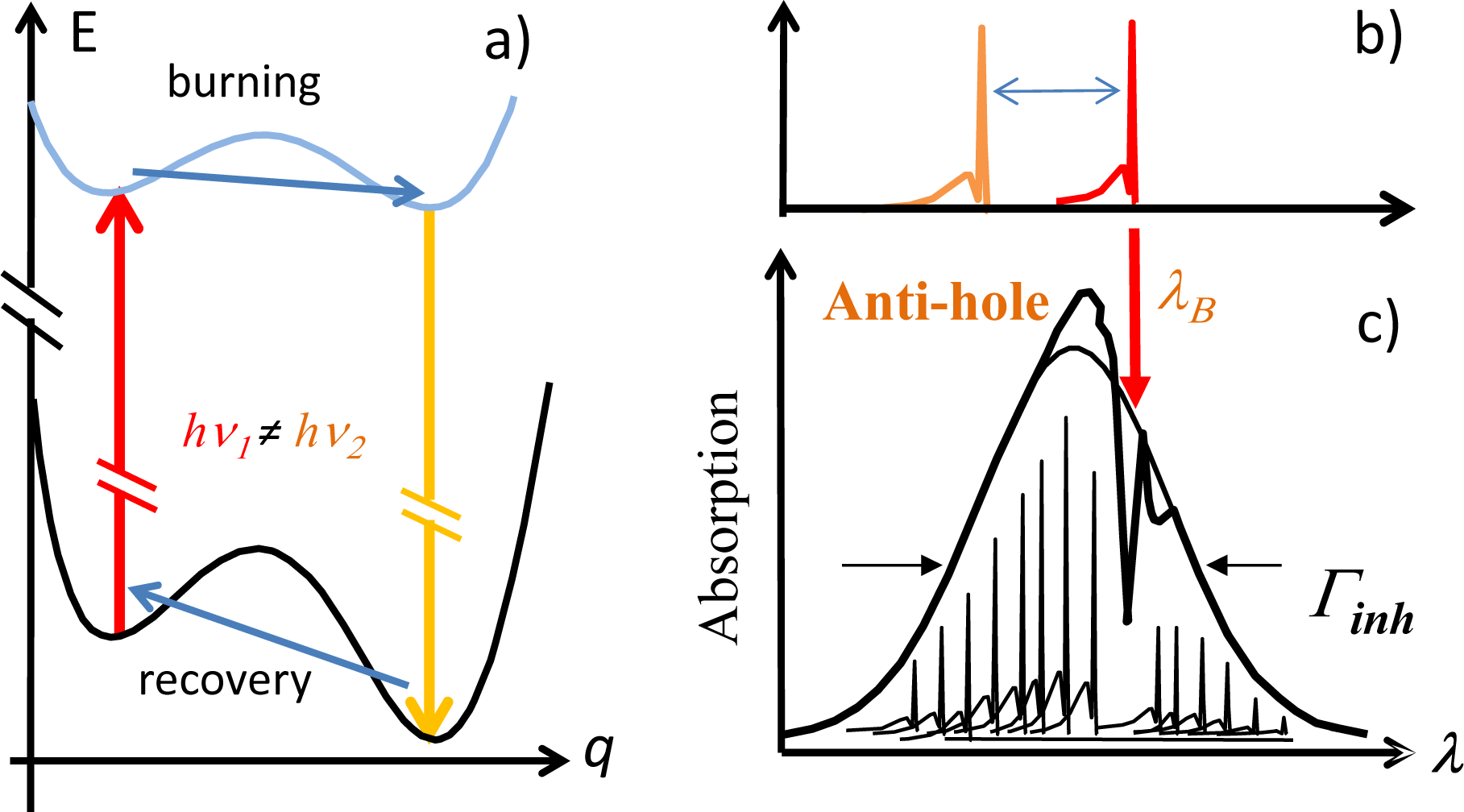
a): Proposed mechanism behind NPHB and single molecule line shifts. Before absorbing a resonant photon, the pigment-protein system is assumed to be in the left well. Upon absorbing a photon, the system finds itself in the left well of the excited electronic state, where the barriers are lower. It has a reasonable ( 10*^−^*^4^) chance to tunnel through the barrier within the ns lifetime of the excited state. Subsequently, the pigment returns to the ground electronic state. Pigment’s transition energies are different for the left and right wells, i.e. a spectral shift is observed, as schematically indicated in (b). The system may return to the original (left) well, but it will take much longer time than in the excited state. (c) In case a macroscopic sample with an inhomogeneously-broadened spectrum Γ*_inh_* is illuminated with a narrow-band laser, a resonant spectral hole will be formed as a result of multiple molecules’ lines shifting out of resonance. These shifted lines contribute to the “anti-hole” at wavelength *λ_B_*.

In this work we employ molecular dynamics (MD) simulations to analyze the dynamics of a representative pigment-protein complex which is known to exhibit NPHB to identify candidates for the generalized coordinate responsible for experimental observations, and to provide support for the TLS/MLS model that has been put forward as the most feasible explanation for observed spectral shifts. MD simulations have been previously used to identify multi-well energy landscapes in proteins other than those involved in pigment-protein complexes.^23,24^ Although many MD simulations of photosynthetic pigment-protein complexes have been reported,^25–29^ none have focused specifically on determining TLS/MLS-like features of their energy landscapes. Previous attempts to combine MD with simulations of optical spectroscopy focused mainly on calculating the time-dependent transition frequencies of the pigments and inter-pigment couplings in the context of studying exciton effects, energy transfer, and charge transfer processes. While distributions of transition frequencies for specific pigments have been occasionally reported,^25^ they have shown no evidence of TLS/MLS. However, TLS/MLS must be present, or (experimentally observed) NPHB and spectral shifts in single complex experiments would be impossible. We hypothesize that classical simulations are well-suited to identify initial candidate generalized coordinates demonstrating TLS/MLS signatures for later investigation in a higher-resolution manner with a focus on specific quantum-related phenomena.

The water-soluble chlorophyll-binding protein (WSCP) is a homotetramer containing one chlorophyll molecule per monomer. Thus, in the first approximation, there is only one type of local environment for each pigment molecule. In the second approximation, the WSCP can be considered to be a dimer of dimers in which the porphyrin rings of the pigments within a dimer are in closer contact (average intra-dimer Mg atoms distance of 0.1 nm) leading to strong intra-dimer excitonic couplings. The dimers are reported to be weakly coupled with inter-dimer Mg atoms being on average 0.2 nm apart.^30^ Unlike most of the pigment-protein complexes from photosynthesizing organisms, WSCP is not a membrane-bound protein and is not directly involved in photosynthetic reactions. It has been suggested that this protein acts as a scavenger of free Chls, transporting them from the thylakoid membrane to the sites of catabolic reactions.^31,32^ It does, however, exhibit NPHB.^7^ Results obtained for the WSCP in this work can be utilized for deliberate and targeted search for the origins of spectral shifts in various reaction center and antenna complexes.

Here, our molecular dynamics (MD) simulations allow us to analyze the dynamics of a representative pigment-protein complex to identify candidates for the generalized coordinate responsible for experimental observations, and to provide support for the TLS/MLS model that has repeatedly been introduced as a possible explanation for observed spectral shifts. We investigate two possible sets of candidates that might produce signatures originating from TLS/MLS: hydrogen bond rearrangements, and rotational motions of single amino acid side chains. We search for multi-well free-energy landscapes exhibiting energy barriers within the experimentally-observed range. We find such signatures in the rotational configurations of a subset of residues in the local environment of the pigment, as well as a single residue moderately correlated with them, which is not in the local environment. We argue based on the symmetry of the energy landscape and the approximate size of the moment of inertia, that local serine residues are most likely to be responsible for the observed experimental signatures.

## 2 Methodology

Although most proposed mechanisms of NPHB and hole recovery are quantum-mechanical in nature (i.e. are dominated by tunneling, at least at low temperatures), we employ unbiased classical MD to allow for all-atom simulation of comparatively long timescales in a comparatively large system, and to pinpoint the exact potential atomic origins of spectroscopic phenomena. Necessarily, this introduces some limitations, and the candidates for relevant residues that we identify with these simulations represent a starting-point for future modelling with the quantum mechanical aspect reintroduced, using QM/MM or DFT approaches. Although we could have employed enhanced sampling methods such as parallel bias metadynamics^33^ or replica exchange sampling^34^, we choose to use unbiased simulations to limit the use of computational resources on a comparatively large system, as well as because we assume we are searching for small-scale motions on the order of individual residues, rather than large-scale slow rearrangement of the entire complex. The latter would be unlikely to result in spectral shifts with the parameters observed in low-temperature experiments.

### 2.1 Molecular Dynamics Simulations

Molecular dynamics simulations were carried out using Gromacs 2021.4 software.^35^ The applied CHARMM36 force field (July 2021 version) is taken from the Mackerell lab website.^36^ The initial crystal structure of class II-B WSCP from *Lepidium virginicum* was obtained from the Protein Data Bank (PDB ID : 2DRE).^30^ Although there is another available structure in the PDB for the same complex, it has slightly lower resolution; moreover, we choose this initial structure since it was for previous MD simulations of the complex. ^37^ The natural Chl a/b ratio of this particular WSCP complex is around 1.6-1.9. However, in this specific crystal structure, only chlorophylls *a* are present. Unlike larger pigment-protein complexes involved in charge transfer reactions, such as Photosystem II, Cytochrome b_6_f, and Photosystem I, that are embedded in a thylakoid membrane, WSCP is simpler and easier to study through MD simulations due to its smaller size and water-based environment. This computational cost reduction allows for longer simulation times, making WSCP an ideal model system for initial exploration of the dynamics and interactions between proteins and chlorophylls through MD simulations.

Missing protein residues (Chain A: residues ILE 1, ASP 163, ASP 164, ASP 165, SER 166, ASP 167, GLU 168, THR 180; Chain B: residues ILE 1, ASN 2, ASP 3, TYR 139, ASP 140, ASN 141, THR 180; Chain C: residues ILE 1, ASN 2, THR 180; Chain D: residues ILE 1, ASN 2, THR 180) were built using the Modeller program, assuming that intact WSCP is a homotetramer.^38^ The protein chirality and the cis / trans configuration of the peptide bonds were verified using the VMD 1.9.4a43 plugins Chirality 1.4 and Cispeptide 1.4.^39^ Chlorophyll *a* parameters, which had to be added to the force field, were obtained from previous MD simulations of the WSCP.^40–45^ Acidic amino acids (ASP, GLU) were modeled as deprotonated and basic amino acids (LYS, ARG, and HIS) were modeled as protonated, as has been done for previous modeling of this complex.^37^ The pKa values of these residues were checked using the PropKa website (https://www.ddl.unimi.it/vegaol/propka_about.htm)^46^ and the results are shown in the Supplementary Information (Figs S1-S2). Although within a protein the pKa of amino acids may change significantly, resulting in different protonation states, our analysis indicates that in WSCP the vast majority of the amino acids obey the simple assumption above and for a protein of this size it was not feasible to perform constantpH MD. The resulting negatively charged system was neutralized by replacing 40 random solvent molecules with 40 sodium ions. The system was energy-minimized using the steepest descent algorithm to a maximum force of 1000.00 kJ/mol/nm, with a maximum number of 50,000 steps of energy step size is 0.01. The protein complex was constrained in a rhombic dodecahedron box with an image distance *d* = 11.757 *nm*^3^, and solvated with 33,774 water molecules in addition to 555 water molecules existing in the crystal structure, for a total of 34,329 TIP3P-CHARMM water molecules.^47^ Periodic boundary conditions were used to ensure a continuous representation of the system and avoid artificial interactions caused by the finite size of the simulation box. The initial velocities of the particles were randomly assigned from a Maxwell-Boltzmann distribution at 300 K. Equilibration steps (NVT and NPT) ran for 1000 ps each to reach respectively the target temperature and the target pressure. For NVT equilibration, the Bussi-Donadio-Parrinello thermostat^48^ (referred as the v-rescale thermostat in Gromacs) was used with a coupling time of *τ_t_* = 0.1 at a temperature of 300K. For NPT equilibration, the Berendsen barostat^49^ was used and the target pressure was set to 1 bar.

Production simulations were performed using the same thermostat and a Parrinello-Rahman barostat^50^ with a coupling constant of *τ_p_* = 2.0 set to a reference pressure of 1 bar as done previously in simulations of the same complex.^37^ To ensure reproducibility and reliability, 5 replicas for each simulation were performed, starting from the same initial energy-minimized structure and assigning different random initial velocities of the particles for all replicas. The equations of motion were integrated using the leapfrog algorithm with a time step of 2 fs.^51^ Hydrogen bonds were constrained with the LINCS algorithm.^52^ Each replica was simulated for 1 *µ*s for a total of 5 *µ*s and, in the analysis, results were averaged across replicas, with standard errors computed from the differences between them. Simulations were conducted at 300 K; although NPHB experiments are performed at cryogenic temperatures, such temperatures are not accessible in MD simulations. While interpreting NPHB data it is usually assumed that the landscape itself is temperature-independent, and only the rates of tunneling and thermally-activated barrier-hopping are temperature-dependent, thus we assume that our results will be able to explain NPHB data.

## 3 Results

### 3.1 Definition of local environment

The tetrameric WSCP complex is a large protein comprising a total of 720 residues. We identified the local protein environment (Fig. S4a) of the pigments to investigate possible direct interactions responsible for experimental spectral dynamics.^7^ The reference paper of this crystal structure^30^ listed 42 residues per chain (a total of 168 residues) forming the hydrophobic cavity in which Chls *a* are enclosed. We ensured that introducing dynamics into the system did not bring other residues within an all-atom cut-off distance of 0.3 nm using a contact map analysis of the simulations computed with the Contact Map Explorer Python package^53^ (cf. Sec. S2). 66.7% of the residues in the local environment are non-polar, 21.4% are polar, while 7.1% are basic, and only 4.8% are acidic (Fig. S4b).

### 3.2 Identification of long hydrogen bonds; no evidence of classical multimodal signature

The purpose of this section is two-fold. First, it is known that hydrogen bonds with lengths within the range of 0.25–0.29 nm exhibit double-well energy landscapes, making them potentially responsible for TLS behavior.^54^ However, since such potentials are associated with quantum tunneling of protons, classical MD cannot be used to find associated energy landscapes or determine their energy barriers. Thus, such hydrogen bonds, if identified in the vicinity of the pigment, will be set aside for more advanced QM analysis (future work). Second, we probed whether conformational rearrangements of such bonds might lead to *classical* double-well potentials. In other words, we investigate both the presence of such bonds and the possibility that a protein-level conformational change resulting in bond rearrangement reflected as two distinct conformations between donor and acceptor atoms participating in h-bonding (cf. Fig. S5) might explain the experimentally-observed spectral dynamics. To investigate this, we computed the distance distributions between these atoms for each hydrogen bond. We identified three residues forming multiple long hydrogen bonds with the chloropylls, none of which were associated with classical double-well potentials.

Hydrogen bonds between the chlorophylls and the protein residues were calculated with the Baker-Hubbard Hydrogen Bond identification function from the Python library MdTraj 1.9.6.^55,56^ Only hydrogen bonds between 0.25–0.29 nm long with an *α* angle Donor-Hydrogen-Acceptor greater than 120 degrees were retained for further investigation. The analysis revealed that each Chlorophyll *a* established long hydrogen bonds with three specific residues, namely Threonine 52 (THR52), Serine 53 (SER53), and Glutamine 57 (GLN57), as illustrated in Fig. 2a. For all these H-bonds, Chlorophyll *a* functions as the acceptor, leaving protein residues to serve as the donors. The central magnesium atom in Chl *a* does not participate in hydrogen bonding due to its relatively low electronegativity, which results in a weaker attraction for electrons compared to highly electronegative atoms. Each residue establishes two H-bonds with its corresponding pigment, resulting in a total of six distinct hydrogen bonds formed per pigment, as illustrated in Fig. 2b. Specifically, Chl *a* forms two H-bonds with the hydroxyl side chain group of THR52, respectively referred to as THR52 sc1 and THR52 sc2, where sc stands for side chain. Additionally, Chl *a* forms one H-bond with the amine backbone group of SER53, denoted as SER53 bb (where bb stands for backbone), and one H-bond with its hydroxyl side chain group, denoted as SER53 sc. Finally, Chl *a* also forms two H-bonds with the hydrogens in the amide side chain group of GLN57, respectively labeled as GLN57 sc1 and GLN57 sc2. The oxygen atoms of Chl *a* involved in these H-bonds possess two lone pairs of electrons, enabling them to form two H-bonds simultaneously. As a result, SER53 bb and SER53 sc1 can coexist, as can GLN57 sc1 and GLN57 sc2. However, since the hydrogen atom within the residues only has one electron, it can only form a single H-bond at a time. Consequently, THR52 sc1 and THR52 sc2 cannot coexist.

**Figure 2:**
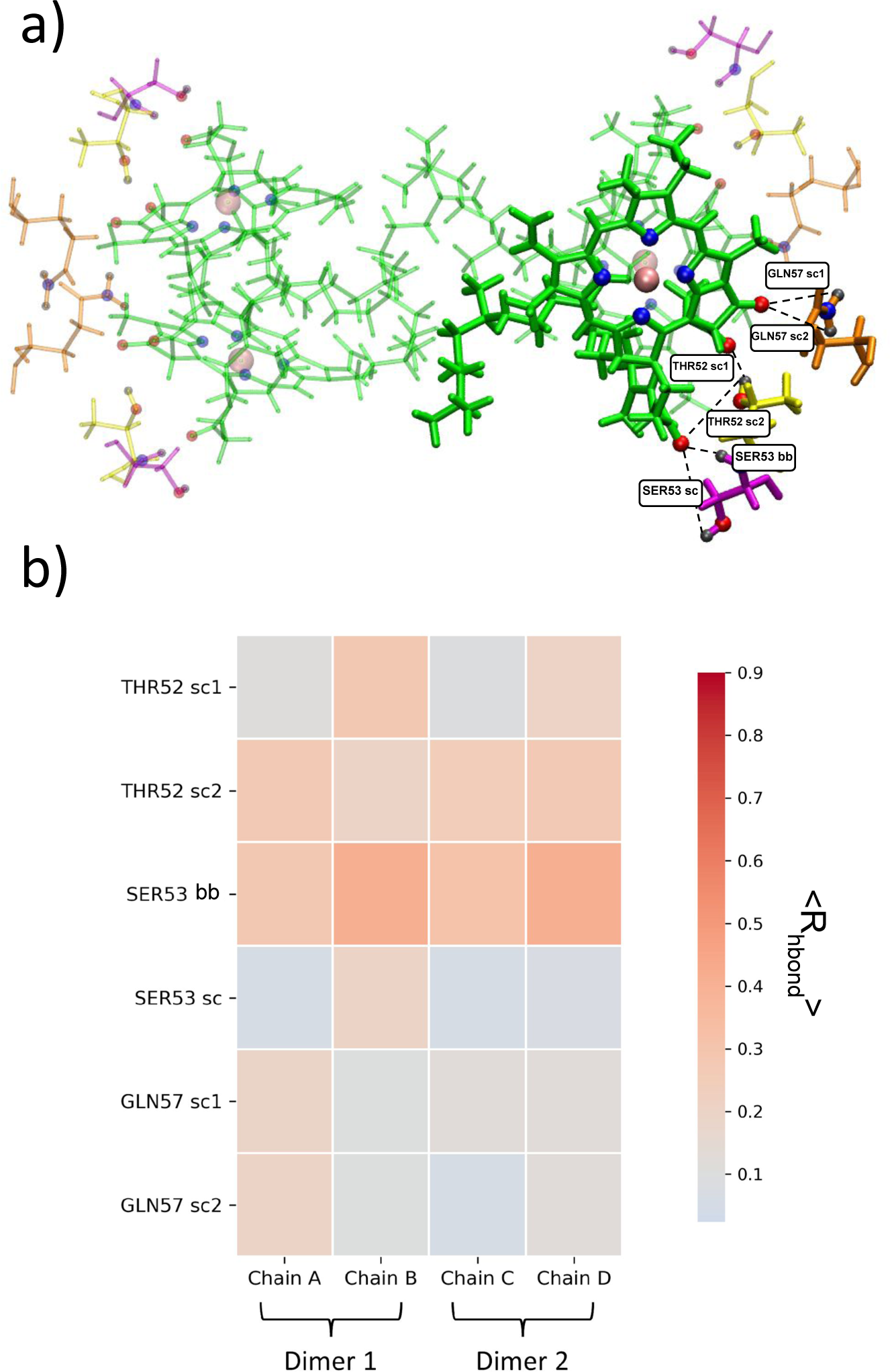
Long hydrogen bonds formed in the local environment. Representations of a) all Chlorophylls *a* (green) interacting with their Threonine 52 (yellow), Serine 53 (magenta), and Glutamine 57 (orange). Oxygen atoms involved in the H-bonds are colored in red, hydrogens in black, and nitrogens in blue. Three of the four symmetric subunits are rendered transparent for clarity. The central Mg atom of each Chl *a* is colored in pink and their nitrogen atoms are also shown in blue. All possible h-bonds are marked by black dashed lines. (b) Heatmap of average occurrence *R*_hbond_ taken across five independent replicas of long hydrogen bonds. In Fig. S7, the standard error is shown.

The occurrence map of the identified hydrogen bonds (Fig. 2c) was computed for each protein chain in the simulation. The data were obtained from five replicas and subsequently averaged. The purpose of these maps was to determine the frequency at which hydrogen bonds occur, as transient hydrogen bonds may indicate potential dynamic behavior in the system. We observed that the majority of hydrogen bonds occurred between 10% and 60% of the simulation time, and the occurrence patterns were relatively consistent across the protein chains. The SER53 bb hydrogen bond is the most persistent, followed by the THR52 sc2 hydrogen bond. The GLN57 hydrogen bonds and the SER53 hydrogen bonds are the least persistent.

To further probe persistence of hydrogen bonds, the average distances between the hydrogen atom and the acceptor atom involved in each hydrogen bond were computed over all frames of the simulations and are presented in Table 1. In most cases, the average distance exceeds our cut-off distance of 0.29 nm; only the SER53 bb hydrogen bond has an average of 0.29 nm or below for all chains, although the GLN57 sc1 hydrogen bond’s mean value is within a single standard deviation of 0.29 nm for all chains. Based on this analysis, other hydrogens bonds are primarily transient, and the SER53 bb and GLN57 sc1 are the primary candidates for further study with a quantum mechanical approach, such as QM/MM or density functional theory (DFT).^57^ We find no evidence of classical double-well potentials due to rearrangement of hydrogen bonds in our analysis; all distributions of distances between donor and acceptor atoms are unimodal.

**Table 1:** Average distance between the hydrogen atom and the acceptor atom for six hydrogen bonds. Distances are averaged over five replicas for each protein chain. Standard deviation of the means are displayed on the right side of the columns. Significant figures are calculated out to the range of standard error.

| Hydrogen Bonds | Average distance Hydrogen-Acceptor (nm) |  |  |  |
| --- | --- | --- | --- | --- |
|  | *Chain A* | *Chain B* | *Chain C* | *Chain D* |
| THR52 sc1 | $0.39 \pm 0.09$ | $0.35 \pm 0.1$ | $0.43 \pm 0.1$ | $0.41 \pm 0.1$ |
| THR52 sc2 | $0.40 \pm 0.1$ | $0.34 \pm 0.1$ | $0.46 \pm 0.1$ | $0.42 \pm 0.1$ |
| SER53 bb | $0.29 \pm 0.1$ | $0.22 \pm 0.07$ | $0.28 \pm 0.1$ | $0.22 \pm 0.04$ |
| SER53 sc | $0.42 \pm 0.1$ | $0.36 \pm 0.1$ | $0.44 \pm 0.1$ | $0.38 \pm 0.09$ |
| GLN57 sc1 | $0.28 \pm 0.1$ | $0.34 \pm 0.1$ | $0.30 \pm 0.09$ | $0.34 \pm 0.1$ |
| GLN57 sc 2 | $0.39 \pm 0.07$ | $0.40 \pm 0.09$ | $0.41 \pm 0.09$ | $0.42 \pm 0.1$ |

### 3.3 Classical multimodal distributions in side chain dihedral angles and associated free energy surfaces

#### 3.3.1 Evidence of multimodal distributions in side chain motions of certain residues

To quantify positional changes of residues in the local environment of the chlorophylls and identify potential multimodal signatures, we calculated root-mean-square deviation of the whole residues. RMSD measures the deviation of atoms in a substructure relative to a reference. In Gromacs 2021.4, the RMSD is defined as^35^ :

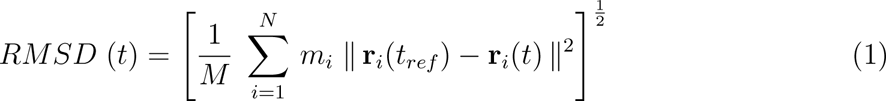

where 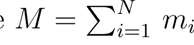, the mass of all particles in a substructure, **r***_i_*(*t_ref_* ) is the position of a particle *i* at the reference time *t_ref_* , and **r***_i_*(*t*) is the position of a particle *i* at time *t*. RMSD (and root mean square fluctuation, RMSF) are good predictors of multi-well landscapes as they provide indications that some residues can be in more than one state with significantly different degrees of flexibility. In this analysis, the 168 residues were treated as separate substructures for RMSD calculation, with the porphyrin ring of the Chl *a* in the first frame of the simulation serving as the reference substructure for all of them (cf. Fig. S8). Of these 168, we identified 55 displaying bimodal RMSD distributions in one or more replicas (cf. Figs S9-S16). It is of interest to note that THR52 (Fig. S12, row 3) and GLN57 (Fig. S13, row 1), which were identified in the previous section as participating in the long hydrogen bonding analysis, are among these 55 residues.

Table 2 presents the average characteristics of the RMSD distributions for the 55 identified residues, considering only the replicas that exhibit a double-peak distribution. The small standard error values for all parameters at both temperatures demonstrate an agreement among those replicas that demonstrated sufficient sampling, indicating that they have captured the same phenomenon for the residues. The average first peak position is below 0.1 nm, indicating a stable initial position.^58^ The second peak position is above 0.2 nm, indicating an apparent shift from the initial position.^58^ Finally, the average distance between the peaks is 0.153 ± 0.006 nm. Therefore, it can be concluded that all residues undergo the same type of structural change, regardless of their type.

**Table 2:** Characteristics of Bimodal RMSD Distributions. Results are averaged over the 55 residues. Standard error of the means taken over 5 independent replicas are displayed on the right side of the columns.

| Characteristics of Bimodal RMSD Distributions |  |
| --- | --- |
| First peak position (nm) | $0.097 \pm 0.006$ |
| Second peak position (nm) | $0.250 \pm 0.010$ |
| Distance between peaks (nm) | $0.153 \pm 0.006$ |

The 55 residues exhibiting bimodal RMSD distributions were categorized by their side chain physico-chemical properties, as shown in Fig. 3. Comparing this classification to the classification of the full local environment (Figure S4, Secs 3.1, S2), we observed that the proportion of residues per category is preserved. Among the 55 residues, 65.5% are non-polar, which is similar ratio considering all 168 residues where 66.7% of them were nonpolar. Additionally, 25.5% of the 55 residues are polar, compared to 21.4% in the overall 168 residues classification. Furthermore, 9.0% of this reduced set of residues have a basic side chain, while the overall proportion is 7.1%. Notably, none of the 8 aspartic acid residues of the local environment displayed a double-peak RMSD distribution, resulting in no acidic residues among the selected 55 residues. Therefore, the main difference observed is the absence of acidic residues. From this classification, it seems that the presence of bimodal RMSD distributions does not appear to be influenced by the general physicochemical properties of the residue.

**Figure 3:**
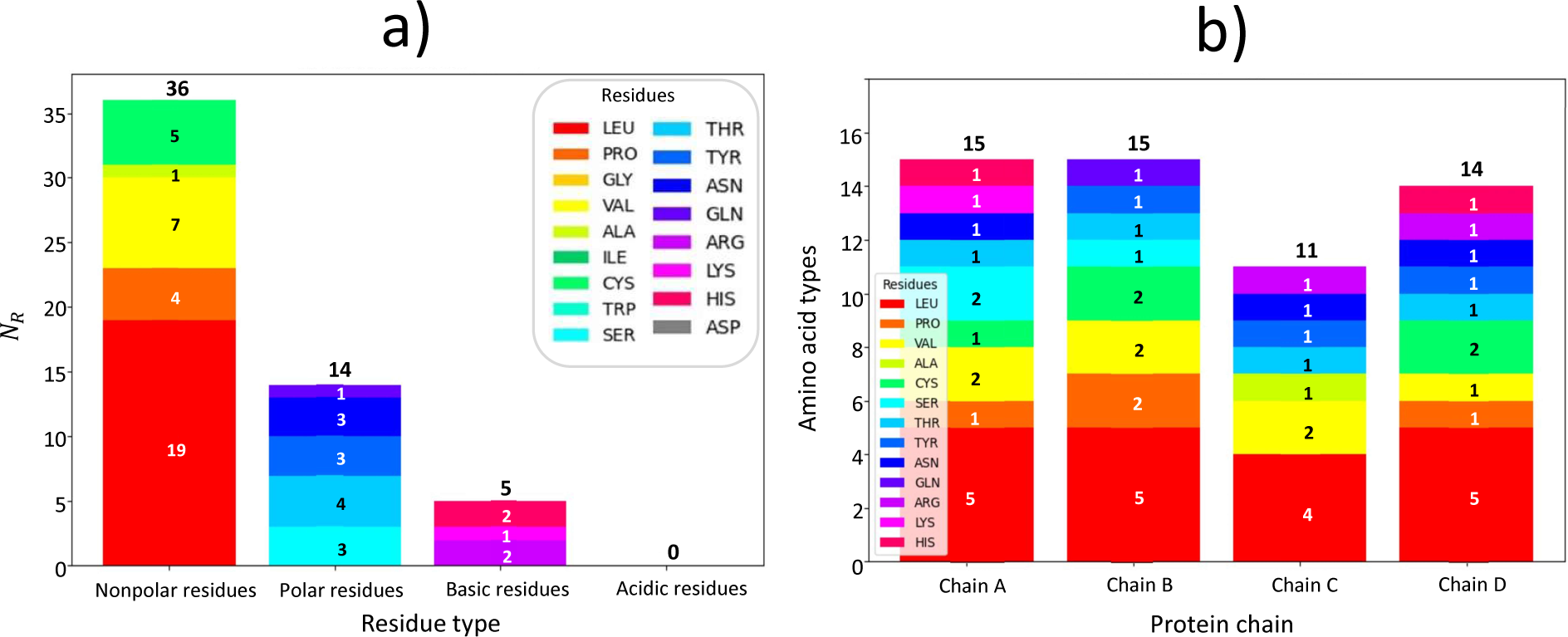
(a) Numbers of residues *N_R_* of different types comprising the 55 residues associated with observed multimodal distributions in the local environment; (b) Amino acid types per chain of the residues comprising the 55 residues associated with observed multimodal distributions in the local environment

The 55 residues were also classified by protein chains, as shown in Fig. 3b. While the amino acid types are generally evenly distributed among the four protein chains, there are slight variations. Notably, protein chain C has only 11 residues displaying a bimodal distribution. Additionally, there are differences in the residue composition. For instance, protein chain C is the only chain that lacks PRO residues exhibiting an RMSD shift but is the only chain containing an ALA residue. These subtle differences may be partially attributed to the fact that although the protein sub-units are chemically identical, they are not fully identical geometrically in the crystal structure (dimer of dimers).^30^ As a result, each protein chain may exhibit slight dynamic variations, leading to the observed distinctions. We note also that this could indicate a lack of full convergence of the employed trajectories. As 5 *µ*s is a substantial amount of simulation time and the effect is moderate, we have not extended the simulations further, but for future simulations, we intend to investigate the use of enhanced sampling, such as metadynamics.

While the RMSD analysis provides valuable insights, it does not directly identify relevant generalized coordinates. We found that the above 55 residues exhibiting bimodal RMSD distributions also exhibited bimodal or trimodal distributions of their side chain dihedral angles (for a comprehensive visualization, see Figs S17-S24). The *χ*_1_ angle is the first side chain dihedral angle in a protein residue (see Fig. 4 and section S4.1 in the Supplementary Information for more details). The proline (PRO) residue has a unique pyrrolidine ring structure, which imposes limitations on the conformational flexibility of its side chain. Consequently, PRO does not possess a *χ*_1_ angle. Instead, calculations for PRO were based on the *ω* angle, which reflects the rotation around the peptide bond connecting the nitrogen atom of the pyrrolidine ring and the carbon atom of the backbone carbonyl group (cf. Fig. S25e-f). We note that an analysis of the root-mean-square fluctuations (RMSF) and residue center of mass motions supports the assertion that motion of the side chain dihedrals is the origin of the bimodal distributions observed in the RMSD (cf. Sec. S4.3 in the Supporting Information.) Although the distributions sampled by the residues of different replicas are not identical, and there is occasional evidence of kinetic trapping in certain replicas (cf. nearly bimodal distribution of HIS 43, chain A, replica 5, Fig. S18, top left), we believe that the overall sampling of 5 *µ*s for the complex is sufficient to at least correctly approximate the free energy landscape features (see below). Figure 5 visually represents the arrangement of these 55 residues within the protein complex.

**Figure 4:**
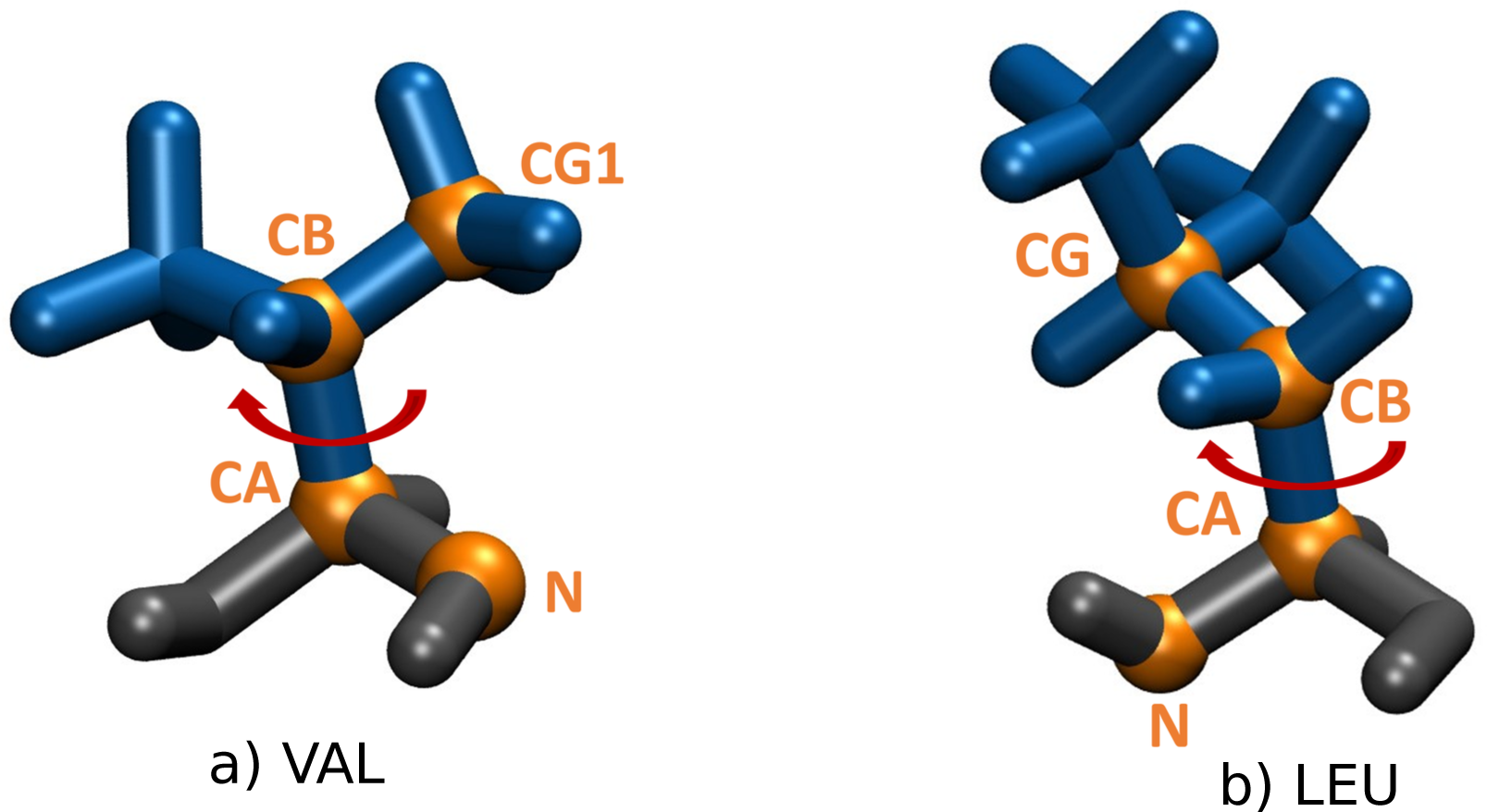
Example of *χ*_1_ angle. Graphical representations of the angle for (a) VAL residue, and (b) LEU residue. Backbone atoms are colored in black, side chain atoms in blue, and atoms used for *χ*_1_ angle calculations are colored and labeled in orange. Rotation of the *χ*_1_ angle are represented by a red arrow.

**Figure 5:**
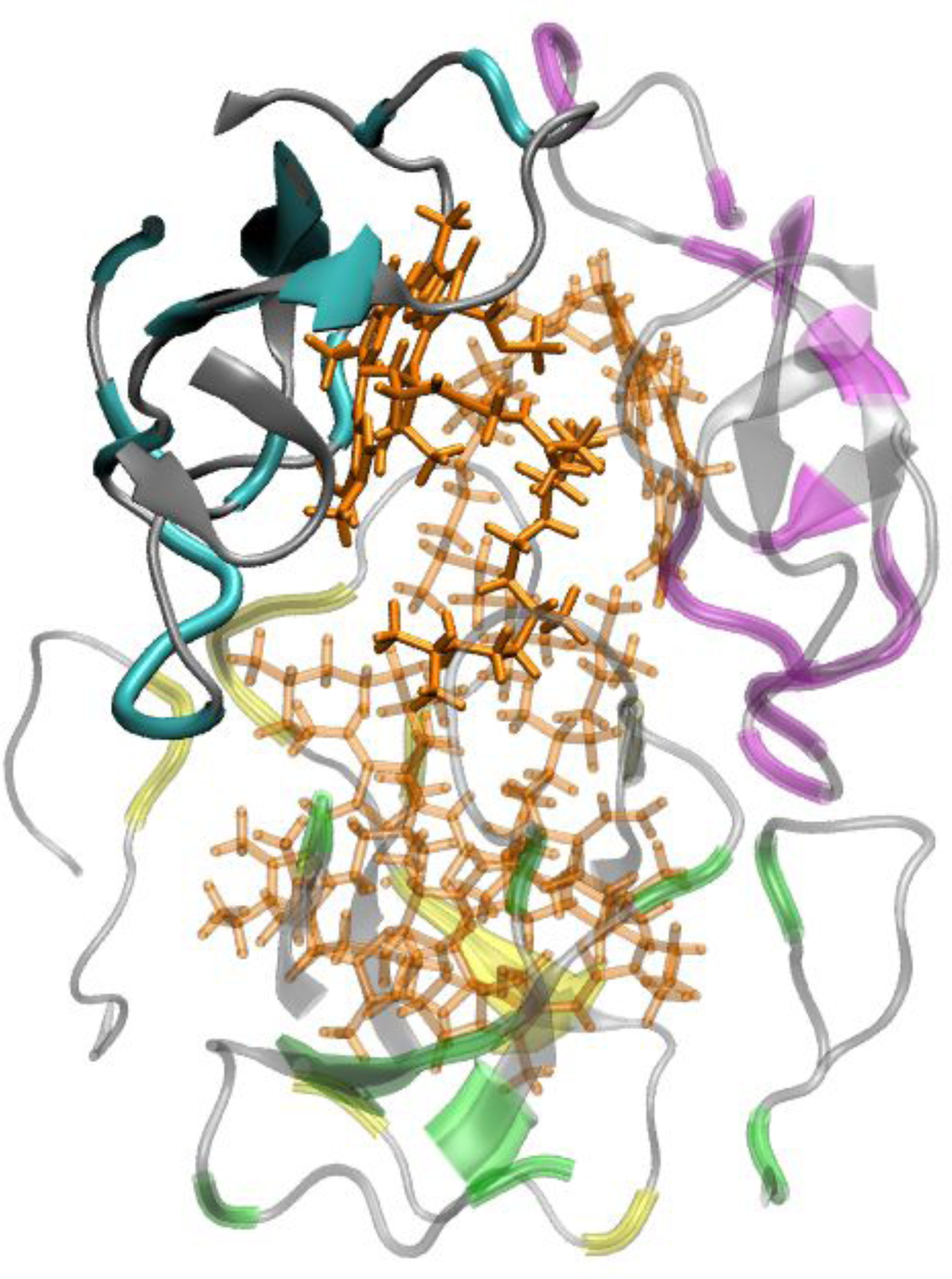
Residues with side chains whose conformational angles demonstrate a multimodal distribution. Shown are the four chlorophylls, in orange, surrounded by the hydrophobic cavity, with residues demonstrating multimodal distributions colored according to their respective chains. Three of the four subunits are rendered transparent for clarity.

Table 3 summarizes the characteristics of the *χ*_1_ angle distributions of the residues, considering only the replicas that exhibit a triple-peak distribution. The peaks are not exactly equally spaced, with the first and third peaks being on average the closest, considering periodicity of the angles. (For a discussion surrounding the correspondance of bimodal RMSD distributions with trimodal *χ*_1_ distributions, see Sec. S4.2 in the Supporting Info).

**Table 3:** Characteristics of Trimodal *χ*_1_ Angle Distributions: first peak position, *χ^∗^*, second peak position, *χ^∗^*, third peak position, *χ^∗^*, and the angular differences between the peaks, Δ*χ^∗^_12_* , Δ*χ^∗^_23_* , Δ*χ^∗^_31_* . Results are averaged over per residue type. Standard error of the means taken over 5 independent replicas are displayed on the right side of the columns.

| Amino Acid | Characteristics of Trimodal Dihedral Angle Distributions($^\circ$) | | | | | |
| --- | --- | --- | --- | --- | --- | --- |
| | $\chi_1^*$ | $\chi_2^*$ | $\chi_3^*$ | $\Delta\chi_{12}^*$ | $\Delta\chi_{23}^*$ | $\Delta\chi_{31}^*$ |
| VAL | $-129 \pm 25$ | $21 \pm 32$ | $115 \pm 29$ | $150 \pm 57$ | $94 \pm 61$ | $116 \pm 54$ |
| LEU | $-122 \pm 22$ | $-7 \pm 40$ | $122 \pm 25$ | $115 \pm 62$ | $129 \pm 65$ | $116 \pm 47$ |
| CYS | $-129 \pm 25$ | $-21 \pm 28$ | $129 \pm 15$ | $108 \pm 53$ | $150 \pm 43$ | $102 \pm 40$ |
| SER | $-129 \pm 29$ | $-7 \pm 43$ | $129 \pm 29$ | $122 \pm 72$ | $136 \pm 72$ | $102 \pm 58$ |
| THR | $-129 \pm 10$ | $-7 \pm 44$ | $129 \pm 25$ | $122 \pm 54$ | $136 \pm 69$ | $102 \pm 35$ |
| ALA | $-129 \pm 29$ | $0 \pm 40$ | $122 \pm 33$ | $129 \pm 69$ | $122 \pm 73$ | $109 \pm 62$ |
| LYS | $-122 \pm 26$ | $14 \pm 35$ | $122 \pm 37$ | $136 \pm 61$ | $108 \pm 72$ | $116 \pm 63$ |
| GLN | $-125 \pm 25$ | $19 \pm 42$ | $123 \pm 31$ | $144 \pm 67$ | $104 \pm 73$ | $112 \pm 56$ |
| TYR | $-122 \pm 25$ | $-28 \pm 35$ | $136 \pm 25$ | $94 \pm 60$ | $164 \pm 60$ | $102 \pm 50$ |
| ASN | $-129 \pm 24$ | $14 \pm 32$ | $108 \pm 25$ | $143 \pm 56$ | $94 \pm 57$ | $123 \pm 49$ |
| ARG | $-136 \pm 21$ | $7 \pm 36$ | $129 \pm 25$ | $143 \pm 57$ | $122 \pm 61$ | $95 \pm 46$ |
| HIS | $-129 \pm 22$ | $-14 \pm 43$ | $122 \pm 32$ | $115 \pm 65$ | $136 \pm 75$ | $109 \pm 54$ |
| PRO | $-115 \pm 29$ | $115 \pm 36$ | | $130 \pm 65$ | | |

In summary, our analysis identified 55 residues exhibiting a bimodal RMSD distribution.

Of these 55, all proline residues exhibited a bimodal distribution of their *ω* angle, and all other residues exhibited a trimodal distribution of their *χ*_1_ side chain angle. Consistent results were obtained across all four protein chains. For PRO residues, this indicates the existence of two energetically preferred side chain conformations as shown in Supporting information in Figure S25; for all other residues, it indicates the existence of three. The conformational changes did not appear to be influenced by the side chain type (non-polar, polar, acidic or basic) of residues or the distance between the residues and their corresponding pigments, confirming that the RMSD bimodality arises from side chain rotation rather than other motions in relation to the Chl *a* pigments.

#### 3.3.2 Protein free energy landscape associated with the *χ*_1_ angle rotation

The multidimensional free energy landscape of the protein can be simplified and represented along a single generalized coordinate *x*. This simplification enables us to gain insights into the system behavior in a reduced-dimensional space. The free energy *F* along a coordinate *x* is expressed by the well-known equation:^59^

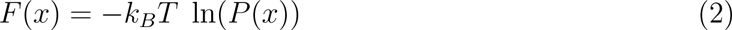

where *k_B_*is the Boltzmann constant, *T* is the temperature, and *P* (*x*) represents the probability distribution function of *x*.

We constructed one-dimensional free energy landscapes associated with the *χ*_1_ and *ω*_1_ values of different amino acids to assess energy barrier heights between the wells for comparison with experimental observations. To obtain the one-dimensional free energy landscapes, we approximate the continuous probability distribution by binning *χ*_1_ values into equally-spaced bins from −180*^◦^* to 180*^◦^*. For each replica, we bin together all occurrences associated with multimodal angular distributions by amino acid. We do not take into account conformations associated with unimodal distributions of residues demonstrating multimodal distributions in other contexts, in order to avoid biasing the estimates of Δ*F* with non-equilibrium data.

For each bin, the probability density was approximated as the number of frames that fell within that particular bin and dividing it by the total number of frames (100,000) in the simulation. It is important to note that any normalization constant causes shift of the landscape up or down in free energy, without altering the relative free energy values. Standard error was estimated from the standard deviation of the means of the five different replicas.

Protein free energy landscapes associated with the *χ*_1_ angles of different residues are depicted in Figs 6-7. We can see that all FELs more or less display three energy wells corresponding to three distinct conformations with three distinct *χ*_1_ angle values, consistent with the trimodal *χ*_1_ angle distributions. Not all multi-well FELs are symmetric: while ALA and SER display similar well depths and barrier heights for all wells and barriers, respectively, the FELs associated with TYR and ASN may be closer to bimodal due to the suppression of the third well. The first energy well corresponds to a preferred conformation around −140*^◦^* for all residues; it is the deepest well for VAL, CYS, LYS, GLN, TYR, ASN, and HIS. The second well corresponds to a preferrsed conformation around 0*^◦^*; it is the deepest well for LEU residues. The third well corresponds to a preferred conformation around 125*^◦^*; it is the deepest well for ARG residues. We note that for SER and ALA residues, all conformations have similar well-depths; while for VAL, CYS, THR, GLN, and HIS, the 2nd and 3rd wells have similar depths, for LYS, the 1st and 3rd have similar depths, and for ARG the 1st and 2nd. Average free energy barrier heights per residue are presented in Table 4. Most average per-residue energy barrier heights (33/38) fall (within error bars) within the range of 1100 to 1600 *cm^−^*^1^ expected to be responsible for experimentally observed spectral signatures, with the exceptions of Δ*F* ^VAL^, Δ*F* ^SER^, Δ*F* ^LYS^, Δ*F* ^ASN^, and Δ*F* ^VAL^.

**Figure 6:**
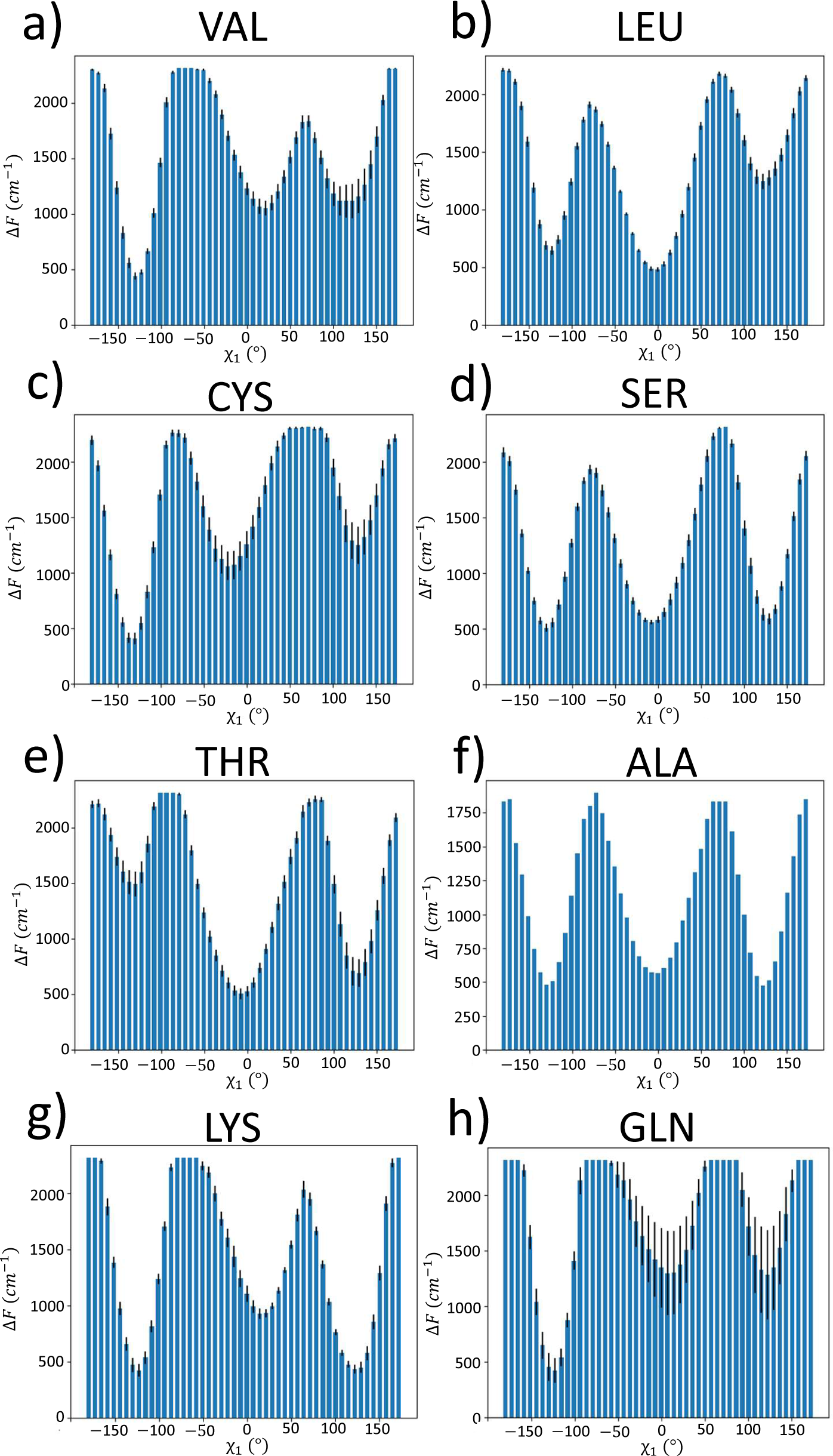
Protein Free energy landscapes associated with the rotation of the *χ*_1_ angle for different amino acid types. Error bars represent standard error of the means taken over 5 independent replicas of the free energy values. Further landscapes shown in Fig. 7

**Figure 7:**
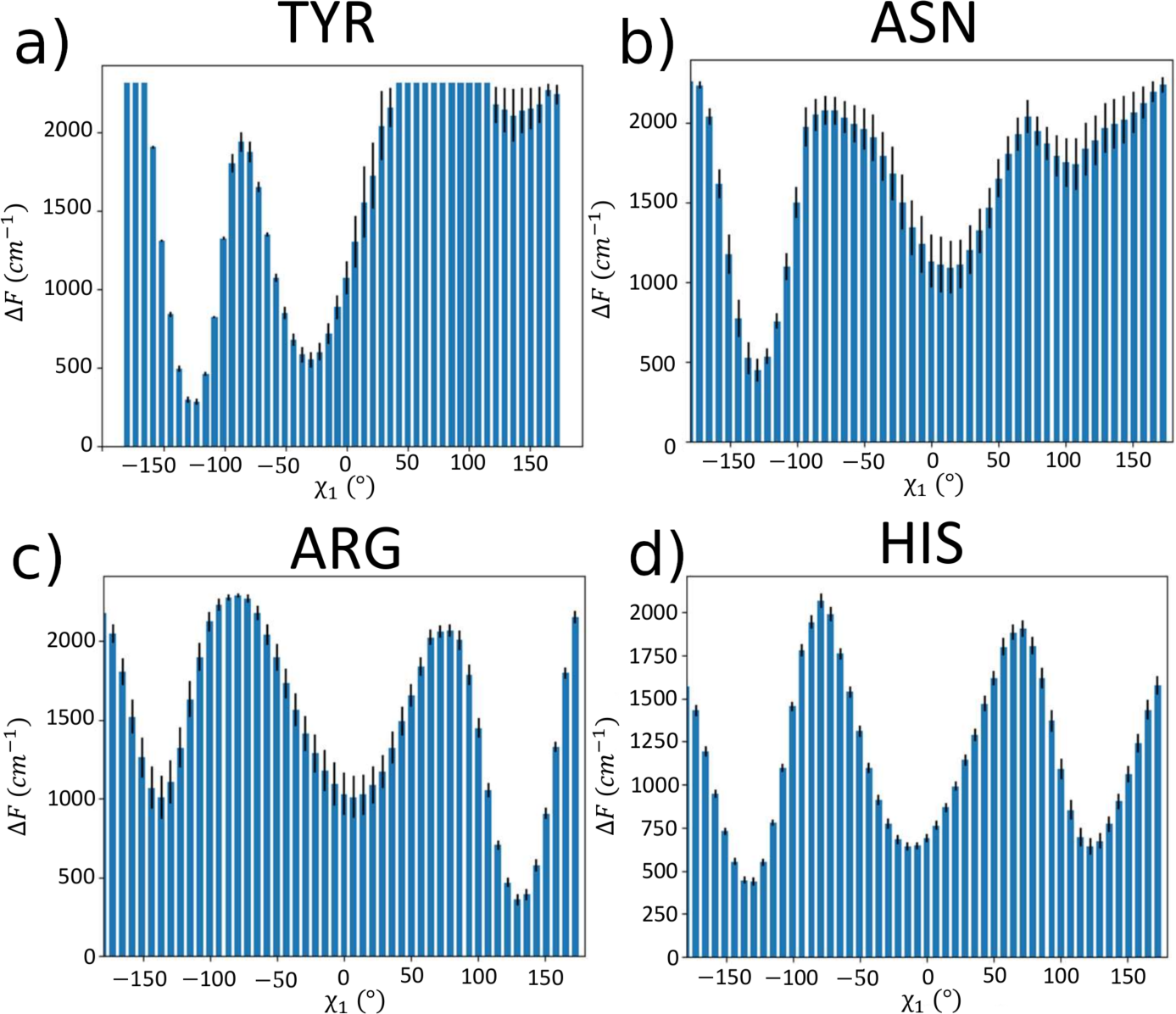
Protein Free energy landscapes associated with the rotation of the *χ*_1_ angle for different amino acid types, cont. Error bars represent standard error of the means taken over 5 independent replicas of the free energy values. The first eight amino acid types are shown in Fig. 6

**Table 4:**
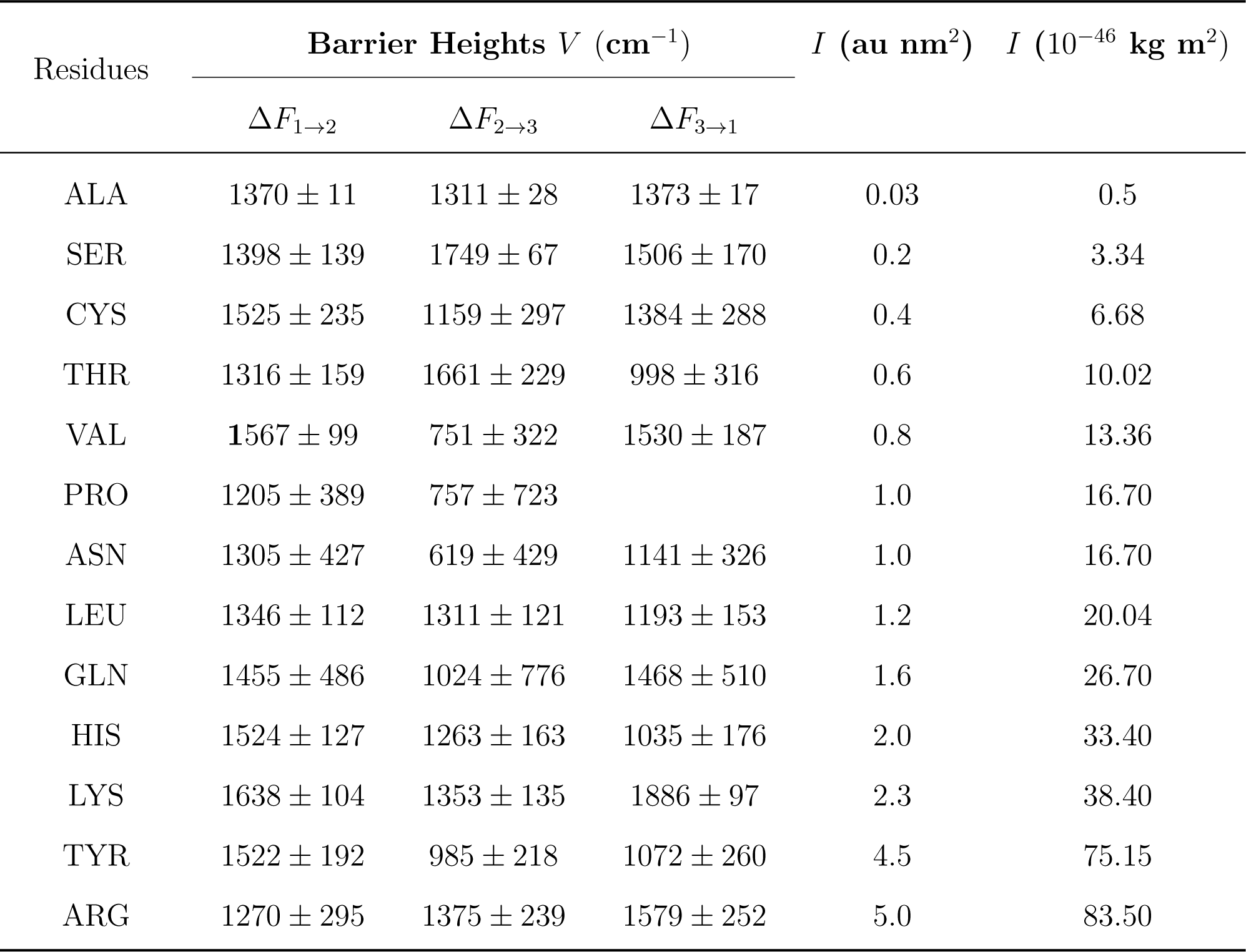
Parameters relevant for NPHB experiments. Energy barrier heights at 300K in units of cm*^−^*^1^ for side chain angles exhibiting multi-modal angledistributions. Standard error is computed from the standard deviation of the means taken over five independent replicas. The rightmost columns include estimated rotational moments of inertia, where au nm^2^ = 1.67 × 10*^−^*^45^ kg m^2^.

| Residues | Barrier Heights $V$ ($\text{cm}^{-1}$) | | | $I$ ($\text{au nm}^2$) | $I$ ($10^{-46} \text{ kg m}^2$) |
| --- | --- | --- | --- | --- | --- |
| | $\Delta F_{1 \rightarrow 2}$ | $\Delta F_{2 \rightarrow 3}$ | $\Delta F_{3 \rightarrow 1}$ | | |
| ALA | $1370 \pm 11$ | $1311 \pm 28$ | $1373 \pm 17$ | 0.03 | 0.5 |
| SER | $1398 \pm 139$ | $1749 \pm 67$ | $1506 \pm 170$ | 0.2 | 3.34 |
| CYS | $1525 \pm 235$ | $1159 \pm 297$ | $1384 \pm 288$ | 0.4 | 6.68 |
| THR | $1316 \pm 159$ | $1661 \pm 229$ | $998 \pm 316$ | 0.6 | 10.02 |
| VAL | $1567 \pm 99$ | $751 \pm 322$ | $1530 \pm 187$ | 0.8 | 13.36 |
| PRO | $1205 \pm 389$ | $757 \pm 723$ | | 1.0 | 16.70 |
| ASN | $1305 \pm 427$ | $619 \pm 429$ | $1141 \pm 326$ | 1.0 | 16.70 |
| LEU | $1346 \pm 112$ | $1311 \pm 121$ | $1193 \pm 153$ | 1.2 | 20.04 |
| GLN | $1455 \pm 486$ | $1024 \pm 776$ | $1468 \pm 510$ | 1.6 | 26.70 |
| HIS | $1524 \pm 127$ | $1263 \pm 163$ | $1035 \pm 176$ | 2.0 | 33.40 |
| LYS | $1638 \pm 104$ | $1353 \pm 135$ | $1886 \pm 97$ | 2.3 | 38.40 |
| TYR | $1522 \pm 192$ | $985 \pm 218$ | $1072 \pm 260$ | 4.5 | 75.15 |
| ARG | $1270 \pm 295$ | $1375 \pm 239$ | $1579 \pm 252$ | 5.0 | 83.50 |

Summarizing, the investigation of the local environment of the pigments in WSCP led to the identification of several different microscopic entities potentially responsible for experimentally observed spectral shifts. The heights of the energy barriers for the residues displaying multimodal distributions were calculated and compared to experimental data. From our simulations, the majority appeared to fall within a similar range of values that agree with the (low-temperature) experimental data. Quantum mechanical calculations would be required to obtain excited-state energy barriers to verify that excited-state barriers are indeed of proper magnitude to explain spectral hole burning and single-complex line shift results.

### 3.4 Dynamical Network analysis identifies an additional residue of interest

While residues that are directly in contact with the pigment are the likeliest candidates for the origin of spectral shifts, we also considered the possibility that allosteric communication between local and distal residues might allow contributions from residues outside of the local environment. We investigate this possibility through the use of dynamical network analysis to see if there were significant correlations between any local residues of interest and any distal residues, and identify a single moderately-correlated residue that also displays a threelevel system in its FEL. In dynamical network analysis,^60^ correlations between residues in a protein or protein complex are considered as edge weights between nodes in a graph. The resulting network may then be analysed using the tenets of network theory. All network analyses were conducted using Python libraries Dynetan 1.2.0 and Networkx 2.8.7.^61,62^ Each residue in the protein is represented by one node corresponding to its *α*-carbon atom, while each Chlorophyll *a* is represented by a node corresponding to its Mg atom. The nodes are classified into communities using Louvain heuristics.^63^ The correlation of motion between all the nodes is calculated, and the resulting network can be visualized and analyzed. Due to limited computational resources, four independent network analyses were performed on each protein complex (one per protein sub-unit) for each replica in the simulation.

The Dynamical Network analysis revealed that high correlations (greater than 0.8) were predominantly observed among residues that are adjacent in the protein sequence, indicating a synchronized motion in certain sub-communities of the local environment. Residues with the same name or side chain type did not exhibit specific correlation patterns. (For a more detailed analysis of the correlations among the 55 residues of interest, please see Sec. S4.4 in the Supporting Information.)

We then expanded our analysis to explore correlations between these 55 residues and those located farther away from the pigments to explore potential correlations beyond the 55 identified local residues. Correlations between these distant residues and the nearby ones could indicate their involvement in larger-scale conformational changes. Our analysis found a single distal residue moderately correlated (average correlation of *>* 0.4 across all replicas, cf. Fig. S32) with nine of the local environment residues displaying multimodal distributions:

Threonine (THR) 180. Fig. 8 illustrates the residues of the local environment with which THR 180 is correlated, all of which display multimodal signatures. These residues are not concentrated in the same region of the protein sub-unit but rather are dispersed within the environment of Chlorophyll *a*.

**Figure 8:**
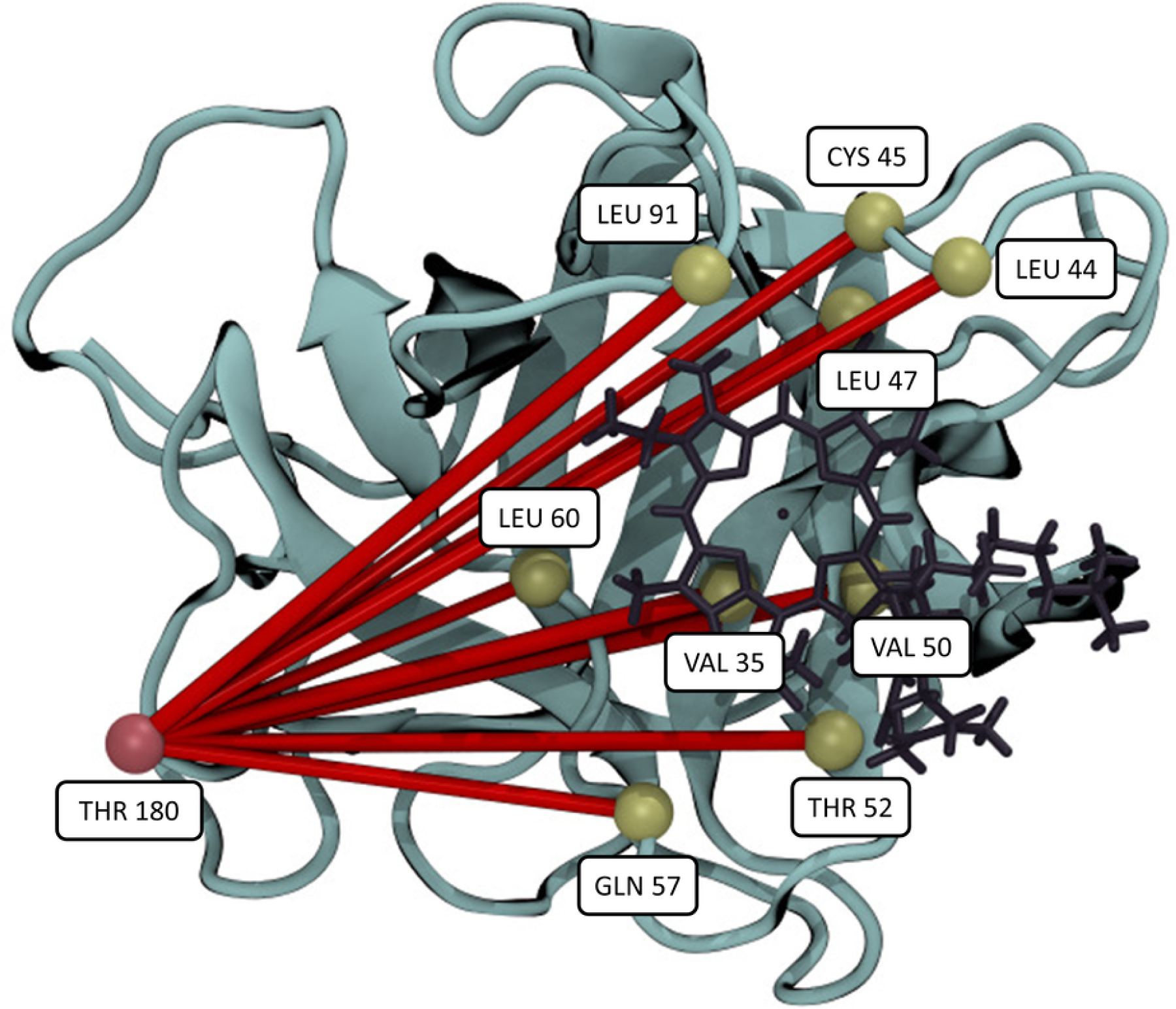
Graphical representations of the dynamical network of THR 180 in protein chain B, averaged over the 5 replicas. The protein chain is colored in blue, the Chlorophyll a in black, the *α* carbon of THR 180 in pink, and the *α* carbon of the identified nearby residues correlated with THR 180 in green. The edges between THR 180 and the other residues are shown in red, and their thicknesses represent their weights.

#### THR 180

The *χ*_1_ angle distribution of THR 180 was computed for all protein chains in all replicas (Fig. S33a-d). All the distributions displayed a trimodal pattern. Furthermore, the average peak positions were found to be approximately −134.0 ± 2.1 *^◦^*, 1.1 ± 0.9 *^◦^*, and 127.8 ± 1.9 *^◦^*, which fall within the same range as the values also reported in Table 3. These results indicate the presence of a similar multi-well conformational change, the rotation of the *χ*_1_ angle, occurring in THR 180, as in the residues with which it is moderately correlated.

Fig. 9 depicts the average one-dimensional protein free energy landscape of THR 180. The associated energy barrier heights (in *cm^−^*^1^) are Δ*F*_1_*_→_*_2_ = 1728 ± 33, Δ*F*_2_*_→_*_3_ = 2094 ± 30, and Δ*F*_1_*_→_*_3_ = 1825 ± 41. Although they are within the same order of magnitude as those found in Section 3.3.2 and in^3–6,64^ these barrier heights are slightly higher. However, based on the observed correlations, it is possible that joint rotations of multiple side chains may be involved in observed experimental signatures.

**Figure 9:**
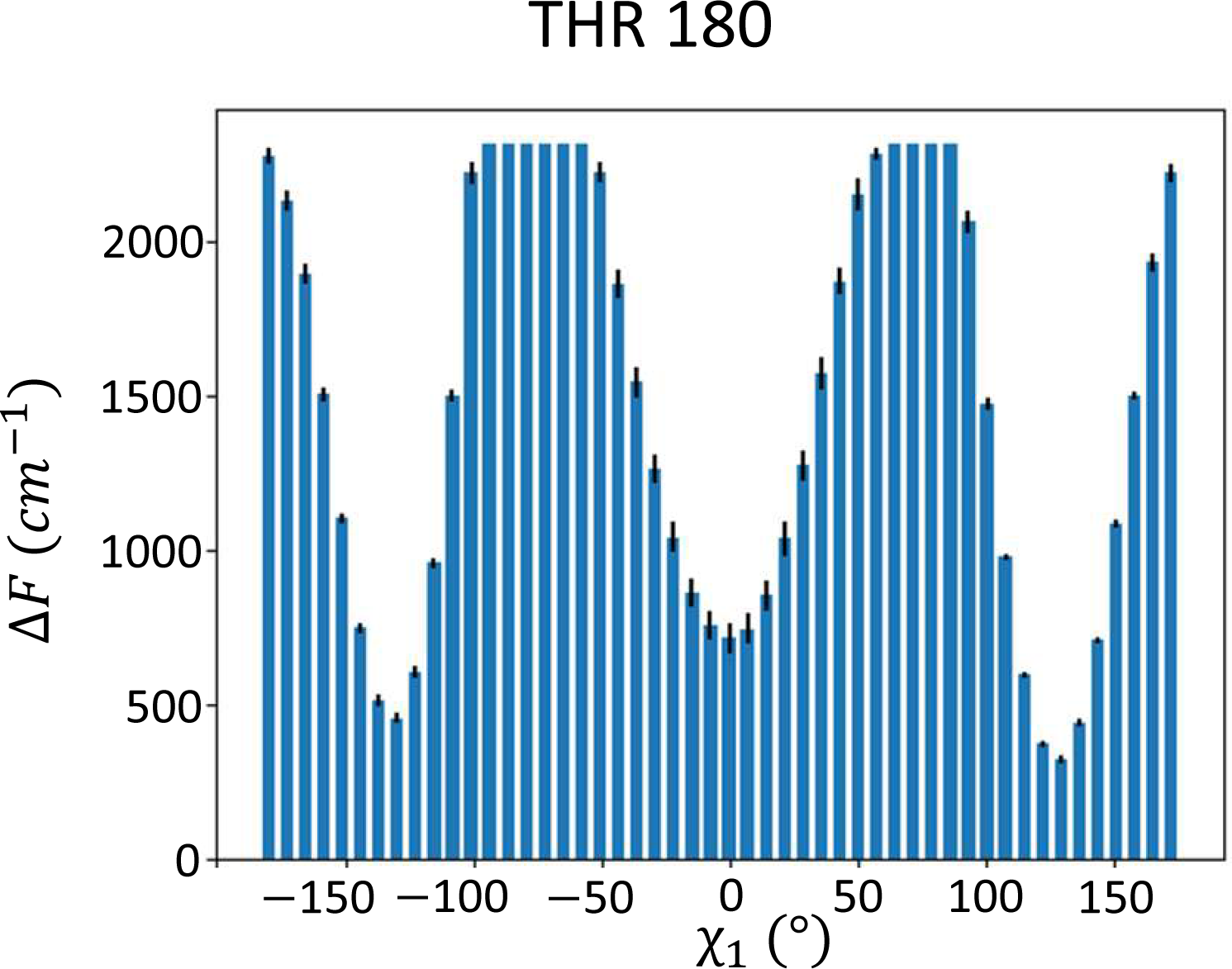
Protein free energy Δ*F* associated with the rotation of the *χ*_1_ angle of distal residue THR 180. Averages and standard errors taken across all protein chains and all replicas.

## 4 Discussion

We have performed molecular dynamics simulations of the Water-Soluble Chlorophyll-a Binding Protein and analysed hydrogen-bonding patterns and local and global residue motions to identify potential molecular-level structural changes responsible for observations of optical line shifts. We found two persistent hydrogen bonds, between the SER53 main chain and the pigment and between the GLN57 side chain and the pigment, that are of the correct length to potentially produce double-well potentials, although the parameters of the respective TLS would have to be determined via quantum-mechanical calculations. Additionally, we found that the GLN57 residue, as well as the THR52 residue, which forms transient long hydrogen bonds, also demonstrate multi-modal potential wells related to their side chain configurations. Although we cannot resolve any free energy barriers along the coordinate of the donor-acceptor distance, and thus identify no classical multi-well potentials, the multimodal states of their side chains might lead to a higher likelihood of donor/acceptor switching or cooperativity between multiple h-bonds and side chain rotations, making these residues of particular interest for high resolution quantum mechanical analysis. We note, also, that the *md*^2^ parameter associated with a *single* hydrogen bond would be at most 2 × 10*^−^*^47^kg-m^2^, too low to account for the NPHB results. Therefore, in future work, it would be best to concentrate on cooperative effects involving multiple hydrogen bonds, perhaps employing multi conformation continuum electrostatics and network analysis with additional MD simulations, similar to previous work on water channel networks in photosystem II.^65^

In total, we found 55 residues forming part of the local environment of the pigments that demonstrate double- and triple-well free energy surfaces with the rotational angles of their side chains serving as the relevant coordinate. Preferred conformations of the *χ*_1_ dihedral angle occur at approximately −130*^◦^*, 1*^◦^*, and 125*^◦^*. Preferred conformations of the *ω*_1_ rotational angle in PRO residues occur at approximately −150*^◦^* and 150*^◦^*. Classical free energy surfaces exhibit barrier heights consistent with those observed in optical experiments.

We also identify a distal residue, THR 180, whose movements are moderately correlated with the multi-well residues of the pigment’s local environment. It also possesses a trimodal distribution of its side chain *χ*_1_, in a similar manner to the residues with which it is correlated.

Taken together, these observations provide support for the TLS/MLS model, as well as offering a number of potential candidates for the origins of the TLS/MLS behavior.

The data presented here should be compared to parameters obtained from the analysis of spectral hole recovery, as the latter is occurring while the pigment is in the ground electronic state. There is a good agreement between calculated barrier heights for rotating side chains and the experimental data.^3–6^ We note, however, that the height of the barrier is just one of the relevant quantities in modeling and understanding the mechanics of NPHB. Analysis of NPHB and hole recovery data also yields another parameter loosely characterizing the size of the entity responsible for the structural change and the extent of its movement, *md*^2^, where *m* is the effective mass of the tunneling entity and *d* is the thickness of the barrier. Note that NPHB and hole recovery data is commonly analyzed using a very simple approximation with rectangular barriers^2,5,6^ with tunneling rate Ω_0_*e^−^*^2^*^λ^*, where Ω_0_ is the attempt frequency and *λ* is the tunneling parameter:^66^

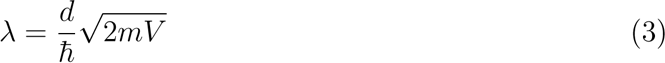

Here *V* is the height of the barrier. In this context, *d* has dimensions of length. Note that larger *λ* corresponds to lower tunneling / NPHB / recovery rates. Adapting similar rectangular-barrier logic to an angle-type generalized coordinate yields

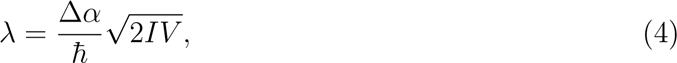

where Δ*α* is an angular barrier thickness in radians and *I* is the moment of inertia. This analogy relies on the similarity of the wavefunctions for a free particle and a quantum mechanical rotor. Both are complex exponentials, although periodic boundary conditions result in quantization of energy in the case of the rotor even in the absence of any potential energy term. However, the barriers depicted in Figures 6 and 7 are not rectangular. Employing the WKB approximation one can demonstrate that for inverse parabolic barriers and energies far enough from the barrier tops,

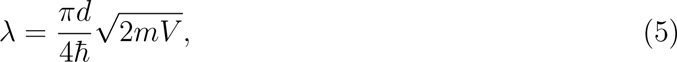

A 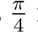 factor reduction that would likely also be preserved in an angular formulation. Although we do not have the means yet to do a fully quantitative comparison, we may make a few simplifying assumptions to compare computed moments of inertia in our system (cf. Table 4) with *md*^2^ values obtained from NPHB experiments, i.e. 1.0-4.0 × 10*^−^*^46^ kg-m^2^.^3–6^ The moments of inertia of the side groups we identified as corresponding to multi-modal angular distributions at 300 K range from 0.5 × 10*^−^*^46^ kg-m^2^ for ALA and 3.3 × 10*^−^*^46^ kg-m^2^ for SER up to 75 × 10*^−^*^46^ kg-m^2^ for TYR. Assuming the angular barrier thickness 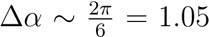 radians, one can directly compare these numerical values to the experimental ones noted above.

Based on these arguments, SER and ALA residues display FELs and moments of inertia most in accord with observed experimental values. However, the moment of inertia for ALA is too small and furthermore the rotation of a methyl group by 120 degrees is known to yield a very specific NPHB signature^67,68^ that has not been observed in pigment-protein complexes so far. In addition, with respect to SER one could note that the wells in Fig 6d have similar depths, which makes tunneling at low temperatures much more feasible than in the case of very different depths calculated, for example, for CYS, VAL, LEU or THR.

For most of the other amino acids, the data presented in Table 4 would yield *λ* values which are significantly (order of magnitude) larger than deduced from spectroscopy experiments. In other words, other hole burning and recovery channels, if present, would out-compete the NPHB involving rotation of most of the above side groups and the latter mechanism would not be the dominant one, making the SER residues of greatest interest for further study.

Further improvement in terms of comparing MD and spectroscopic data could be achieved by introducing more rigorous quantum-mechanical modeling of tunneling between the wells of the landscape, including realistic energy levels, stationary-state wavefunctions, attempt frequencies and tunneling probabilities.^69^ Although simple models employed for simulating spectroscopy data feature temperature-independent barriers (only transition rates are temperature-dependent),^5,6^ it is possible that barrier heights do depend on temperature. The magnitudes of the spectral shifts, another parameter that could be compared between simulations and experiments, would depend on the distance between the rotating side-group and the pigment molecule and their mutual orientation. Determination of transition energies of the pigments embedded in the protein is impossible from MD simulations alone and requires DFT / QM modeling. In conjunction with improving the theory underlying the spectroscopic data, now that exploratory unbiased simulations have been performed and motions of interest identified, the use of enhanced sampling molecular dynamics methods can be used to further explore and refine the energy landscapes.

## 5 Conclusions

Overall, the investigation of the local environment of the pigments in WSCP led to the identification of amino acid side-group rotation as a particularly feasible mechanism responsible for spectral dynamics. This mechanism appears fairly universal and not specific to one particular amino acid. The heights of the energy barriers of the residues displaying multimodal distributions were calculated and the majority appeared to fall within a similar range of values as the (low-temperature) experimental data. Once the size of the side-groups is taken into account, the best qualitative and quantitative agreement is achieved for SER.

The next steps in terms of simulations will be to investigate this pool of candidates at a more refined level of modeling, either via density functional calculations or through quantum mechanical-molecular mechanical ones to assess whether the suggested hydrogen bonds demonstrate a double-well landscape and further to examine potential excited-state energy barriers to fully characterize the dual-TLS model.

Furthermore, the energy landscapes obtained from our simulations can be utilized to model the system evolution incorporating Quantum Mechanical tunneling. Approaches described in Najafi and Zazubovich ^64^ and Garashchuk et al. ^69^ can be employed for this purpose. Additionally, with the identified generalized coordinate, it would be beneficial to examine and compare the rates of conformational changes between different residues, investigating potential differences or similarities among them. Moreover, conducting simulations on other pigment-protein complexes such as Cytochrome b_6_f or the LH 2 antenna complex of purple bacteria could provide insights into whether similar small conformational changes are observed in these systems. Notably, high-quality non-photochemical hole burning data focusing explicitly on spectral dynamics are already available for these two complexes, enabling direct comparisons. The next obvious step in terms of experiments would be to subject WSCP to as detailed NPHB and recovery experiments and respective analysis as have been employed so far for CP43,^3^ cytochrome b6f^4–6^ and LH2 (^2^ and work in progress).

This article provides a stepping stone in bridging the gap between theory and experiments in terms of phenomena observed in NPHB and single-complex spectroscopy experiments in the field of photosynthesis. A basic science understanding of the relationship between theory and experiments in optical spectroscopy of pigment-protein complexes leads us nearer to a comprehensive understanding of the primary processes of photosynthesis. Such an understanding is crucial, as it not only enables the development of renewable energy solutions but also enhances our understanding of fundamental biological processes.

## Supporting Information

PDF file “SI.pdf” containing supporting text (four sections) and supporting figures (33). Zip file “params.zip” containing parameters for nonstandard residues. Zip file “inputfiles.zip” containing input files for simulation.

## Supporting information

input files for MD

force field parameters

## Acknowledgement

The authors thank Dr. Laszlo Kalman and Dr. Deniz Meneksedag-Erol of Concordia University for useful feedback. This research was enabled in part by support provided by Calcul Quebec (www.calculquebec.ca) and the Digital Research Alliance of Canada (https://alliancecan.ca). Support by Concordia University via Chair Research Funds is acknowledged by VZ. This research was undertaken, in part, thanks to funding from the Canada Research Chairs Program under grant number CRC-2020-00225 (RM). This research was supported in part by Discovery Grant #RGPIN-2021-03470 from the National Sciences and Engineering Research Council of Canada (RM).

## TOC Graphic

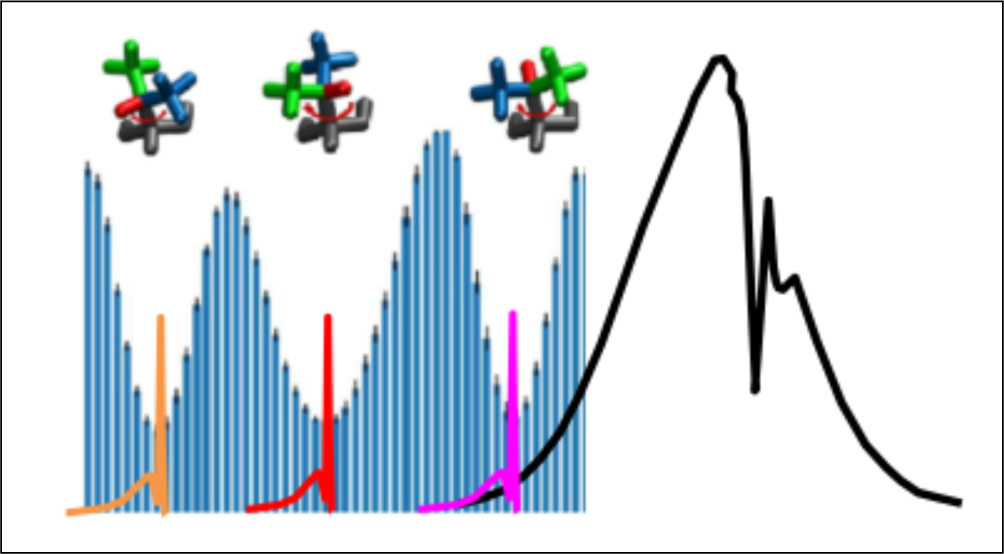

