## supplemental information for "Identification of residues potentially involved in optical shifts in the water-soluble chlorophyll-a binding protein through molecular dynamics simulations"

### S1 PropKa Results

1 10 20 30 40
I N D E E P V K D T N G N P L K I E T R Y F I Q P A S D N N G G G L V P A N V D L S H L C

50 60 70 80 90
P L G I V R T S L P Y Q P G L P V T I S T P S S S E G N D V L T N T N I A I T F D A P I W

100 110 120 130
L C P S S K T W T V D S S S E E K Y I I T G G D P K S G E S F F R I E K Y G N G K N T Y K

140 150 160 170 180
L V R Y D N G E G K S V G S T K S L W G P A L V L N D D D D S D E N A F P I K F R E V D T

Figure S1: Protein sequence of one protein chain of the homotetramer Water-soluble chlorophyll protein, from *Lepidium virginicum*, PDB ID : 2DRE. Acidic amino acids Glumatic acid (E), and Aspartic acid (D) are colored in blue, and are considered deprotonated (negatively charged). Basic amino acids Lysine (K), Arginine (R), and Histidine (H) are colored in red, and are considered protonated (positively charged).

| RESIDUE | pKa | pKmodel |
|---|---|---|
| ASP 3A | 4.02 | 3.80 |
| ASP 9A | 2.73 | 3.80 |
| ASP 28A | 3.80 | 3.80 |
| ASP 40A | 2.38 | 3.80 |
| ASP 74A | 3.10 | 3.80 |
| ASP 86A | 3.80 | 3.80 |
| ASP 101A | 3.31 | 3.80 |
| ASP 114A | 3.11 | 3.80 |
| ASP 140A | 3.15 | 3.80 |
| ASP 162A | 3.87 | 3.80 |
| ASP 163A | 4.15 | 3.80 |
| ASP 164A | 3.69 | 3.80 |
| ASP 165A | 2.37 | 3.80 |
| ASP 167A | 4.01 | 3.80 |
| ASP 179A | 3.29 | 3.80 |
| ASP 3B | 2.71 | 3.80 |
| ASP 9B | 2.84 | 3.80 |
| ASP 28B | 3.80 | 3.80 |
| ASP 40B | 2.30 | 3.80 |
| ASP 74B | 3.02 | 3.80 |
| ASP 86B | 3.15 | 3.80 |
| ASP 101B | 3.51 | 3.80 |
| ASP 114B | 2.82 | 3.80 |
| ASP 140B | 3.87 | 3.80 |
| ASP 162B | 3.80 | 3.80 |
| ASP 163B | 3.99 | 3.80 |
| ASP 164B | 4.15 | 3.80 |
| ASP 165B | 0.13 | 3.80 |
| ASP 167B | 3.31 | 3.80 |
| ASP 179B | 3.87 | 3.80 |
| ASP 3C | 3.38 | 3.80 |
| ASP 9C | 2.61 | 3.80 |
| ASP 28C | 3.10 | 3.80 |
| ASP 40C | 2.35 | 3.80 |
| ASP 74C | 3.07 | 3.80 |
| ASP 86C | 3.44 | 3.80 |
| ASP 101C | 2.25 | 3.80 |
| ASP 114C | 2.80 | 3.80 |
| ASP 140C | 3.99 | 3.80 |
| ASP 162C | 3.80 | 3.80 |
| ASP 163C | 4.06 | 3.80 |
| ASP 164C | 4.01 | 3.80 |
| ASP 165C | 3.60 | 3.80 |
| ASP 167C | 2.73 | 3.80 |
| ASP 179C | 4.09 | 3.80 |
| ASP 3D | 4.09 | 3.80 |
| ASP 9D | 2.63 | 3.80 |
| ASP 28D | 3.32 | 3.80 |
| ASP 40D | 2.44 | 3.80 |
| ASP 74D | 3.80 | 3.80 |
| ASP 86D | 3.42 | 3.80 |
| ASP 101D | 3.03 | 3.80 |
| ASP 114D | 2.31 | 3.80 |
| ASP 140D | 2.66 | 3.80 |
| ASP 162D | 3.80 | 3.80 |
| ASP 163D | 2.57 | 3.80 |
| ASP 164D | -0.30 | 3.80 |
| ASP 165D | 4.56 | 3.80 |
| ASP 167D | 2.86 | 3.80 |
| ASP 179D | 3.87 | 3.80 |

(a) Estimations of the pKa values for the acidic ASP residues in the homotetramer Water-soluble chlorophyll protein, from *Lepidium virginicum*, PDB ID : 2DRE, obtained from the PropKa website. The first column lists the residues, the second column the estimations of the pKa values, and the third column the theoretical pKa values.

| RESIDUE | pKa | pKmodel |
|---|---|---|
| GLU 4A | 4.64 | 4.50 |
| GLU 5A | 4.01 | 4.50 |
| GLU 18A | 4.44 | 4.50 |
| GLU 71A | 4.36 | 4.50 |
| GLU 105A | 4.50 | 4.50 |
| GLU 106A | 3.56 | 4.50 |
| GLU 119A | 3.99 | 4.50 |
| GLU 125A | 2.60 | 4.50 |
| GLU 143A | 3.78 | 4.50 |
| GLU 168A | 5.39 | 4.50 |
| GLU 177A | 4.22 | 4.50 |
| GLU 4B | 4.71 | 4.50 |
| GLU 5B | 3.83 | 4.50 |
| GLU 18B | 4.69 | 4.50 |
| GLU 71B | 4.64 | 4.50 |
| GLU 105B | 4.89 | 4.50 |
| GLU 106B | 3.86 | 4.50 |
| GLU 119B | 4.22 | 4.50 |
| GLU 125B | 8.42 | 4.50 |
| GLU 143B | 4.58 | 4.50 |
| GLU 168B | 4.50 | 4.50 |
| GLU 177B | 4.57 | 4.50 |
| GLU 4C | 4.50 | 4.50 |
| GLU 5C | 3.98 | 4.50 |
| GLU 18C | 4.57 | 4.50 |
| GLU 71C | 3.73 | 4.50 |
| GLU 105C | 4.50 | 4.50 |
| GLU 106C | 3.44 | 4.50 |
| GLU 119C | 3.78 | 4.50 |
| GLU 125C | 3.27 | 4.50 |
| GLU 143C | 4.50 | 4.50 |
| GLU 168C | 4.50 | 4.50 |
| GLU 177C | 4.61 | 4.50 |
| GLU 4D | 4.50 | 4.50 |
| GLU 5D | 4.10 | 4.50 |
| GLU 18D | 4.50 | 4.50 |
| GLU 71D | 4.45 | 4.50 |
| GLU 105D | 3.54 | 4.50 |
| GLU 106D | 3.98 | 4.50 |
| GLU 119D | 4.64 | 4.50 |
| GLU 125D | 7.43 | 4.50 |
| GLU 143D | 4.64 | 4.50 |
| GLU 168D | 4.50 | 4.50 |
| GLU 117D | 4.57 | 4.50 |

(b) Estimations of the pKa values for the acidic GLU residues in the homotetramer Water-soluble chlorophyll protein, from *Lepidium virginicum*, PDB ID : 2DRE, obtained from the PropKa website. The first column lists the residues, the second column the estimations of the pKa values, and the third column the theoretical pKa values. GLU 125 in protein chains B and D (in blue) were kept deprotonated in the simulations but could have been kept protonated at pH = 7 based on the estimated pKa values.

| RESIDUE | pKa | pKmodel | RESIDUE | pKa | pKmodel | RESIDUE | pKa | pKmodel |
|---|---|---|---|---|---|---|---|---|
| HIS 43A | 7.04 | 6.50 | LYS 145B | 9.94 | 10.50 | LYS 145D | 10.08 | 10.50 |
| HIS 43B | 6.50 | 6.50 | LYS 151B | 10.50 | 10.50 | LYS 151D | 10.36 | 10.50 |
| HIS 43C | 7.09 | 6.50 | LYS 174B | 10.29 | 10.50 | LYS 174D | 10.43 | 10.50 |
| HIS 43D | 6.38 | 6.50 | LYS 8C | 10.15 | 10.50 | ARG 20A | 12.08 | 12.50 |
| LYS 16A | 10.15 | 10.50 | LYS 16C | 10.36 | 10.50 | ARG 51A | 11.87 | 12.50 |
| LYS 96A | 10.29 | 10.50 | LYS 96C | 10.29 | 10.50 | ARG 123A | 11.87 | 12.50 |
| LYS 107A | 10.22 | 10.50 | LYS 107C | 10.29 | 10.50 | ARG 138A | 11.56 | 12.50 |
| LYS 116A | 10.50 | 10.50 | LYS 116C | 10.50 | 10.50 | ARG 176A | 11.60 | 12.50 |
| LYS 126A | 10.08 | 10.50 | LYS 126C | 9.87 | 10.50 | ARG 20B | 12.01 | 12.50 |
| LYS 131A | 10.50 | 10.50 | LYS 131C | 10.50 | 10.50 | ARG 51B | 11.43 | 12.50 |
| LYS 135A | 12.18 | 10.50 | LYS 135C | 11.10 | 10.50 | ARG 123B | 11.24 | 12.50 |
| LYS 145A | 10.29 | 10.50 | LYS 145C | 10.29 | 10.50 | ARG 138B | 11.66 | 12.50 |
| LYS 151A | 9.80 | 10.50 | LYS 151C | 10.29 | 10.50 | ARG 176B | 12.20 | 12.50 |
| LYS 174A | 8.57 | 10.50 | LYS 174C | 10.36 | 10.50 | ARG 20C | 12.08 | 12.50 |
| LYS 8B | 10.36 | 10.50 | LYS 8D | 10.43 | 10.50 | ARG 51C | 11.74 | 12.50 |
| LYS 16B | 10.15 | 10.50 | LYS 16D | 10.29 | 10.50 | ARG 123C | 11.66 | 12.50 |
| LYS 96B | 9.45 | 10.50 | LYS 96D | 10.01 | 10.50 | ARG 138C | 11.72 | 12.50 |
| LYS 107B | 10.36 | 10.50 | LYS 107D | 10.43 | 10.50 | ARG 176C | 11.55 | 12.50 |
| LYS 116B | 10.29 | 10.50 | LYS 116D | 10.29 | 10.50 | ARG 20D | 12.01 | 12.50 |
| LYS 126B | 10.08 | 10.50 | LYS 126D | 9.94 | 10.50 | ARG 51D | 11.09 | 12.50 |
| LYS 131B | 10.50 | 10.50 | LYS 131D | 10.50 | 10.50 | ARG 123D | 11.73 | 12.50 |
| LYS 135B | 13.57 | 10.50 | LYS 135D | 13.87 | 10.50 | ARG 138D | 10.46 | 12.50 |
| | | | | | | ARG 176D | 11.80 | 12.50 |

(c) Estimations of the pKa values for the basic HIS, LYS, and ARG residues in the homotetramer Water-soluble chlorophyll protein, from *Lepidium virginicum*, PDB ID : 2DRE, obtained from the PropKa website. The first column lists the residues, the second column the estimations of the pKa values, and the third column the theoretical pKa values. HIS 43 in protein chains B and D (in red) were kept protonated in the simulations but could have been kept deprotonated at pH = 7 based on the estimated pKa values.

Figure S2: Estimations of the pKa values for the acidic ASP and GLU residues, and the basic HIS, LYS, and ARG residues in the homotetramer Water-soluble chlorophyll protein, from *Lepidium virginicum*, PDB ID : 2DRE, obtained from the PropKa website. The first column lists the residues, the second column the estimations of the pKa values, and the third column the theoretical pKa values.

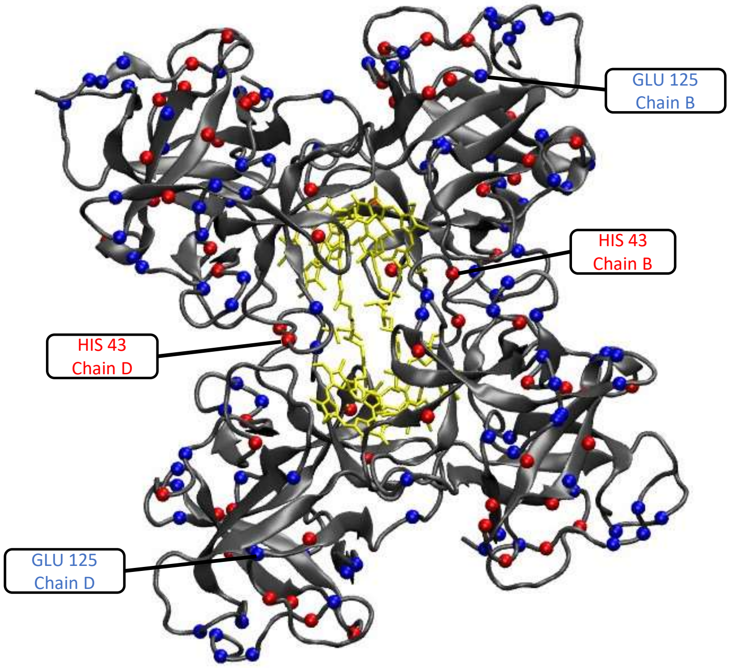


Figure S3: Crystral structure of the homotetramer Water-soluble chlorophyll protein, from *Lepidium virginicum*, PDB ID : 2DRE. Chlorophylls a are colored in yellow, and the four protein chains in grey. Acidic amino acids are colored in blue and are considered negatively charged, basic amino acids are colored in red and are considered positively charged.

#### S2 Local environment identification

To assess the relevance of these 168 residues in a dynamic context, we computed the contacts between the pigments and all residues with the Contact Map Explorer Python package.[S1]

To ensure the inclusion of closely associated residues, an all-atom cut-off distance of 3.0 Å was set between the Chlorophyll *a* and the residues. As the exact definition of "close environment" was somewhat arbitrary, the choice of the cut-off distance was made based on the approximate expected hydrogen bond lengths (Section 3.2). Our contact analysis identified 31 residues per chain as having an occurrence frequency of greater than 0.4, all of which were listed as part of the hydrophobic cavity (Fig. S4). The eleven per protein chain that were identified as part of the hydrophobic cavity but had a smaller occurrence frequency than 0.4 were VAL39, ASP40, CYS45, PRO46, PRO55, TYR56, ASP86, SER95, LYS96, SER152, and TPR154. We decided to use the less strict definition of the environment associated with the hydrophobic cavity and include the above 11 residues in our definition of the "local environment".

The above residues in the pigments' local environment can be categorized based on the type of their side chains into polar, non-polar, basic, and acidic (Fig. S4b). The prevalence of non-polar residues is not surprising, as these are the hydrophobic residues that are expected to contribute to the formation of hydrophobic cavities around Chls *a*. The small percentages of basic and acidic residues are also as anticipated, as these types of residues are hydrophilic and their respective positively and negatively charged side chains are typically engaged in hydrogen bonding with water molecules. We can see that among the 168 selected residues, the most abundant residue types are the hydrophobic leucine (LEU), with 28 residues, and the hydrophobic proline (PRO) with 28 residues as well. This is in line with the high contribution of LEU and PRO in the stabilization of the hydrophobic environment and may play a role in facilitating specific protein interactions within that region.

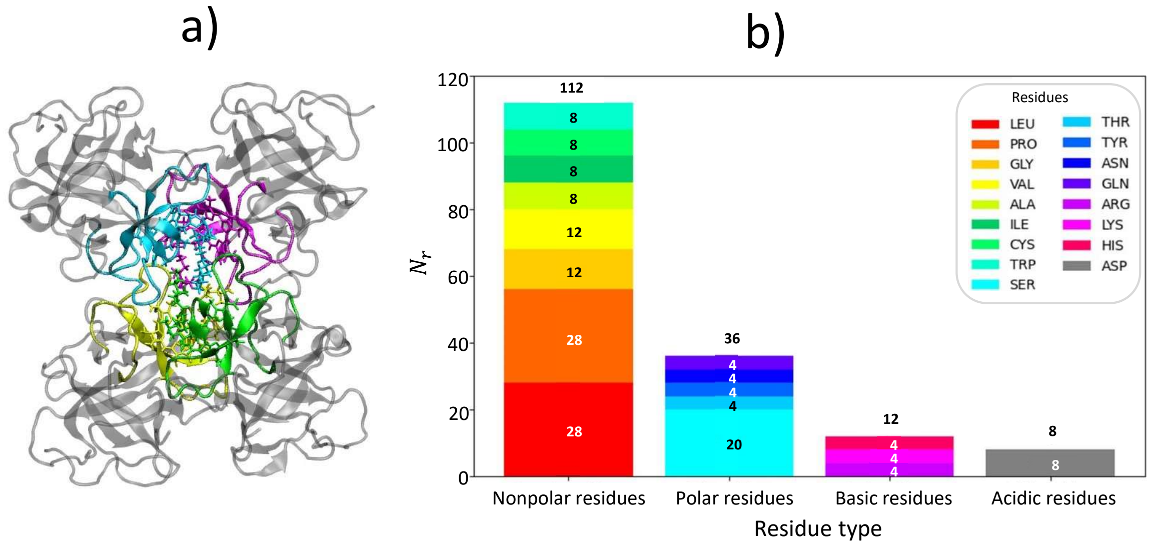


Figure S4: Composition of local environment. (a) Graphical representation of the studied WSCP complex, with the local environment highlighted. Monomers are distinguished by color; pigments are shown in ball-and-stick representation which the rest of the complex is shown in terms of secondary structure. (b) Residue count ($N_r$) by type and amino acid comprising the full local environment of all four monomers. All visualization of pigment-protein complexes was performed using VMD 1.9.4a43.[S2]

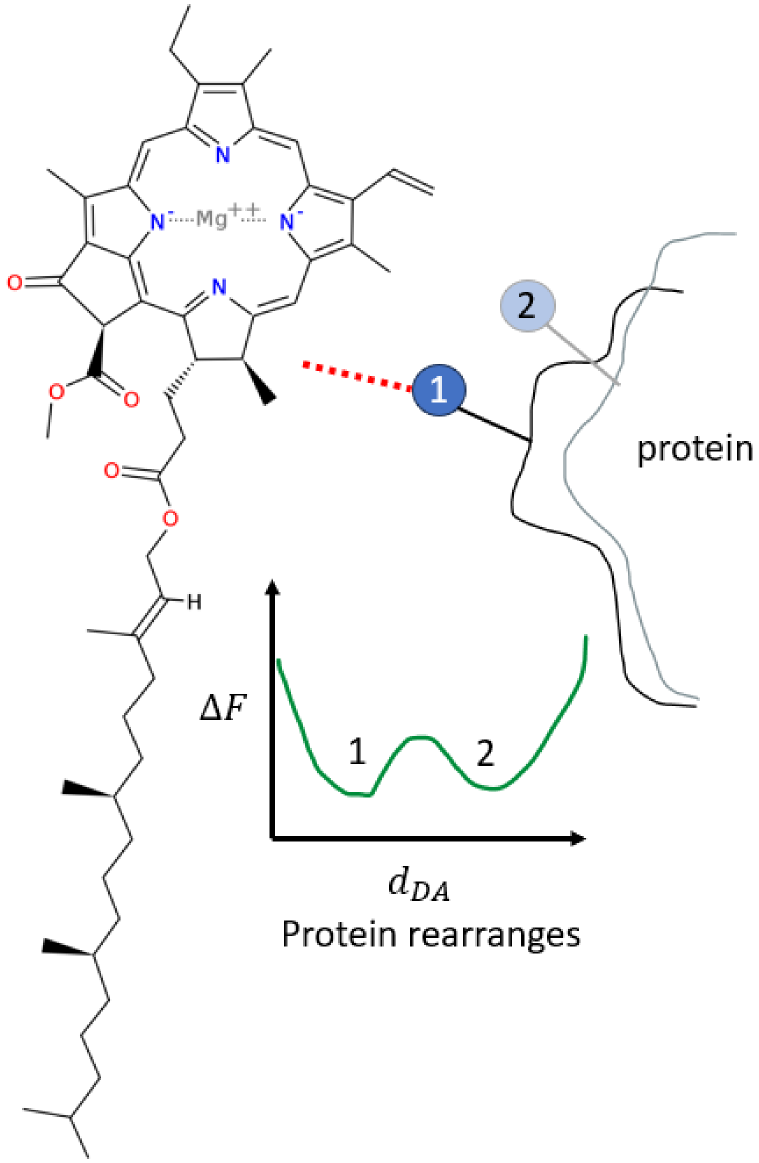


Figure S5: Schematic illustration of possible situation responsible for optical shifts involving long hydrogen bonds. The blue atom functions as a hydrogen bond donor interacting with an acceptor on Chl *a*. Due to structural rearrangements of the protein, this results in two distinct distances between donor and acceptor $d_{DA}$, separated by a free energy barrier in $\Delta F$. In state 1, a hydrogen bond forms; in state 2, it does not.

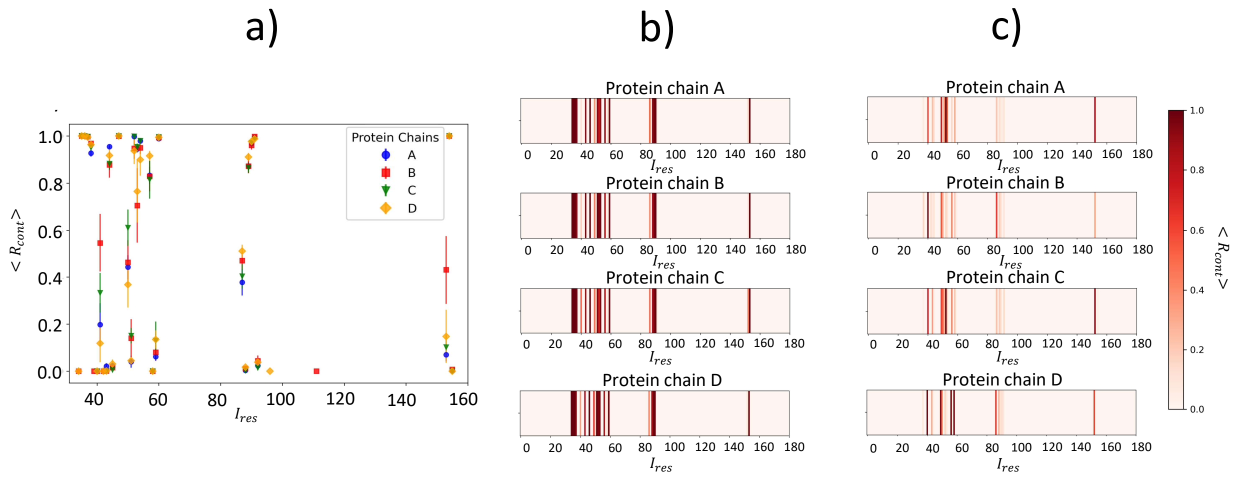


Figure S6: Contacts from which local environments were defined. (a) Frequency of contact occurrence $\langle R_{\text{cont}} \rangle$ versus residue index $I_{\text{res}}$ for the four monomers. b) Heatmaps of mean contact occurrence $\langle R_{\text{cont}} \rangle$ in separate chains. The standard error computed from five independent replicas is shown in (c).

### S3 Analysis of hydrogen bonding

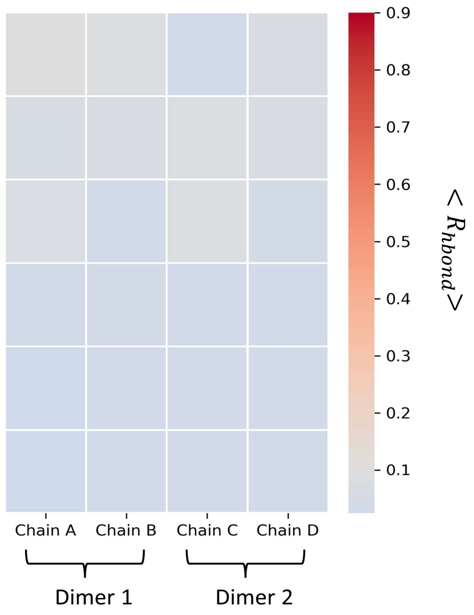


Figure S7: Standard error of average occurrence $\langle R_{\mathrm{hbond}} \rangle$ taken across five independent replicas of long hydrogen bonds. The average occurrence is shown in Fig. 2

### S4 Multimodal distributions associated with side chain angle dihedrals of certain residues

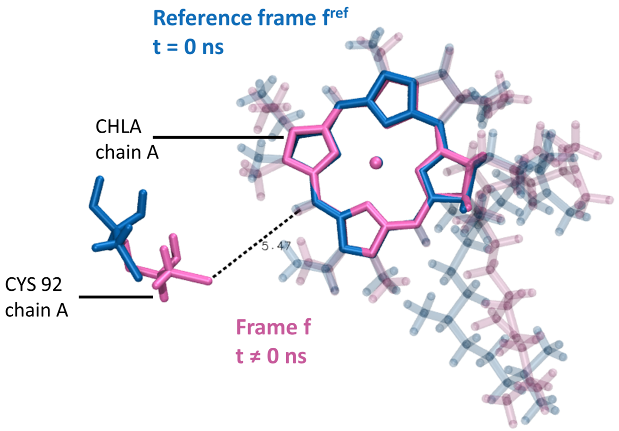


Figure S8: Graphical representation of the RMSD calculation of residue CYS 92 in chain A. The reference frame, $f^{ref}$, is colored in blue, and the frame used for calculation, $f$, is in pink. The fit is made on the Chl $a$ phorphyrin ring allowing calculation of the deviation in the residue position with respect to Chl $a$ phorphyrin ring. The figure was rendered using VMD 1.9.4a43.

#### S4.1 Specifics of dihedral angle calculations for VAL, LEU

Dihedral angles represent the orientational relationship between four consecutive atoms along a chain. They describe the rotation around a bond connecting two of the four atoms, while the other two atoms define the plane in which the rotation occurs. In the case of $\chi_1$ dihedral angle, the atoms involved are the N and CA atoms of the backbone, and two atoms specific to the side chain of the residue. For VAL, the $\chi_1$ angle is defined by atoms N-CA-CB-CG1, as shown in Figure 4a in the main text, with CB being the $\beta-$carbon, the first carbon atom in the VAL side chain, and CG1 being the $\gamma_1-$carbon, the second carbon atom in the VAL side chain. For LEU, the $\chi_1$ angle is defined by atoms N-CA-CB-CG, as shown in Figure

4b in the main text, with CG being the $\gamma-$carbon, the second carbon atom in the LEU side chain. $\chi_1$ angle is determined similarly for other amino acid residues.

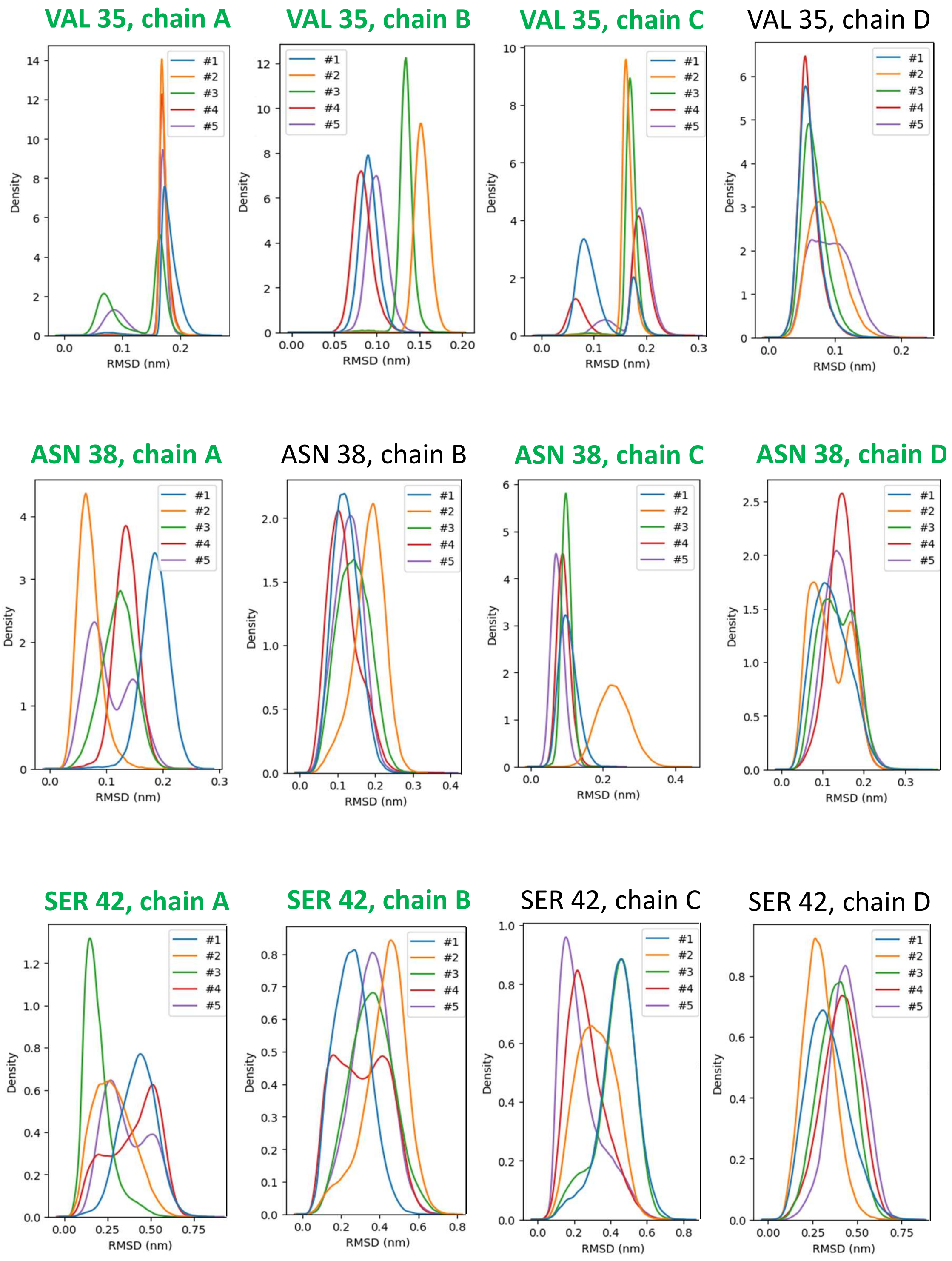


Figure S9: RMSD distributions of residues demonstrating bimodal distributions. Green highlights denote observed multimodal distributions. (Additional distributions presented in Figs S10-S16)

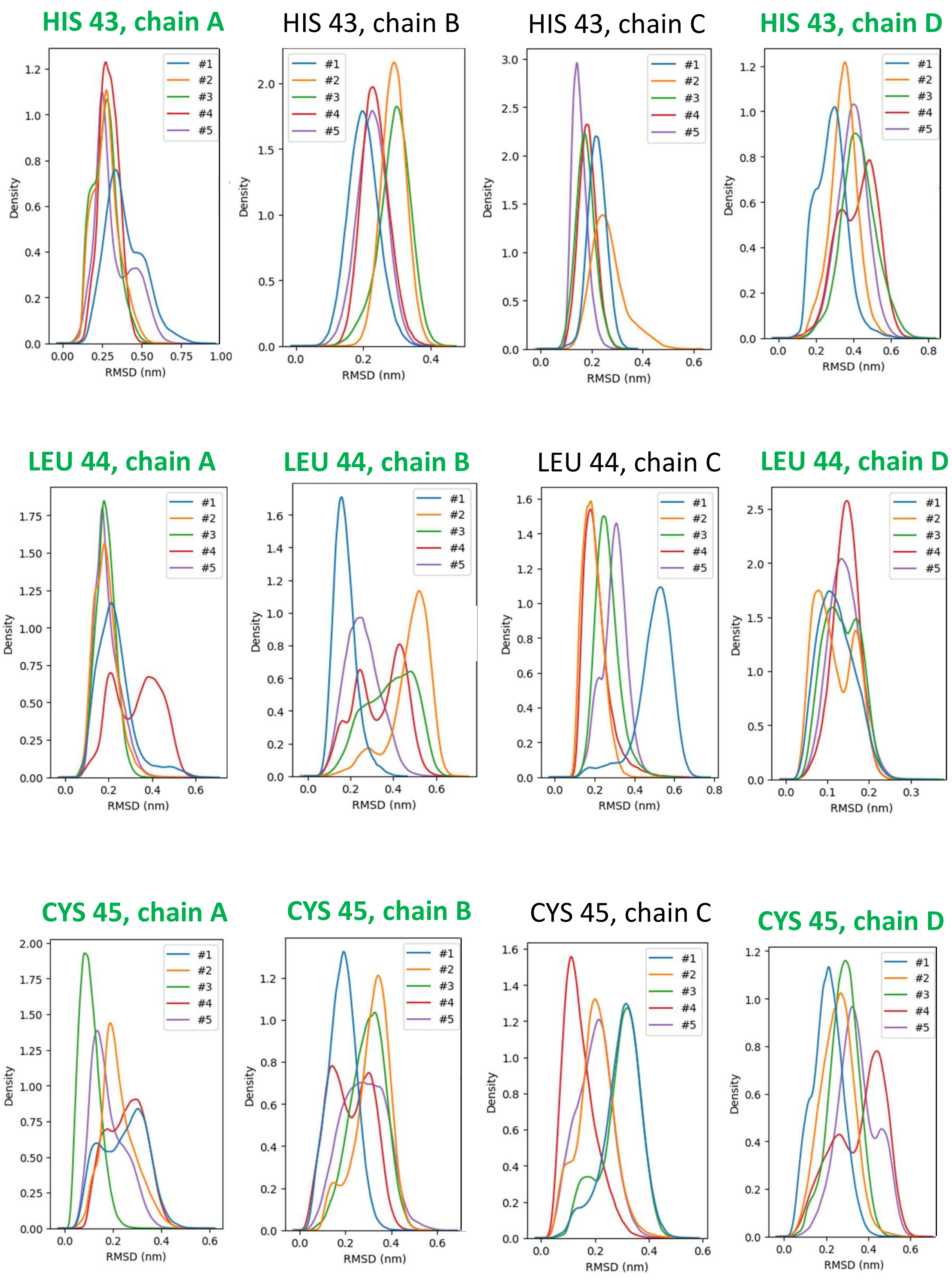


Figure S10: RMSD distributions of residues demonstrating bimodal distributions, continued. (Additional distributions presented in Figs S9, S11-S16)

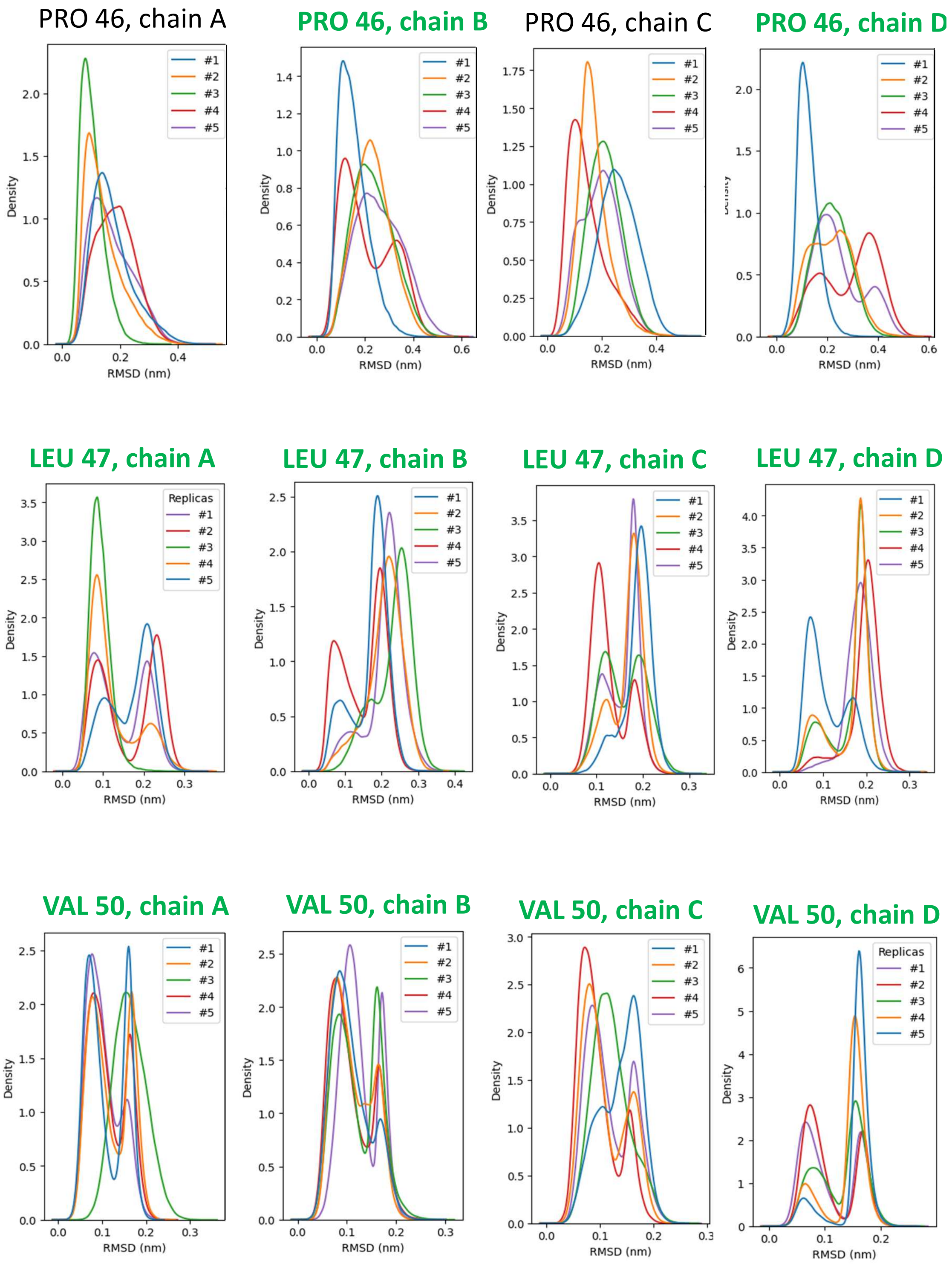


Figure S11: RMSD distributions of residues demonstrating bimodal distributions, continued. (Additional distributions presented in Figs S9-S10, S12-S16)

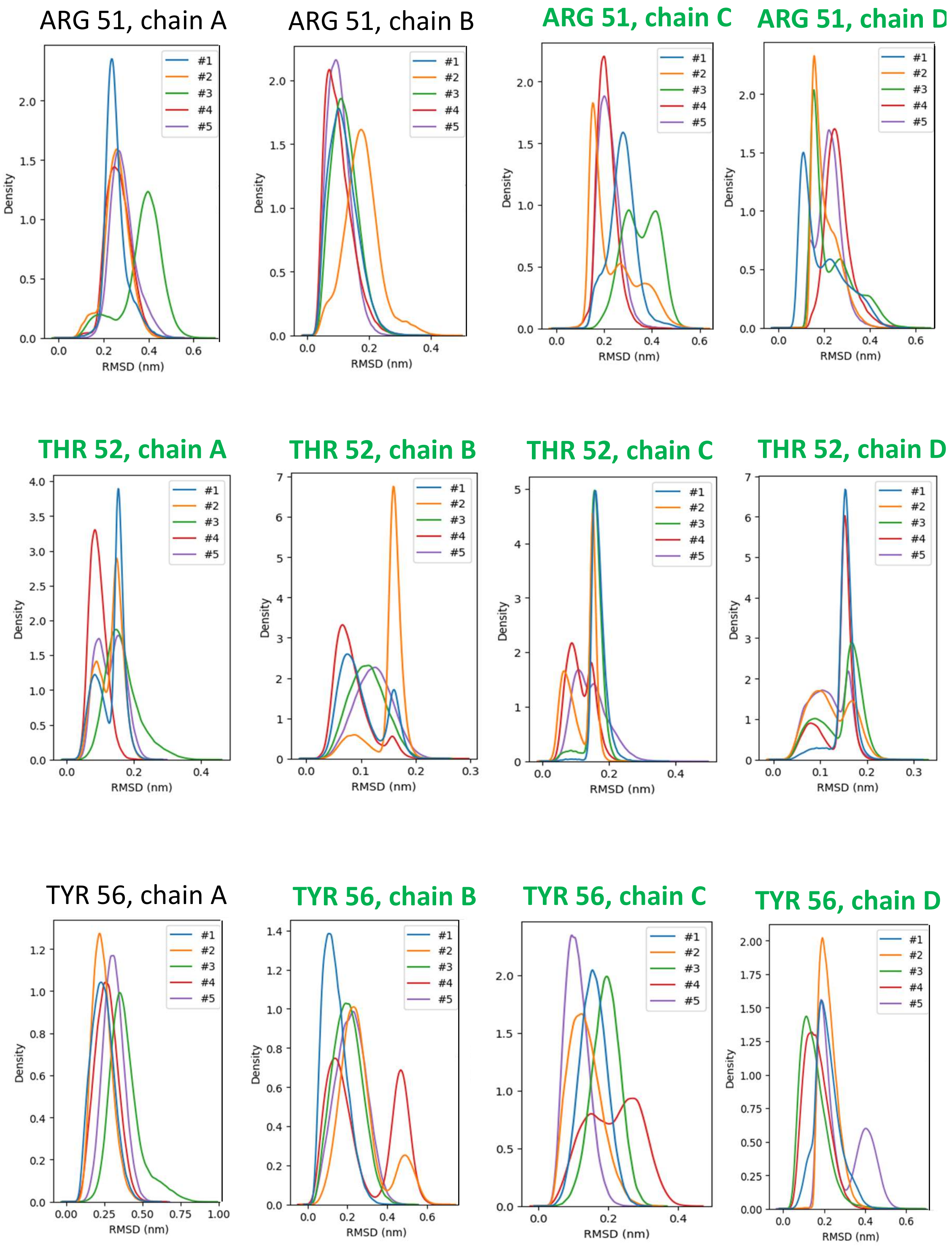


Figure S12: RMSD distributions of residues demonstrating bimodal distributions, continued. (Additional distributions presented in Figs S9-S11,S13-S16)

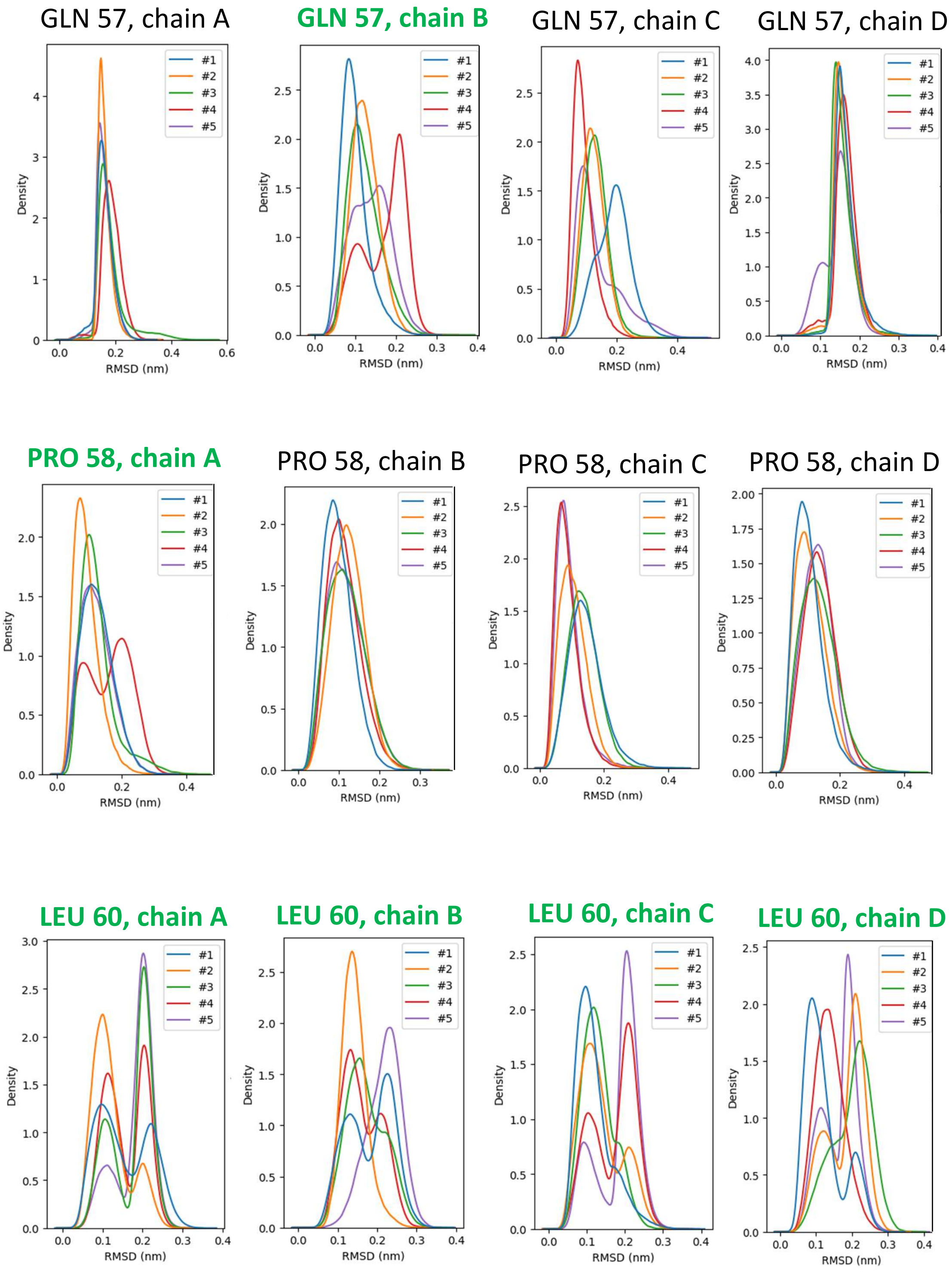


Figure S13: RMSD distributions of residues demonstrating bimodal distributions, continued. (Additional distributions presented in Figs S9-S12,S14-S16)

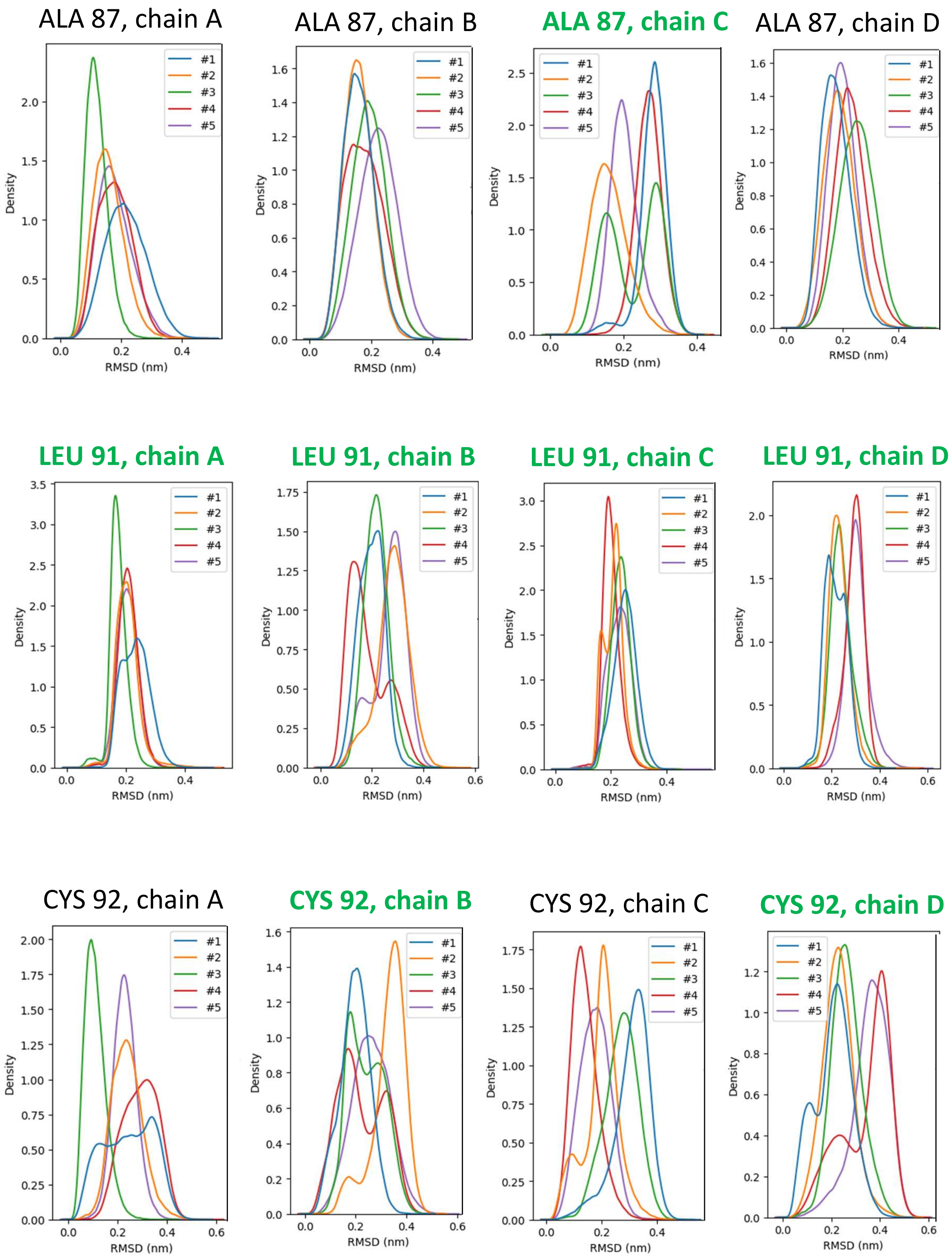


Figure S14: RMSD distributions of residues demonstrating bimodal distributions, continued. (Additional distributions presented in Figs S9-S13,S15-S16)

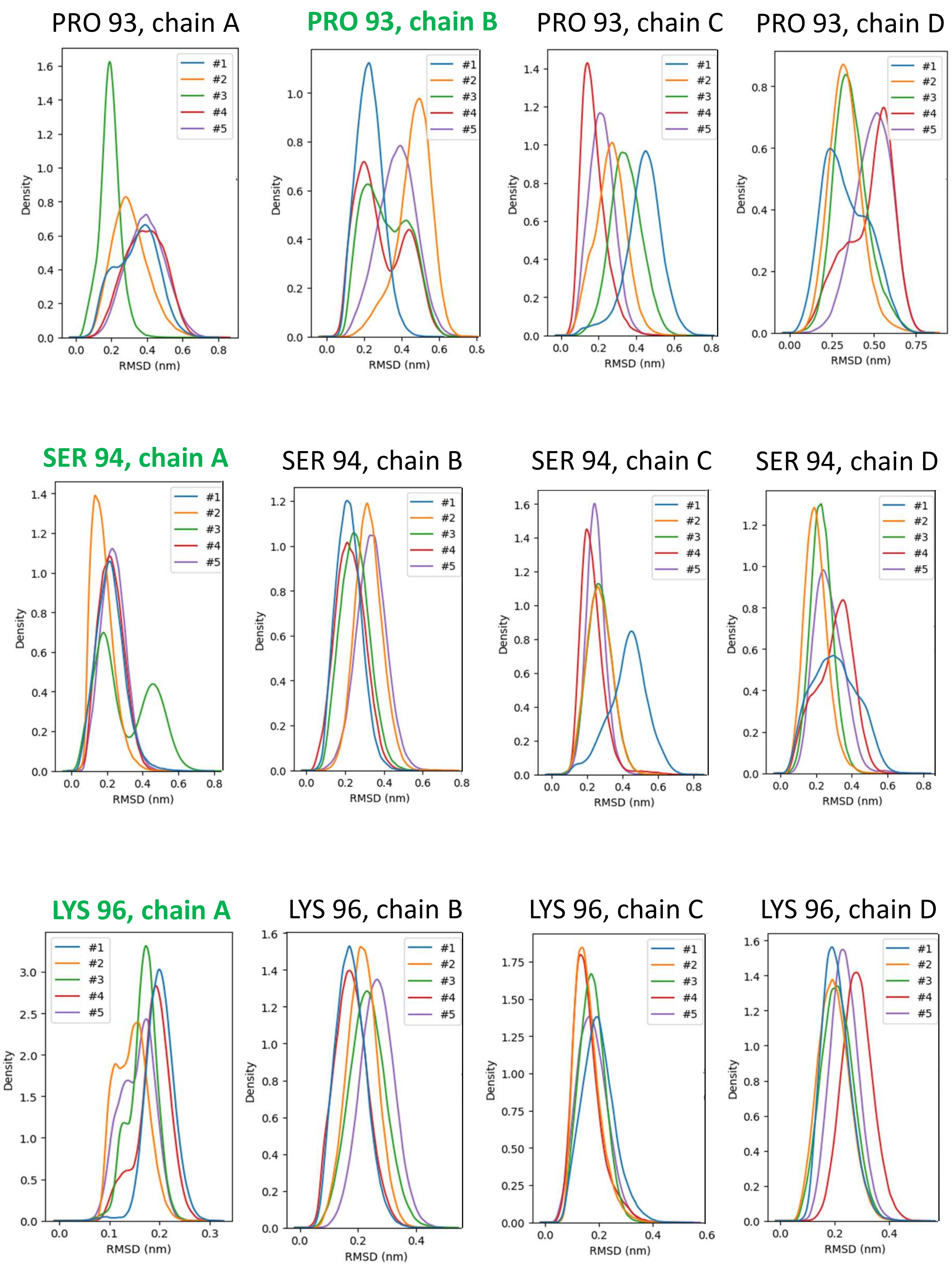


Figure S15: RMSD distributions of residues demonstrating bimodal distributions, continued. (Additional distributions presented in Figs S9-S14,S16)

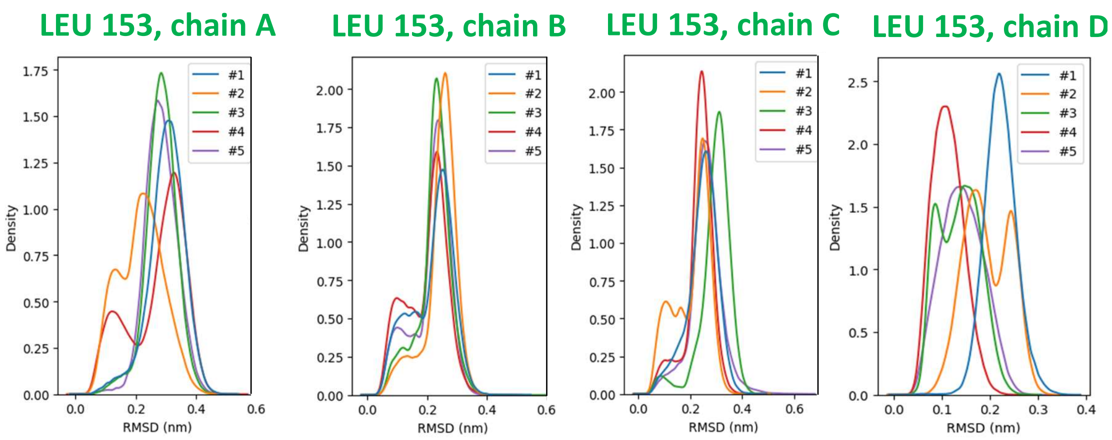


Figure S16: RMSD distributions of residues demonstrating bimodal distributions, continued. (Additional distributions presented in Figs S9-S15)

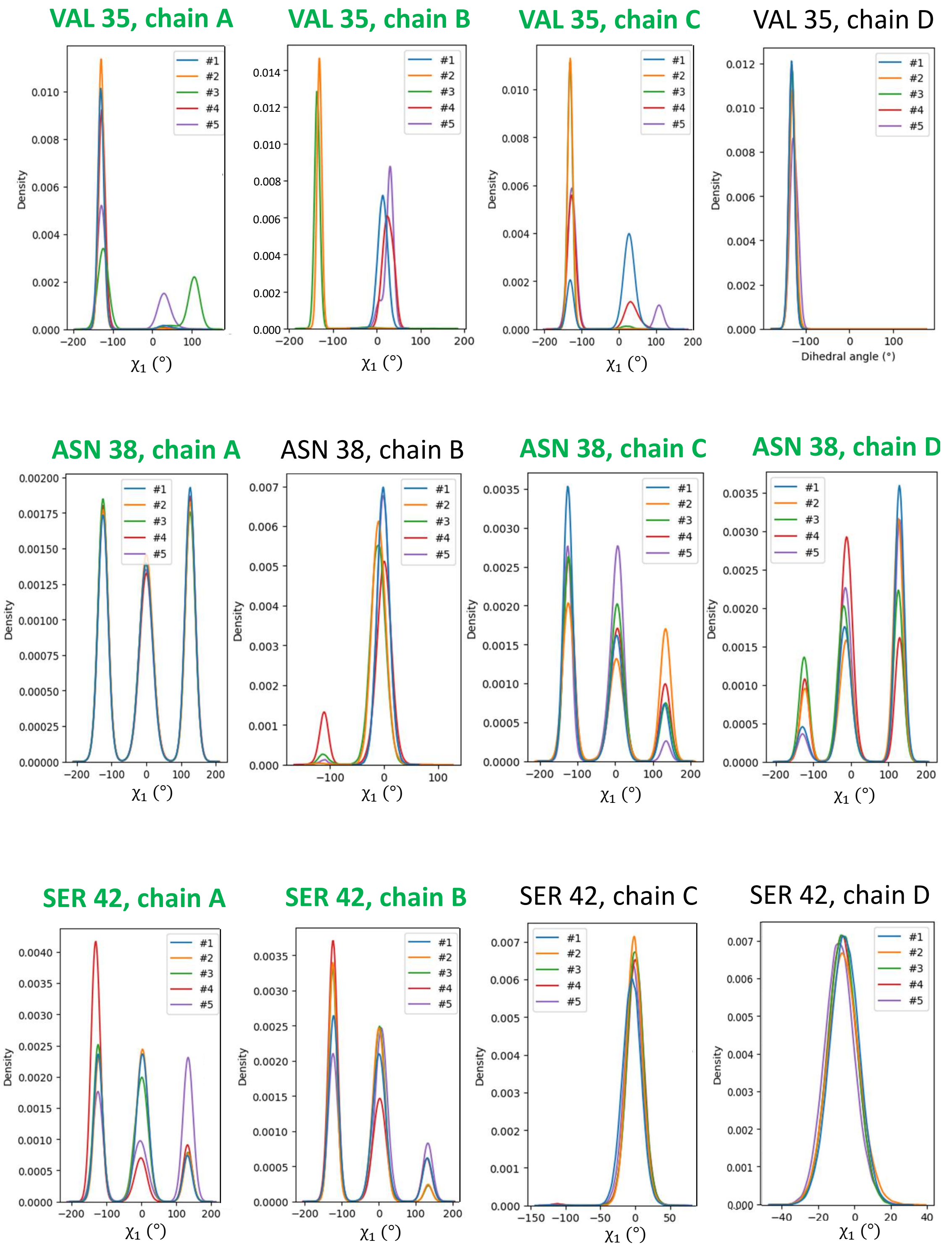


Figure S17: Dihedral angle distributions of residues demonstrating multimodal distributions. Green highlights demonstrate observed multimodal distributions used to calculate FELs. (Additional distributions presented in Figs S18-S24)

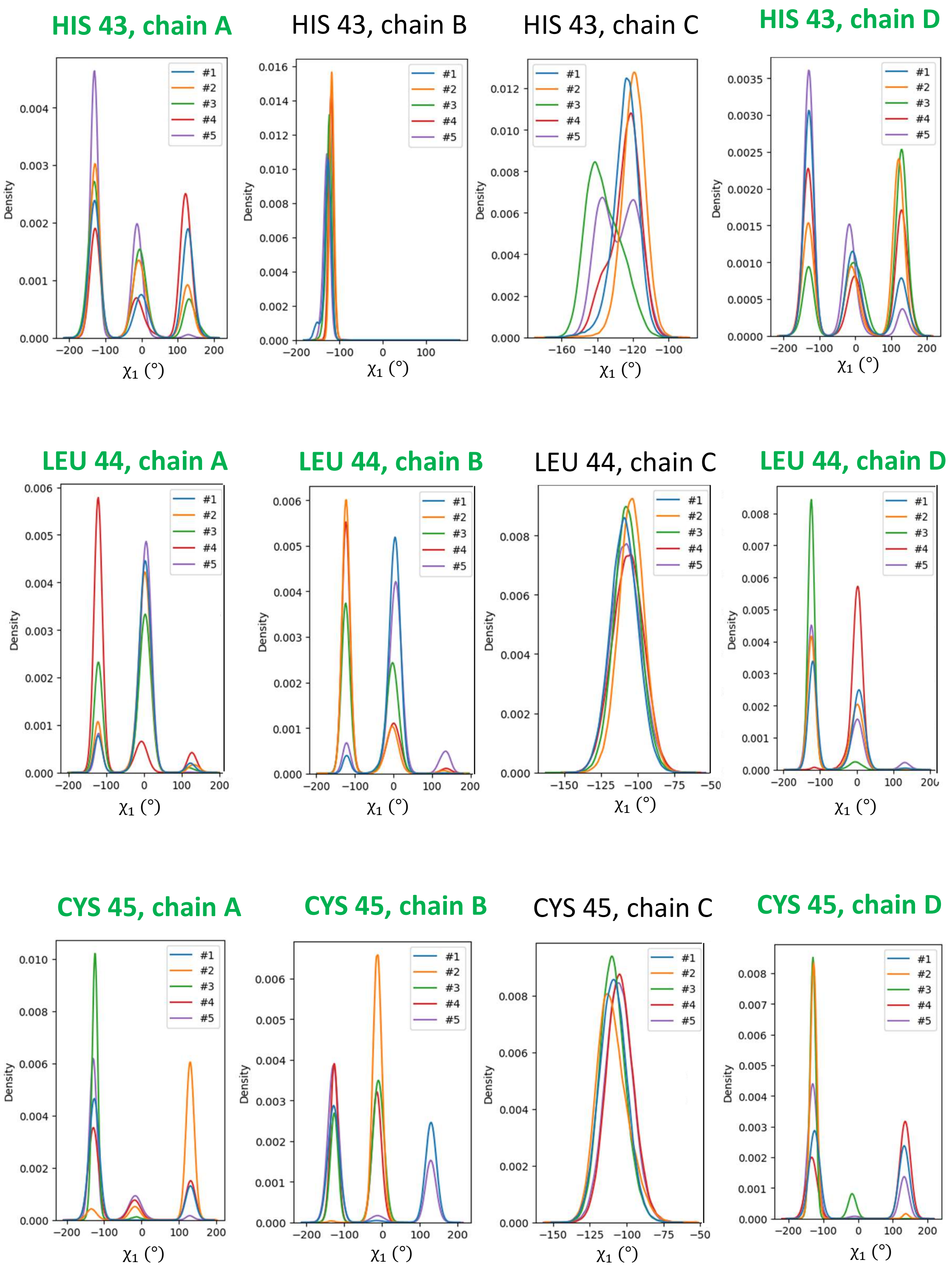


Figure S18: Dihedral angle distributions of residues demonstrating multimodal distributions, continued. (Additional distributions presented in Figs S17,S19-S24)

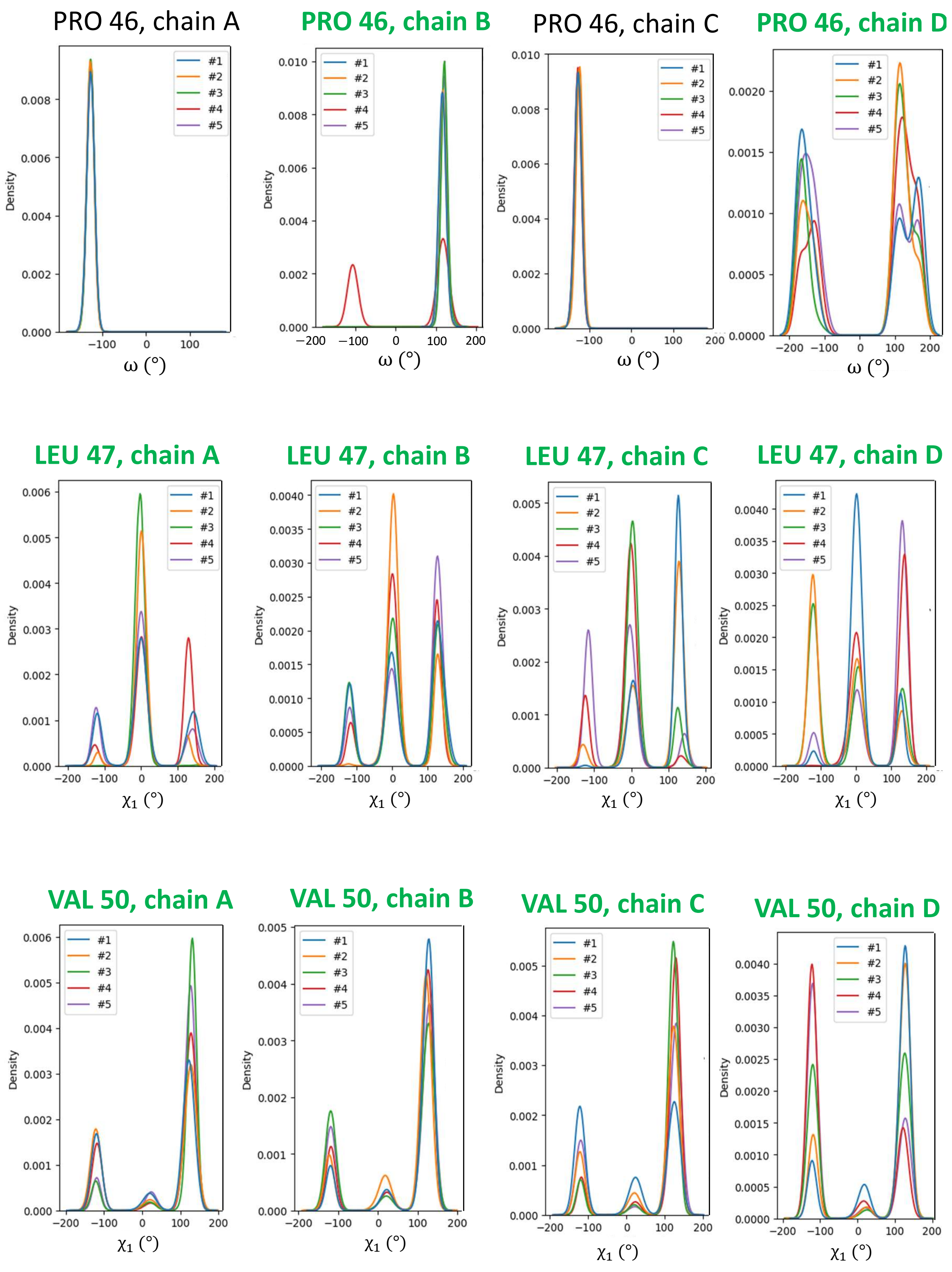


Figure S19: Dihedral angle distributions of residues demonstrating multimodal distributions, continued. (Additional distributions presented in Figs S17-S18,S20-S24)

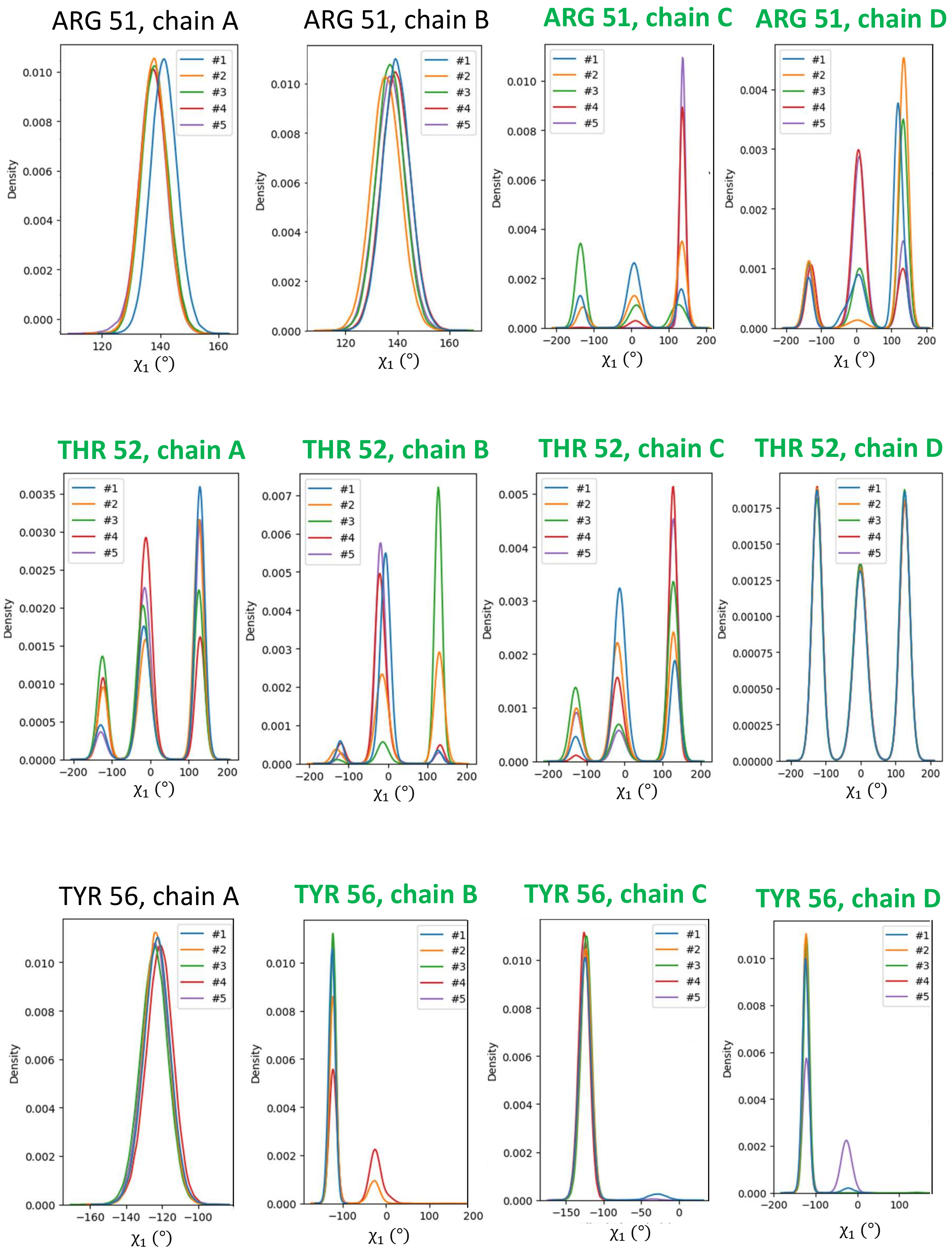


Figure S20: Dihedral angle distributions of residues demonstrating multimodal distributions, continued. (Additional distributions presented in Figs S17-S19,S21-S24)

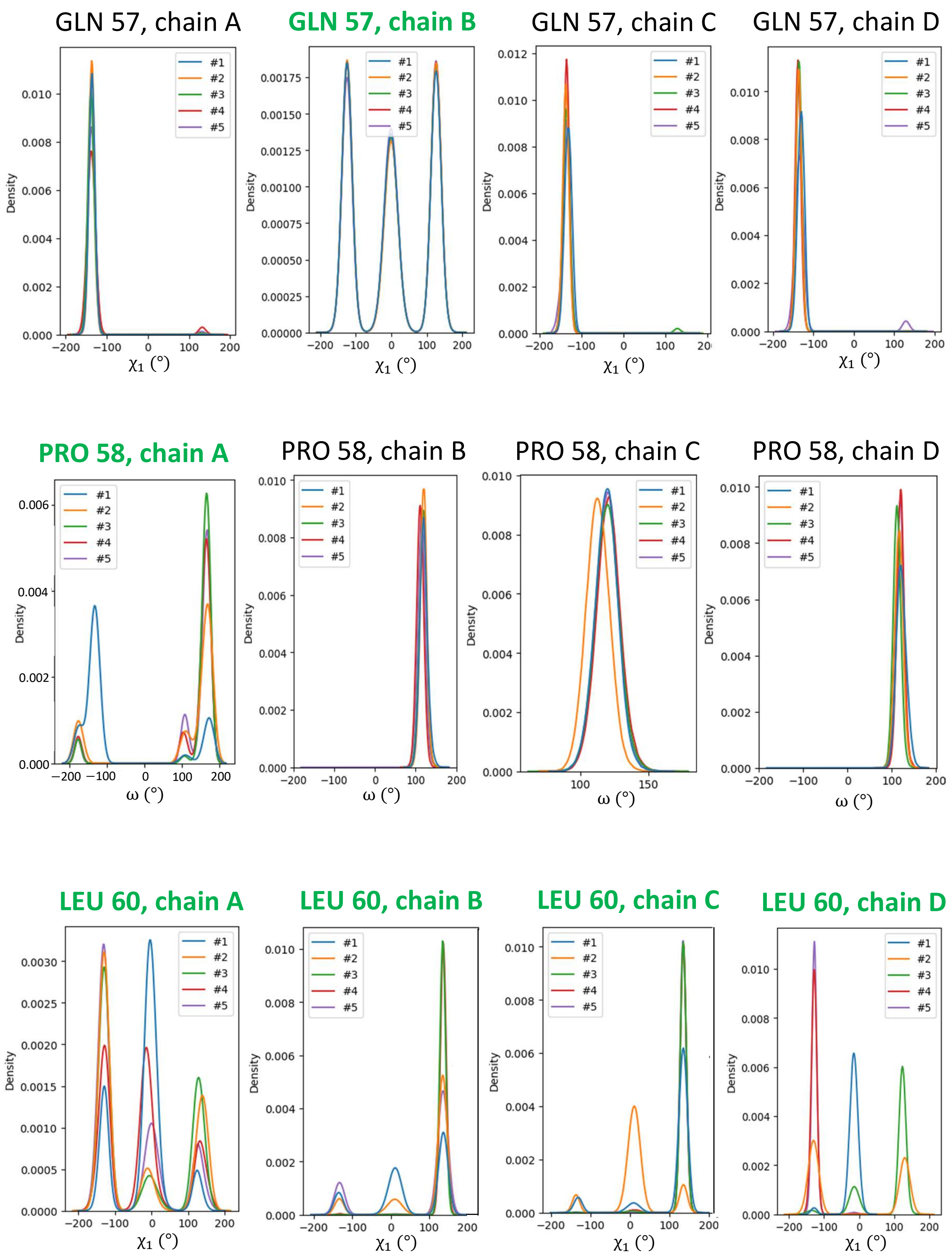


Figure S21: Dihedral angle distributions of residues demonstrating multimodal distributions, continued. (Additional distributions presented in Figs S17-S20,S22-S24)

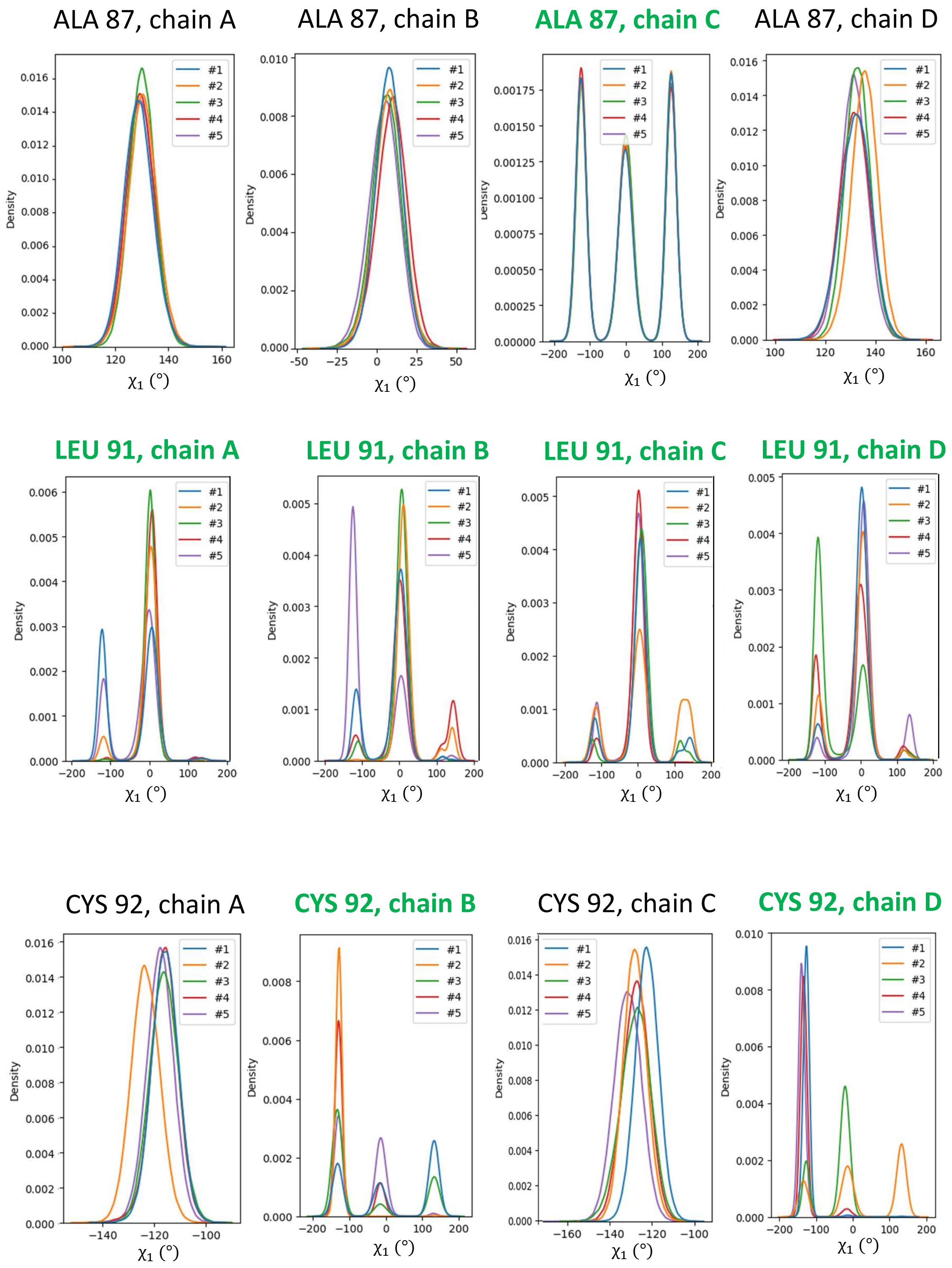


Figure S22: Dihedral angle distributions of residues demonstrating multimodal distributions, continued. (Additional distributions presented in Figs S17-S21,S23-S24)

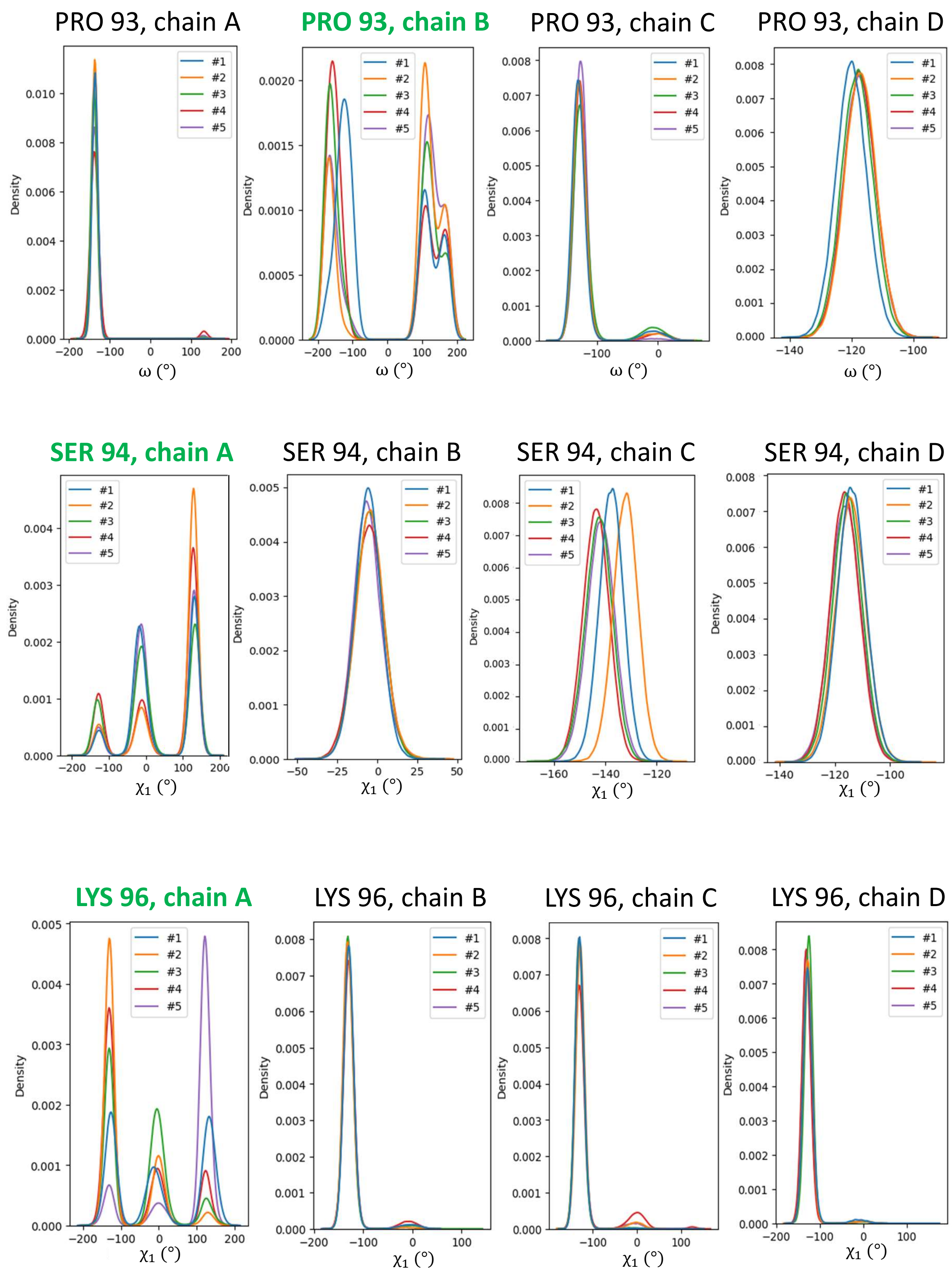


Figure S23: Dihedral angle distributions of residues demonstrating multimodal distributions, continued. (Additional distributions presented in Figs S17-S22,S24)

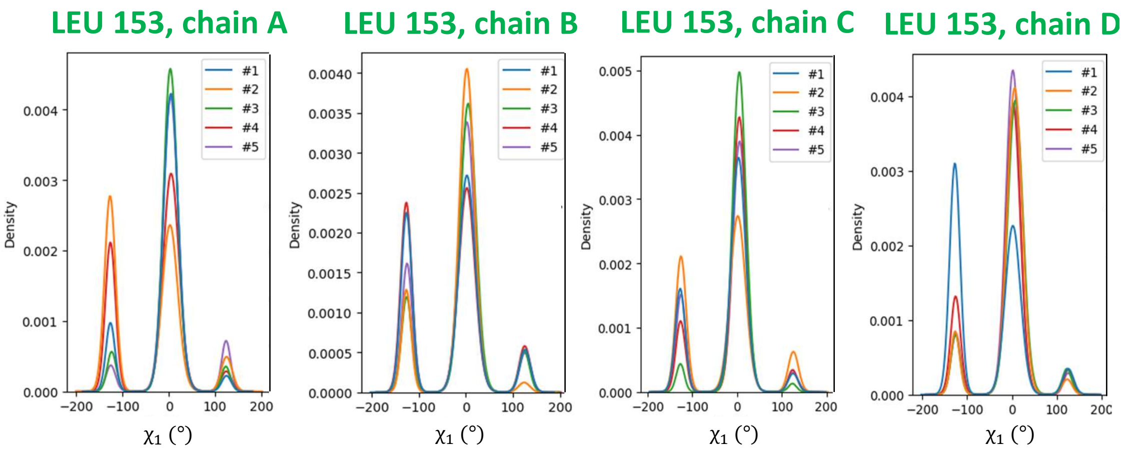


Figure S24: Dihedral angle distributions of residues demonstrating multimodal distributions, continued. (Additional distributions presented in Figs S17-S23)

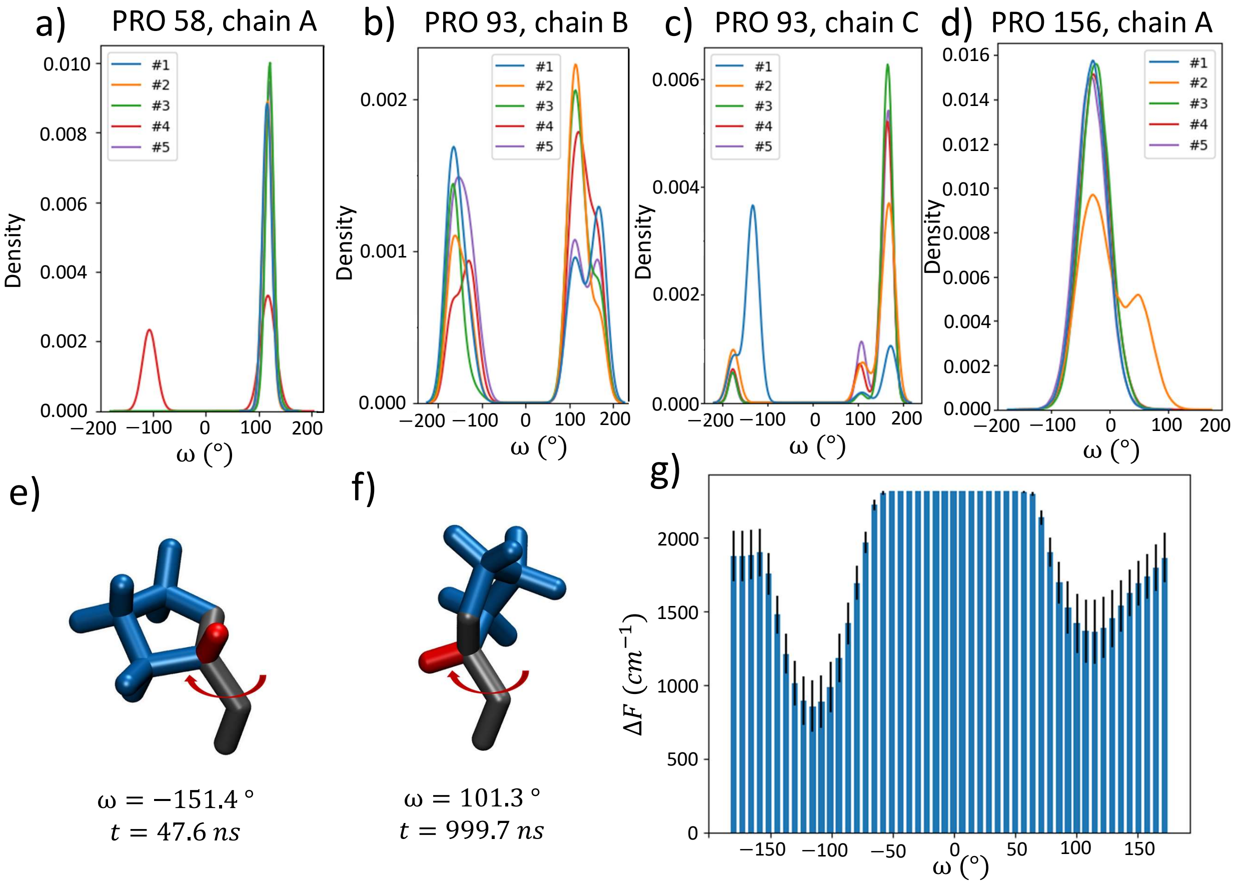


Figure S25: Bimodal distributions and FELs observed in rotation of PRO side chains. (a) Dihedral angle distributions of PRO 58, chain A, (b-c) PRO 93, in chains B and C, and (d) PRO 156, chain A. In e) and f) we demonstrate the two observed minimum free energy angular position, taken from PRO 93 in chain B, in replica #2 of the 300 K simulation at *e)* $t = 47.6\ ns$ with $\omega = -151.4°$, *f)* $t = 999.7\ ns$ with $\omega = 101.3°$. Heavy backbone atoms are depicted in black, the hydrogen backbone atom in red, the side chain pyrrolidine ring in blue. The rotation of the dihedral angle is represented by a red arrow. In g) we visualize the average free energy surface of PRO residues demonstrating bimodality.

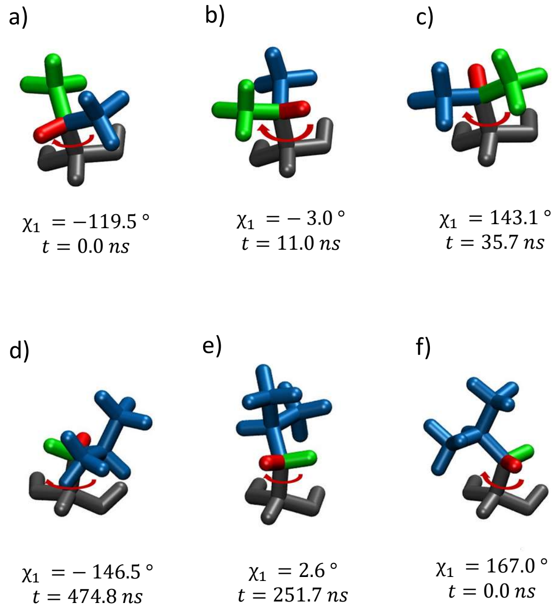


Figure S26: Visualization of primary positions of sidechain $\chi_1$ angles. (a-c) Graphical representations of VAL 50 in chain D, in replica #5 at *a)* $t = 0.0$ *ns* with $\chi_1 = -119.5°$, *b)* $t = 11.0$ *ns* with $\chi_1 = -3.0°$, and *c)* $t = 37.7$ *ns* with $\chi_1 = 143.1°$. (d-f) Graphical representations of LEU 47 in chain A, in replica #1 at *d)* $t = 0.0$ *ns* with $\chi_1 = 167.0°$, *e)* $t = 251.7$ *ns* with $\chi_1 = 2.6°$, and *f)* $t = 474.8$ *ns* with $\chi_1 = -146.5°$. Backbone atoms are depicted in black, the first side chain group in green, the second in red, and the last in blue. The rotation of the dihedral angle is represented by a red arrow.

#### S4.2 Bimodality in RMSD corresponds to trimodality in $\chi_1$ angles

A comparison of the RMSD distributions (Figs S9-S12) to the dihedral angle distributions (Figs S17-S20) for the non-PRO residues underscores that replicas displaying a double-peak in the RMSD distributions also exhibit a triple-peak pattern in the $\chi_1$ angle distributions. On the other hand, replicas with unimodal RMSD distributions demonstrated unimodal $\chi_1$ angle distributions. One example of this correlation can be seen in the case of TYR 56, in chain A, where all five replicas displayed unimodal RMSD distributions, and the $\chi_1$ angle distributions followed the exact same trend. This suggests that the variation in RMSD values can be attributed to the rotation of the dihedral angles. (For the PRO residues, we see bimodality in RMSD distributions corresponding to bimodality in $\omega_1$ distributions).

Note that while the RMSD distributions appear bimodal, the corresponding $\chi_1$ angle distributions nearly always exhibit three peaks, not two. To understand this discrepancy, it is important to consider the complex relationship between these two measures, which is influenced by various factors. The RMSD reflects the overall structural deviation of the protein, taking into account all residue atoms, while the $\chi_1$ angle focuses solely on the side chain conformation. As a result, the $\chi_1$ angle distribution provides more localized information about the conformational changes occurring in the side chain, which may not be fully captured by the RMSD. In particular, distinct $\chi_1$ angle values can result in conformations with minimal differences in the overall residue structure, resulting in RMSD values that do not show distinct peaks for these states . Instead, they might appear as a single peak due to their similarity in terms of flexibility. The norm position might also contribute to understanding the variation in peak numbers. If two $\chi_1$ values display minimal deviation from each other in terms of their atomic norm position, it indicates that the corresponding conformations may have negligible differences in the overall protein structure. Consequently, these subtle deviations might not be fully captured in the RMSD calculations, resulting in the observation of only two peaks in the RMSD distribution. However, considering the localized conformational changes reflected in the $\chi_1$ angle, these small differences become distinguishable, leading to the presence of three peaks in the $\chi_1$ angle distribution.

#### S4.3 Data supporting the relationship between bimodal RMSD distributions and multimodal sidechain dihedral distributions

The Root-mean-square fluctuation (RMSF) of the 168 residues were calculated to quantify the positional changes of individual atoms within each residue. This analysis aimed to gain a detailed understanding of residue dynamics by examining the behavior of individual atoms. The RMSF measures the average deviation of the position of each atom within a molecule throughout simulation, relative to a reference position. In Gromacs 2021.4, the RMSF of a particle $i$ is defined as[S3] :

$$RMSF_i = \left[ \frac{1}{T} \sum_{t_j=1}^{T} \parallel \mathbf{r}_i(t_j) - \mathbf{r}_i(t_{ref}) \parallel^2 \right]^{\frac{1}{2}} \tag{1}$$

where $T$ is the total simulation time, $\mathbf{r}_i(t_j)$ is the position of a particle $i$ at time $t_j$, and $\mathbf{r}_i(t_{ref})$ is the position of a particle $i$ at the reference time $t_{ref}$. The RMSF is averaged over time, giving a single value for each particle $i$. In our project, the reference position corresponds to the initial position of the residues at the start of the simulations. RMSF calculations were then averaged over the five replicas, at both temperatures. The RMSF analysis revealed that among the 168 residues, the same 55 residues identified in the RMSD analysis exhibited significant fluctuations in the positions of their side chains, while their backbone structure remained relatively stable. This finding confirms the hypothesis proposed in Section 3.3.1, suggesting that the bimodal RMSD distributions of these residues are primarily influenced by the movements of specific subsets of atoms (specifically the side chains), thus being at the origin of the conformational change. The subsequent paragraphs will explain these RMSF results, providing analysis of the observed fluctuations and their implications in the movements of the residues.

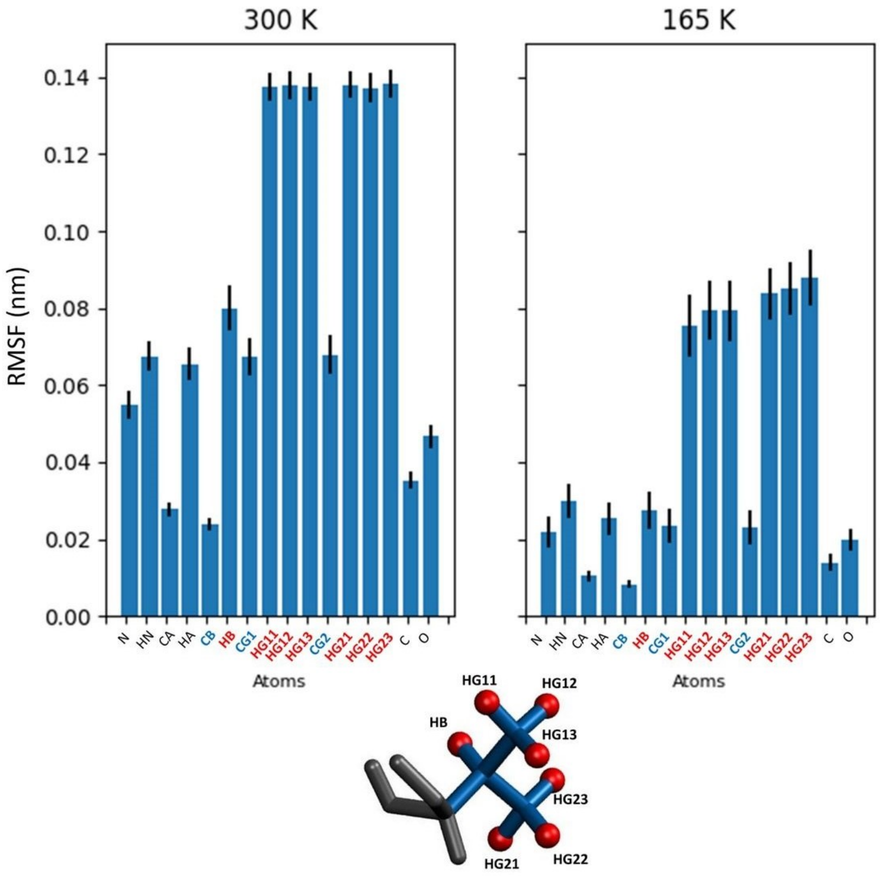


Figure S27: Average RMSF values of the 7 Valine residues (VAL 50 in all protein chains and VAL 35 in protein chains A, B, and C), at 300 K and 165 K. Backbone atoms are represented in black, side chain hydrogen atoms in red, and side chain non-hydrogen atoms in blue.

For readability, only results of the 55 residues are shown and the RMSF values were averaged by residue name. Figure S27 and Figure S28, depict the RMSF values of respectively the 7 VAL (VAL 50 in all protein chains and VAL 35 in protein chains A, B, and C) and the 19 LEU (LEU 47, LEU 60, LEU 91, LEU 153, in all protein chains, and LEU 44 in protein chains A, B and D), that display bimodal RMSD distributions. RMSF plots of the other less abundant residues can be found in Figure S30.

Hydrogen atoms exhibit the highest RMSF values, meaning they exhibit the highest change in position over the course of the simulations. This is expected as their smaller mass allows them to experience less inertia, making it easier for them to move and respond to

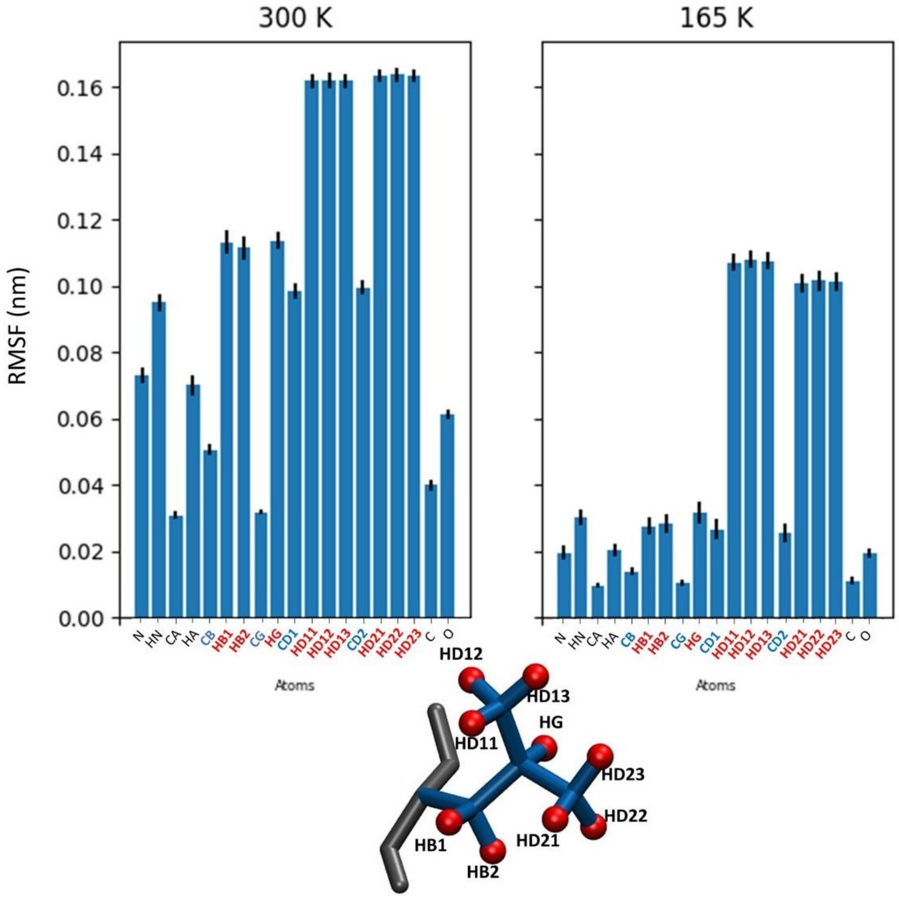


Figure S28: Average RMSF values of the 19 Leucine residues (LEU 47, LEU 60, LEU 91, LEU 153, in all protein chains, and LEU 44 in protein chains A, B and D), at 300 K and 165 K. Backbone atoms are represented in black, side chain hydrogen atoms in red, and side chain non-hydrogen atoms in blue.

external forces. Additionally, their implications in molecular vibrations, such as stretching and bending of chemical bonds, can also contribute to their motion. Nevertheless, the hydrogen atoms (H-atoms) of the side chains still have significantly higher RMSF values than H-atoms of backbone. It implies a greatest motion of the side chain H-atoms over the backbone H-atoms. Therefore, the side chain might be notably more flexible than the backbone. This assumption is supported by the fact, on average, non-heavy atoms in the side chains also exhibit higher RMSF values compared to non-heavy atoms in the backbone, sometimes even being equal or greater than the RMSF values of backbone H-atoms. This is the case for VAL: the backbone hydrogens HN and HA have the same RMSF values of almost 0.07 nm as the side chain carbons CG1 and CG2. This observation is also applicable to LEU: the backbone hydrogen HN has the same RMSF value of almost 0.10 nm as the side chain carbons CD1 and CD2. On the other hand, the other LEU backbone hydrogen HA has a significant lower RMSF value close to 0.07 nm. Consequently, the side chains seem to be the subsets of atoms responsible for the bimodal RMSD distributions, which consequently lead to the observed conformational change.

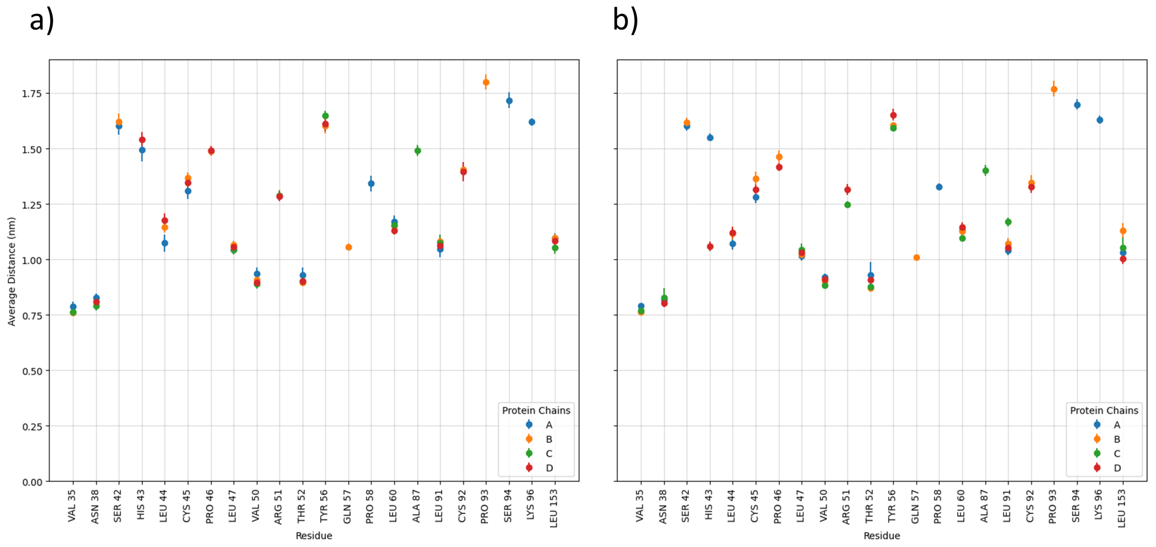


Figure S29: Distance between COM of Chlorophylls a and the COM of their respective residues at *a)* 300 K, and *b)* 165 K. Error bars represent standard errors taken over 5 independent replicas.

Distances between the center of mass (COM) of each residue and the COM of its corresponding Chl a were calculated. The results are presented in Figure S29 and were average over the five replicas, at both temperatures. The small standard errors suggest that the position of the residues remains relatively stable throughout the simulations. This finding further supports that the RMSD shift is likely not caused by a movement in the position of the residues, but rather a smaller conformational change involving specific subsets of atoms within the residues. Furthermore, the distances appear to be dispersed within the range of 0.75 nm to 1.80 nm. Therefore, it seems that the presence of a bimodal RMSD distribution is not influenced by the distance between the residue and its corresponding Chl a.

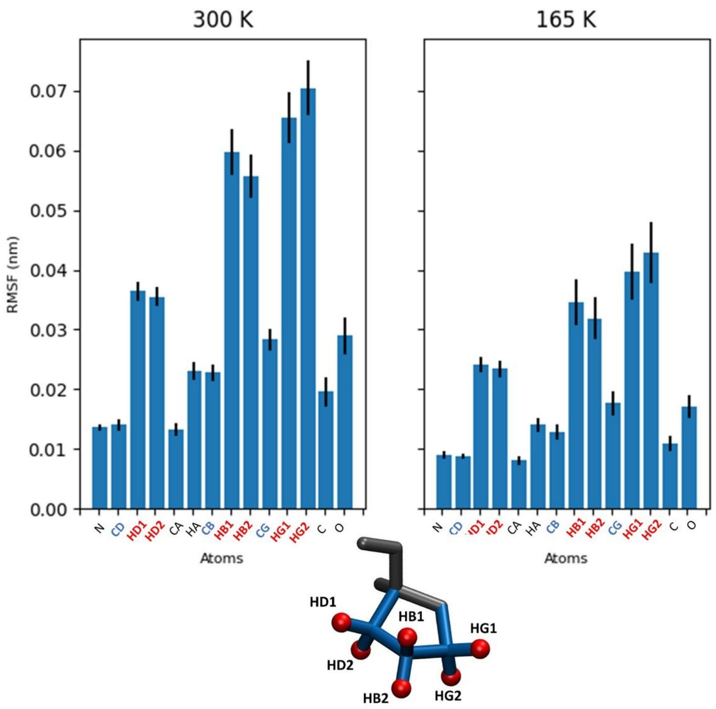


(a) Average RMSF values of the 4 Proline residues, at 300 K and 165 K.

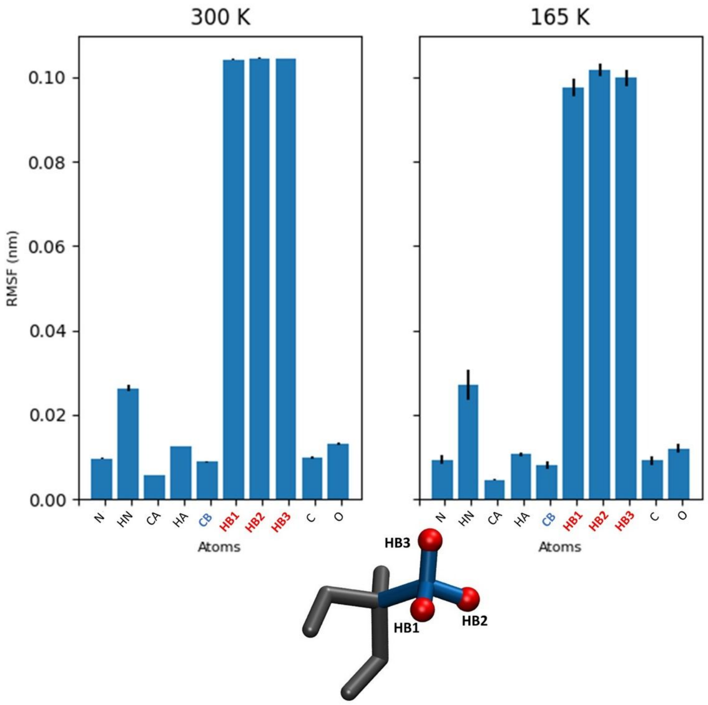


(b) Average RMSF values of the Alanine residue, at 300 K and 165 K.

#### S4.4 Motion correlations between close residues undergoing conformational change

This section focuses on examining the correlations in the motion of the 55 known residues that have been observed to undergo conformational changes, which are reflected in the rotation of their $\chi_1$ angle. The analysis employed a Dynamical Network approach. By analyzing the network, we aimed to identify correlations in the motion of these residues throughout the simulations. These correlations are represented as edges connecting the residues, with the thickness of each edge indicating its normalized weight. An edge with a thickness of 1 indicates that the residues move in a synchronized manner throughout all frames of the simulation. The primary goal was to determine if residues sharing similar properties, such as residue name, residue side chain type, or proximity in protein sequence, exhibited high correlations in their motion. Such correlations could suggest that conformational changes occur simultaneously and at comparable rates between these residues. The findings revealed high correlations (greater than 0.8) among residues that are adjacent in the protein sequence. Conversely, lower correlations (between 0.4 and 0.6) were observed for residues farther apart in the protein sequence. These results indicate that the specific residue name and side chain type do not play a substantial role in determining motion correlations. The most influential factors appear to be the proximity of residues in both the protein sequence and solution space[1]. These results may imply that conformational changes occur simultaneously and at a similar rate for closely located residues but non-simultaneously and at varying rates for farther residues. In other words, sub-groups of residues within the hydrophobic cavity around Chls a may undergo conformational changes independently, with their own distinct kinetics.

Figure S31 displays symmetrical heatmaps illustrating the weights of the edges and network representations for each protein chain at 300 K. The results presented in this study

[1] The solution space refers to the set of all possible conformations or structures that a protein can adopt.

were averaged over five independent replicas of the simulations. The standard errors of the means, which indicate the variability between replicas, were not displayed as they were too small to be visually discernible. The small magnitude of the standard errors, ranging from 0.001 to 0.010, suggests a high level of agreement between the independent runs of the simulations. This agreement reinforces the consistency of the observed correlations in motion among the residues.

For a given temperature, we can see slight divergence in residue correlations between the protein chains. For instance, in protein chain B, at 300 K, VAL 50 and LEU 153 exhibit an average edge weight of 0.4, while no significant correlation is observed between these residues at the same temperature in the other three protein chains. This slight discrepancy in correlations between the protein chains can be attributed to the relatively short simulation time. Despite these discrepancies, the overall trend in the results remains consistent across the protein chains. Residues located in close proximity to each other, either through direct covalent bonds or within a short distance of up to 5 residues, exhibit correlations greater than 0.8. As an example, this is the case of VAL 50 and THR 52, which display a weight edge greater than 0.8 in all protein chains and at both temperatures. These high correlations observed among nearby residues can be attributed to the covalent bonds, as residues that are directly bonded are typically part of the same secondary structure element, such as a $\alpha$-helix or $\beta$-strand. Covalent bonds restrict the relative motion between these residues, leading to a higher correlation in their motion. Therefore, residues such as LEU 91 and LEU 47, which are farther apart, exhibit a lower correlation of approximately 0.4 to 0.6 in each protein chain.

As a consequence these results, residues within the same community tend to be located in close proximity. In network analysis, a community is referred as group of residues more densely connected to each other than to the residues outside the community. In line with this concept, we can observe that residues within the same community, such as LEU 44, cysteine (CYS) 45, and PRO 46 in protein chain B, and LEU 91, CYS 92, and lysine (LYS) 96 in chain D, are located in close spatial proximity. This analysis might suggest that residues within the same community might display a rotation of their $\chi_1$ angle at the same rate.

Additionally, the results also indicate that residues with the same name or side chain type (non-polar, polar, acidic or basic) do not exhibit a specific correlation pattern. This can be exemplified by the case of LEU 60 and LEU 91, which are not correlated in any of the protein chains, despite having the same residue name, and therefore the same chemical composition. Similarly, the lack of correlation between VAL 35 and LEU 153, both non-polar residues, as well as between THR 52 and SER 94, both polar residues, supports this finding. These results suggest that the specific residue name or side chain type alone is not a determining factor for correlations in motion.

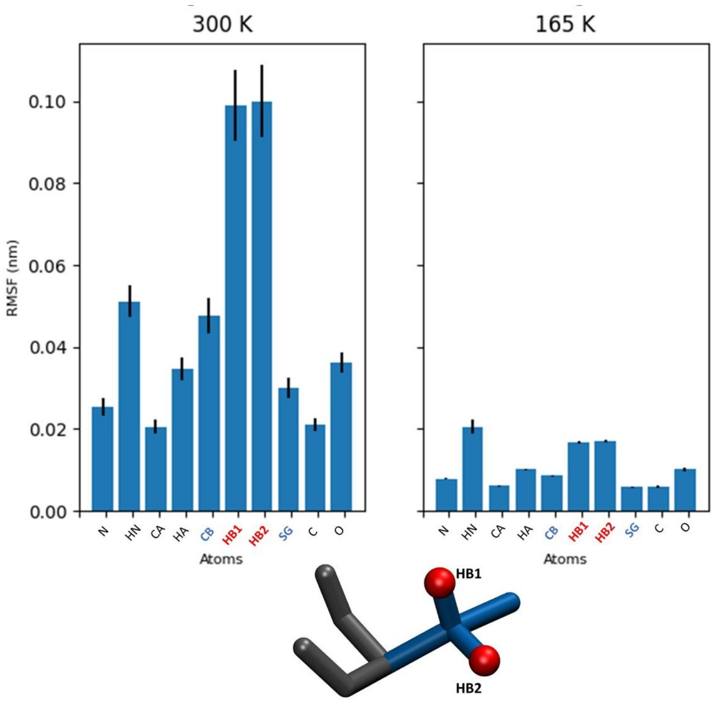


(c) Average RMSF values of the 5 Cysteine residues, at 300 K and 165 K.

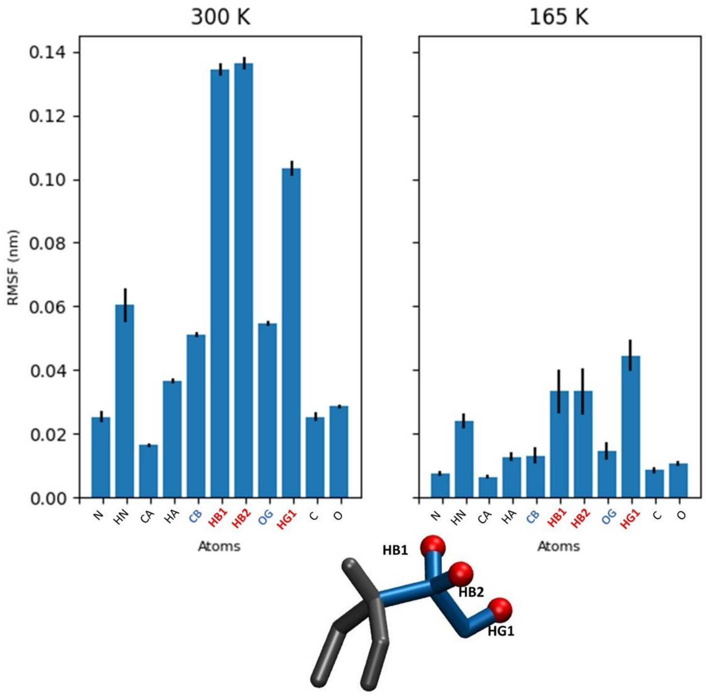


(d) Average RMSF values of the 3 Serine residues, at 300 K and 165 K.

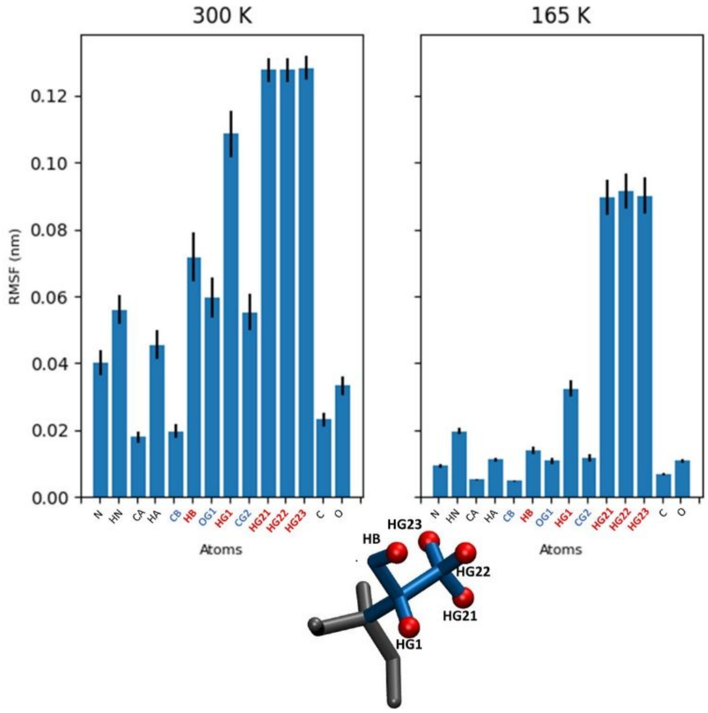


(e) Average RMSF values of the 4 Threonine residues, at 300 K and 165 K.

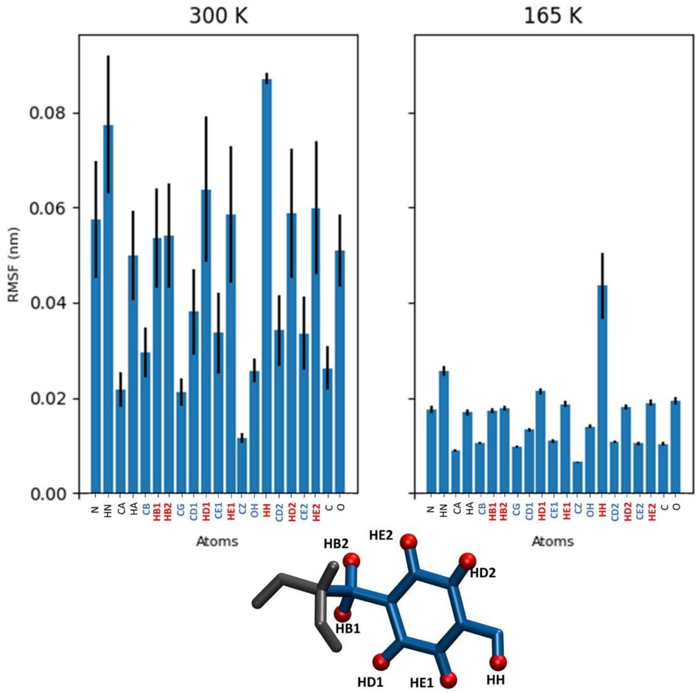


(f) Average RMSF values of the 3 Tyrosine residues, at 300 K and 165 K.

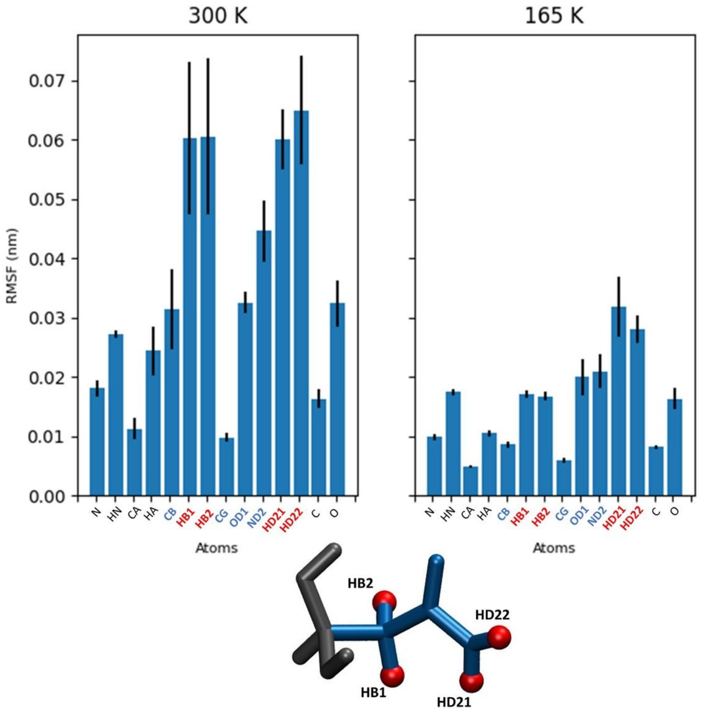


(g) Average RMSF values of the 3 Asparagine residues, at 300 K and 165 K.

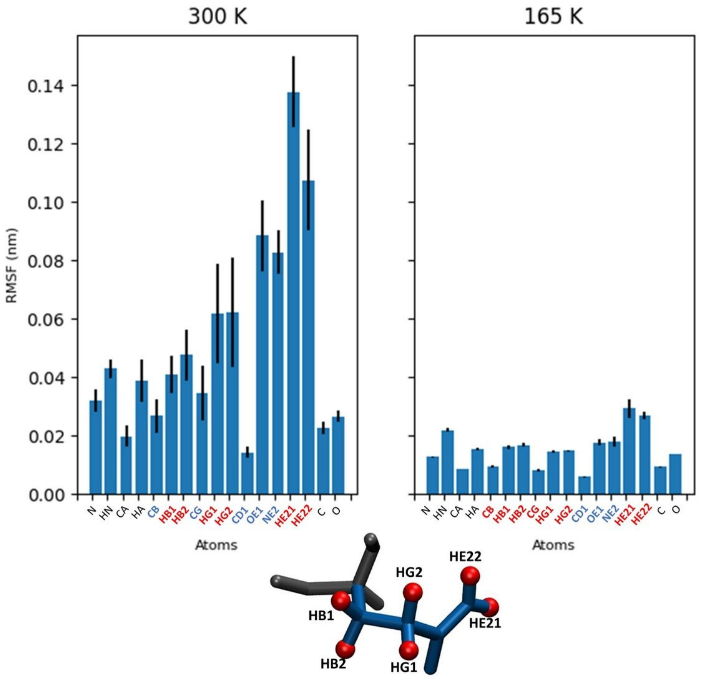


(h) Average RMSF values of the Glutamine residue, at 300 K and 165 K.

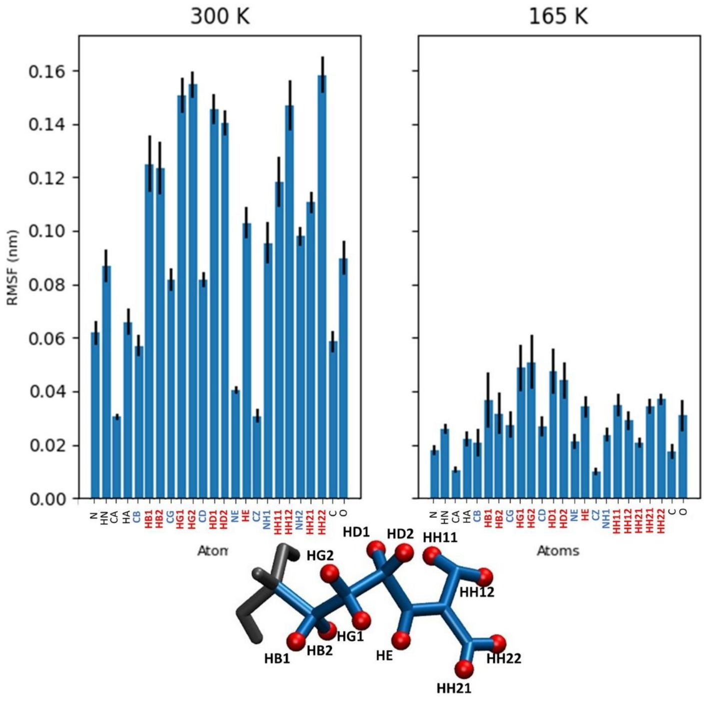


(i) Average RMSF values of the Arginine residues, at 300 K and 165 K.

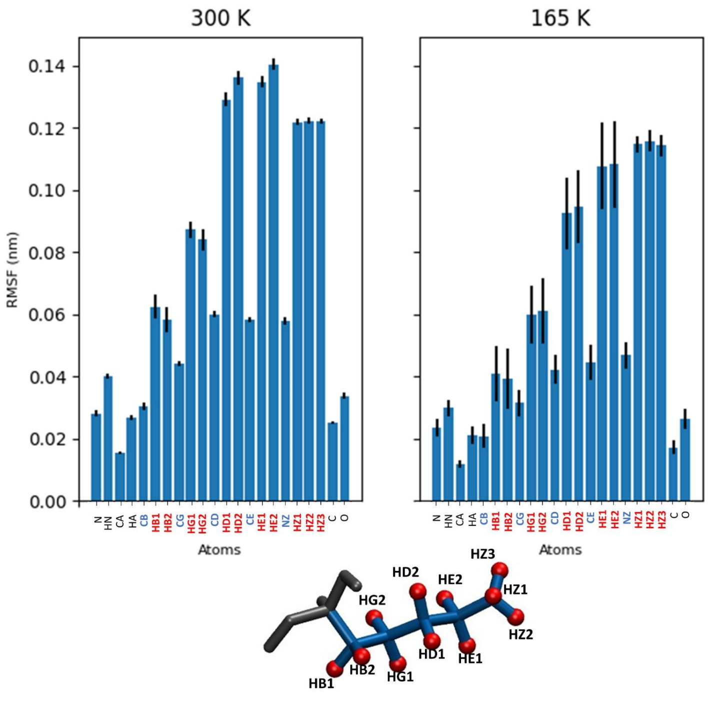


(j) Average RMSF values of the Lysine residue, at 300 K and 165 K.

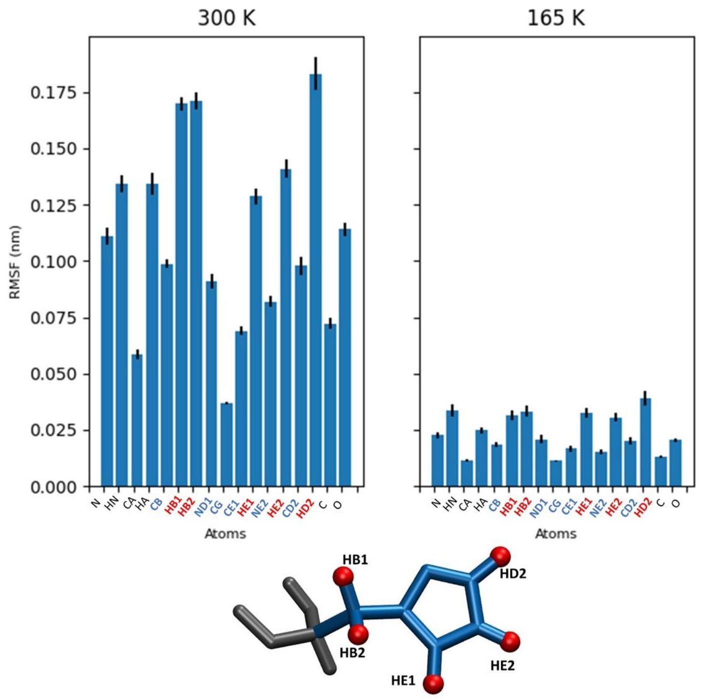


(k) Average RMSF values of the 2 Histidine residues, at 300 K and 165 K.

Figure S30: Average RMSF values classified by residue types, at 300 K and 165 K. Backbone atoms are labeled and represented in black, side chain hydrogen atoms in red, and side chain non-hydrogen atoms in blue.

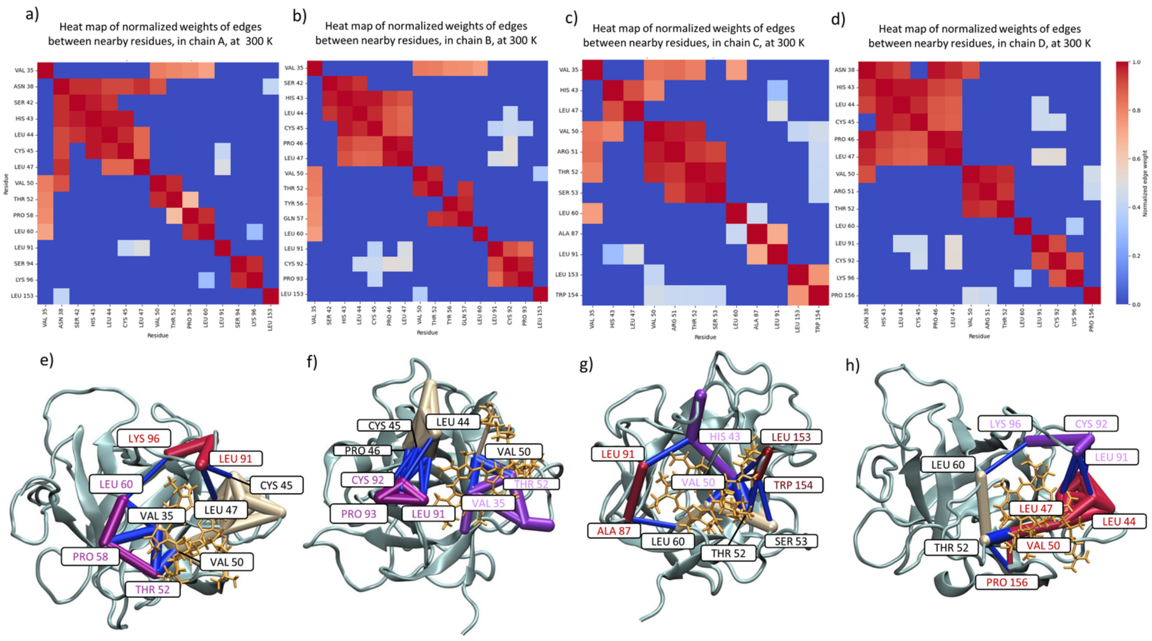


Figure S31: a), b), c), and d), represent the heat maps of the normalized weights of edges between residues near Chls a that undergo conformational change, in respectively, protein chains A, B, C, and D, at 300 K. e), f), g), and h) illustrate their respective networks, and residues involved in inter-communities interactions are labeled. Communities are colored in beige, pink, magenta, burgundy and purple. Inter-community links are colored in navy blue. The Chl a is colored in orange and the protein sub-unit in blue. The thicknesses of the edges correspond to the normalized weights of these edges.

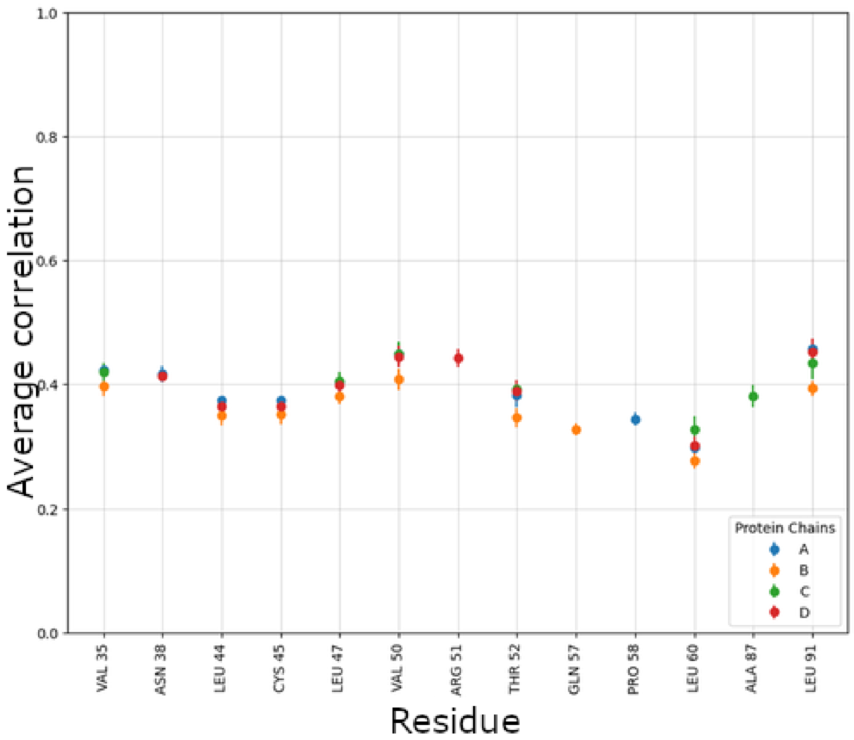


Figure S32: Normalized weights of edges (average correlation) between THR 180 and the nearby residues. Error bars represent standard errors taken over 5 independent replicas.

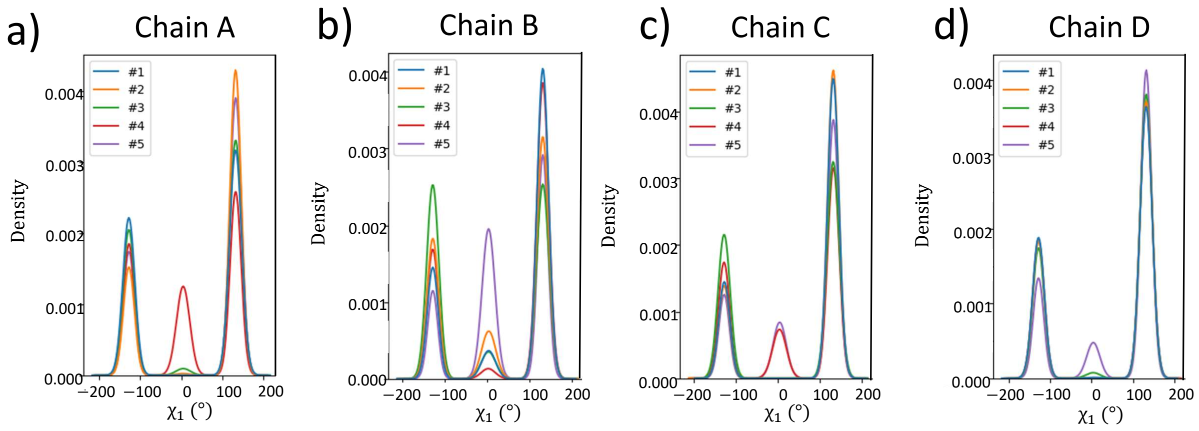


Figure S33: Dihedral angle ($\chi_1$ ) distributions of THR 180 in different chains.

### References

(S1) Swenson, David W. H. and Roet, Sander Contact Map Explorer. https://contact-map.readthedocs.io/en/latest/index.html.

(S2) Humphrey, W.; Dalke, A.; Schulten, K. VMD: visual molecular dynamics. *J. Mol. Graphics* **1996**, *14*, 33–38.

(S3) Abraham, M. J.; Murtola, T.; Schulz, R.; Páll, S.; Smith, J. C.; Hess, B.; Lindahl, E. GROMACS: High performance molecular simulations through multi-level parallelism from laptops to supercomputers. *SoftwareX* **2015**, *1*, 19–25.
